# Engineering *Yarrowia lipolytica* as a chassis for *de novo* synthesis of five aromatic-derived natural products and chemicals

**DOI:** 10.1101/2020.04.04.025288

**Authors:** Yang Gu, Jingbo Ma, Yonglian Zhu, Xinyu Ding, Peng Xu

## Abstract

*Yarrowia lipolytica* is a novel microbial chassis to upgrade renewable low-cost carbon feedstocks to high-value commodity chemicals and natural products. In this work, we systematically characterized and removed the rate-limiting steps of the shikimate pathway and achieved *de novo* synthesis of five aromatic chemicals in *Y. lipolytica*. We determined that eliminating amino acids formation and engineering feedback-insensitive DAHP synthases are critical steps to mitigate precursor competition and relieve the feedback regulation of shikimate pathway. Further overexpression of heterologous phosphoketolase and deletion of pyruvate kinase provided a sustained metabolic driving force that channels E4P (erythrose 4-phosphate) and PEP (phosphoenolpyruvate) precursors through the shikimate pathway. Precursor competing pathways and byproduct formation pathways were also blocked by inactivating chromosomal genes. To demonstrate the utility of our engineered chassis strain, three natural products, 2-phenylethanol (2-PE), *p*-coumaric acid and violacein, which were derived from phenylalanine, tyrosine and tryptophan, respectively, were chosen to test the chassis performance. We obtained 2426.22 ± 48.33 mg/L of 2-PE, 593.53 ± 28.75 mg/L of *p*-coumaric acid, 12.67 ± 2.23 mg/L of resveratrol, 366.30 ± 28.99 mg/L of violacein and 55.12 ± 2.81 mg/L of deoxyviolacein from glucose in shake flask. The 2-PE production represents a 286-fold increase over the initial strain (8.48 ± 0.50 mg/L). Specifically, we obtained the highest 2-PE, violacein and deoxyviolacein titer ever reported from the *de novo* shikimate pathway in yeast. These results set up a new stage of engineering *Y. lipolytica* as a sustainable biorefinery chassis strain for *de novo* synthesis of aromatic compounds with economic values.

---

*Y. lipolytica*, as a ‘generally regarded as safe’ (GRAS) yeast ^1^, has been extensively engineered for the production of oleochemicals, fuels and commodity chemicals ^2-5^. The abundant acetyl-CoA and malonyl-CoA precursors in *Y. lipolytica* have been harnessed for synthesizing plant secondary metabolites in recent years, including flavonoids ^6-9^, polyketides ^10-11^, polyunsaturated fatty acids ^12-13^ and isoprenoids ^14-17^. A large collection of customized genetic toolkits, including Golden-gate cloning ^18-20^, genome integration ^6, 21-22^, CRISPR-Cas9/Cpf1 genome editing ^23-25^, transposons ^26^, auxotrophic markers ^27^ and promoter libraries ^28-30^ have accelerated our ability to perform targeted genetic manipulations. Different from *S. cerevisiae, Y. lipolytica* lacks Crabtree effects generating no overflowed metabolism toward ethanol under high glucose conditions ^31^, which might be more suitable for high gravity fermentation and process control. *Y. lipolytica* is reported to valorize a broad range of cheap and renewable feedstocks ^32-33^, including sugars, volatile fatty acids, alkanes and municipal organic wastes. This substrate flexibility provides us an environmentally friendly approach to upgrade low-value carbons to high value chemicals with reduced carbon footprint and improved process economics.

Aromatic compounds have wide applications ranging from health care to food industry, nutraceutical supplements and pharmaceutical intermediates, which collectively represent a multi-billion-dollar global market ^34-37^. Although metabolically-engineered *E. coli* has achieved gram-per-liter levels of aromatic compounds, primarily phenylpropanoids ^38-39^, yeast proves to be a more attractive host, due to its GRAS status, robust cell growth, tolerance of harsh conditions (such as low pH and high osmolarity), and the spatially-organized subcellular compartment for regio- or stereo-activity of cytochrome P450 enzymes ^7, 34, 37, 40^. Up to date, various metabolic engineering strategies have been implemented in *E. coli* and *S. cerevisiae* to improve aromatics production, including expression of the feedback-insensitive DAHP synthases ^41^ and chorismite synthase, enhancing the shikimate pathway ^42^ and modular microbial coculture ^43-45^. A number of studies have been focused on optimizing the endogenous Ehrlich pathway to produce 2-phenylethanol up to 3-6 g/L ^46-48^, however, the expensive precursor L-phenylalanine was fed into the bi-phasic bioreactors ^49-50^, restricting its economic potential for large-scale production. The highest titer of 2-phenylethanol production from the *de novo* shikimate pathway was reported recently as 1.58 g/L (13 mM) from *S. cerevisiae* ^41^. There is a pressing need to explore the metabolic potential of alternative yeast for aromatics production.

In this work, we attempted to engineer the nonconventional oleaginous yeast (*Y. lipolytica*) as a competitive platform host to produce aromatic derivatives (Figure 1). With 2-phenylethanol (2-PE) as the testbed molecule, we systematically characterized and removed the bottlenecks of the endogenous shikimate pathway, resulting in the production of 2426.22 ± 48.33 mg/L (19.85 ± 0.40 mM) of 2-PE from glucose, the highest titer ever reported from the *de novo* shikimate pathway. Using this yeast as chassis, we further redirected the shikimate flux toward other aromatics and achieved high tier of *p*-coumaric acid (593.53 ± 28.75 mg/L), violacein (366.30 ± 28.99 mg/L, highest titer reported in yeast) and deoxyviolacein (55.12 ± 2.81 mg/L, highest titer reported in yeast) in shaking flasks, indicating the superior metabolic potential of *Y. lipolytica* as an aromatics-producing host. This report highlights the prominent metabolic characteristics of *Y. lipolytica* as chassis for production of various aromatics and natural products with economic values.

**Figure 1.**
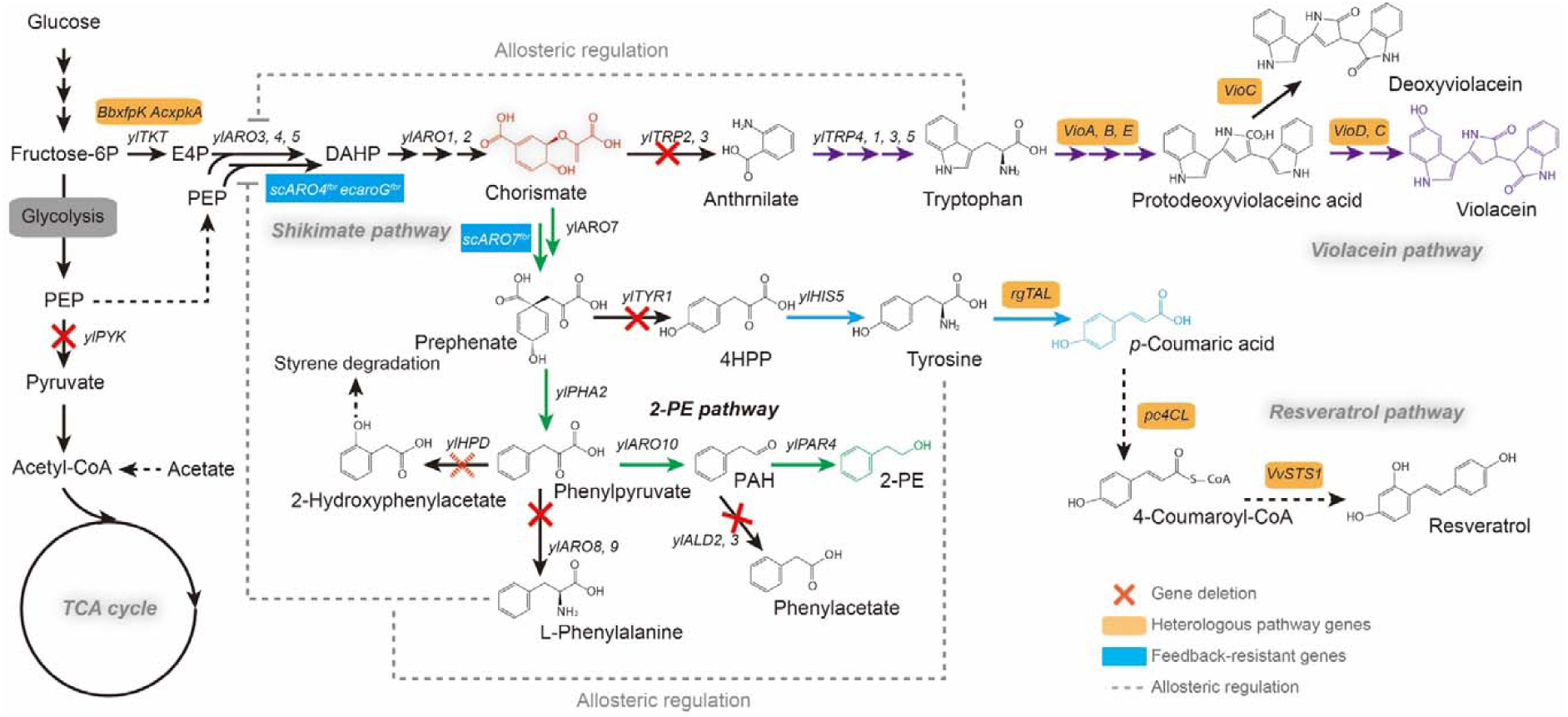
De novo biosynthetic routes for 2-PE, *p*-coumaric acid, resveratrol and violacein production through the shikimate pathway in *Y. lipolytica*. PEP, phosphoenolpyruvate; E4P, erythrose-4P; DAHP, 3-deoxy-arabino-heptulonate-7-phosphate; 4HPP, 4-hydroxyphenylpyruvate; PAH, phenylacetaldehyde; 2-PE, 2-phenylethanol; ylTKT, transketolase; ylPYK, pyruvate kinase; ylARO3, 3-deoxy-7-phosphoheptulonate synthase; ylARO4, 3-deoxy-7-phosphoheptulonate synthase; ylARO5, 3-deoxy-7-phosphoheptulonate synthase; ylARO1, pentafunctional AROM polypeptide; ylARO2, chorismate synthase; ylARO7, chorismate mutase; ylTRP2, anthranilate synthase; ylTRP3, anthranilate synthase; ylTRP4, anthranilate phosphoribosyltransferase; ylTRP1, phosphoribosylanthranilate isomerase; ylTRP5, tryptophan synthase; ylTYP1, prephenate dehydrogenase; ylARO8, aromatic amino acid aminotransferase; ylARO9, aromatic amino acid aminotransferase; ylPHA2, prephenate dehydratase; ylARO10, phenylpyruvate decarboxylase; ylPAR4, phenylacetaldehyde reductases; ylHPD, 4-hydroxyphenylpyruvate dioxygenase; ylALD2, aldehyde dehydrogenase; ylALD3, aldehyde dehydrogenase; scARO7^fbr^, the feedback-resistant chorismate mutase from S. cerevisiae; BbxfpK, phosphoketolase from *Bifidobacterium breve*; AcxpkA, phosphoketolase from *Acidobacterium capsulatum*; ecaroG^fbr^, the feedback-resistant 3-deoxy-7-phosphoheptulonate synthase from *E. coli*; scARO4^fbr^, the feedback-resistant 3-deoxy-7-phosphoheptulonate synthase from *S. cerevisiae*; rgTAL, tyrosineammonia-lyase from *Rhodotorula toruloides*; pc4CL, 4-coumarate-CoA ligase from *Petroselinum crispum*; VvSTS1, resveratrol synthase from *Vitis vinifera*; VioA, tryptophan oxidase; VioB/VioE, protodeoxyviolaceinate synthase; VioD, protodeoxyviolaceinate monooxygenase; VioC, violacein synthase. All violacein pathway genes were amplified from the genomic DNA of *Chromobacterium violaceum*. Right corner shows the chromosomal integration of the chosen pathways.

## Results and discussions

### *De novo* synthesis of 2-PE from shikimate pathway

In this work, the *Y. lipolytica* endogenous metabolite 2-phenylethnol (2-PE, an aromatic alcohol that is widely used as flavor and fragrance agent) was chosen as a target molecule to determine the bottlenecks of the shikimate pathway. 2-PE synthesis is a metabolic branch derived from shikimate pathway, in which 2-PE is synthesized through four enzyme-dependent cascade reactions starting from chorismate (Figure 2a). To identify the potential limiting-steps in 2-PE synthesis, we firstly systematically overexpressed all genes that are involved in 2-PE synthesis under the control of strong constitutive pTEF-intron promoter ^30^, including genes *ylPAR4* (*YALI0D07062g*, encoding phenylacetaldehyde reductase), *ylARO10* (*YALI0D06930g*, encoding phenylpyruvate decarboxylase), *ylPHA2* (*YALI0B17336g*, encoding prephenate dehydratase) and *ylARO7* (*YALI0E17479g*, encoding chorismate mutase). Besides gene *ylARO10*, individual overexpression of genes *ylPAR4, ylPHA2*, and *ylARO7* did not result in a significant increase in 2-PE titer (Supplementary Figure S2), however, combining all four genes (strain **YL5**) led the strain to produce 55.53 ± 0.85 mg/L of 2-PE from glucose (Supplementary Figure S2), a 6.55-fold increase in comparison with the control strain (without any gene overexpression, 8.48 ± 0.50 mg/L), indicating that the four enzymes in the Ehrlich pathway are essential to channel the carbon flux toward 2-PE synthesis. In addition, chorismate mutase (ARO7) is known to be strictly feedback inhibited by aromatic acids ^42^. To resolve this issue, introduction of a feedback-resistant chorismate mutase ScARO7^G141S^ originating from *Saccharomyces cerevisiae* further improved 2-PE production up to 87.63 ± 4.67 mg/L (strain **YL6**, Figure 2b), which was consistent with the previous report ^51^. We have configured the Ehrlich pathway to achieve *de novo* 2-PE synthesis from glucose with enhanced titer.

**Figure 2.**
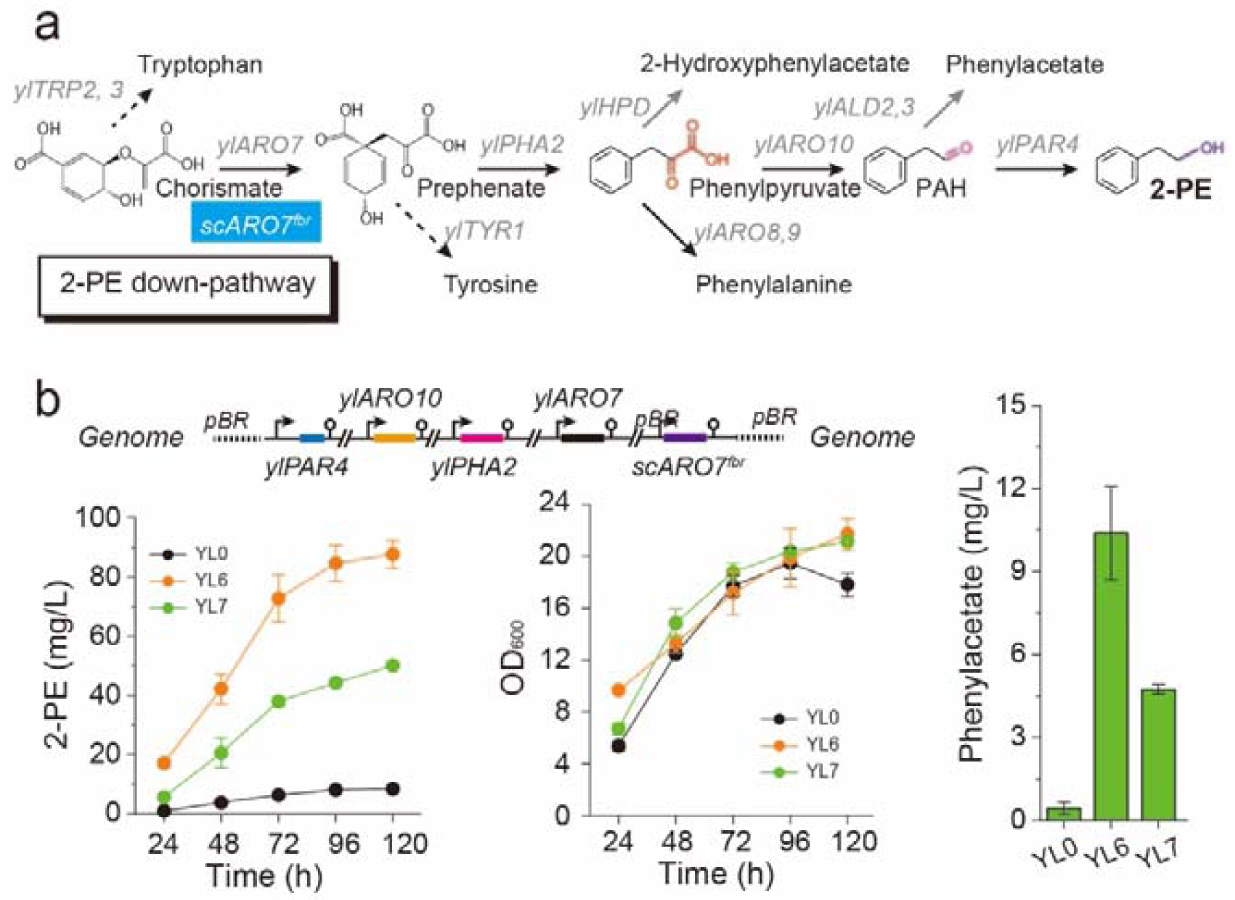
Validating and relieving bottlenecks in the Ehrlich pathway for 2-PE synthesis. (a) Enzyme cascade reactions of 2-PE production from chorismate; (b) Time profiles of 2-PE titer, cell growth, and phenylacetate titer of strains carrying the 2-PE pathway. All experiments were performed in triplicate and error bars represent standard deviations (SD).

We subsequently optimized the shikimate pathway to unlock the potential of *Y. lipolytica* for production of aromatic compounds. The entire Ehrlich pathway for 2-PE synthesis contains multiple genes with total size more than 16,000 bp ^52^. To simplify the genetic manipulations, we integrated the 2-PE pathway at the genomic *YALI0E30965g* loci (encoding acetyl-CoA hydrolase) with the integration plasmid *pURLA-ylPAR4-ylARO10-ylARO7-ylPHA2-scARO7* ^*G141S*^. The resultant strain **YL7** produced 50.13 ± 0.62 mg/L of 2-PE from glucose (Figure 2b). In *Y. lipolytica*, shikimate pathway is consisted of seven enzymatic steps, which starts from precursors erythrose-4-phosphate (E4P) and phosphoenolpyruvate (PEP) and ends up with chorismate (Figure 1). By overexpressing genes ylARO1 (YALI0F12639g, encoding pentafunctional aromatic protein), *ylARO2* (*YALI0D17930g*, encoding bifunctional chorismate synthase), *ylARO3* (*YALI0B20020g*, encoding DAHP synthase), *ylARO4* (*YALI0B22440g*, encoding DAHP synthase), and *ylARO5* (*YALI0C06952g*, encoding DAHP synthase) in strain **YL7**, we generated strains **YL8, YL9, YL10, YL11**, and **YL12** that were able to produce 2-PE with titers at 62.53 ± 1.28, 52.80 ± 2.18, 59.49 ± 1.70, 54.71 ± 0.87, and 57.16 ± 2.64 mg/L (Supplementary Figure S3), respectively. Strains **YL8, YL10**, and **YL12** showed slight increase of 2-PE titer compared with strain **YL7**. However, by simultaneous overexpression of genes *ylARO1* and *ylARO2*, 2-PE production was increased to 84.64 ± 4.42 mg/L (strain **YL13**, Supplementary Figure S3). Further overexpression of *ylARO3, ylARO4*, and *ylARO5* did not have significant effect on 2-PE titer (strain **YL14**, 78.28 mg/L, Supplementary Figure S3), which is possibly due to the allosteric regulation of DAHP synthase by aromatic amino acids, a common phenomenon in *S. cerevisiae* ^34, 42^.

### Relieving metabolic bottlenecks in the shikimate pathway

Heterologous DAHP synthases scARO4^K229L^ (originating from *S. cerevisiae*), AroG^L175D^ and AroG^S180F^ (originating from *E. coli*) previously have been shown to be feedback-resistant of aromatic amino acids regulation ^42^. We next introduced three variants of the feedback-resistant DAHP synthases into the 2-PE chassis strain, and generated strains **YL15, YL16**, and **YL17**. Shake flask cultivation of strains **YL15, YL16**, and **YL17** led to the production of 313.13 ± 10.16, 48.70 ± 5.92, and 430.11 ± 33.05 mg/L of 2-PE (Figure 3a, b), respectively. This result clearly demonstrates that overexpression of scARO4^K229L^ and AroG^S180F^ are effective to remove the shikimate pathway bottleneck in *Y. lipolytica*. Likewise, the results indicated the importance of relieving allosteric regulation of DAHP synthase to increase carbon flux toward shikimate pathway.

**Figure 3.**
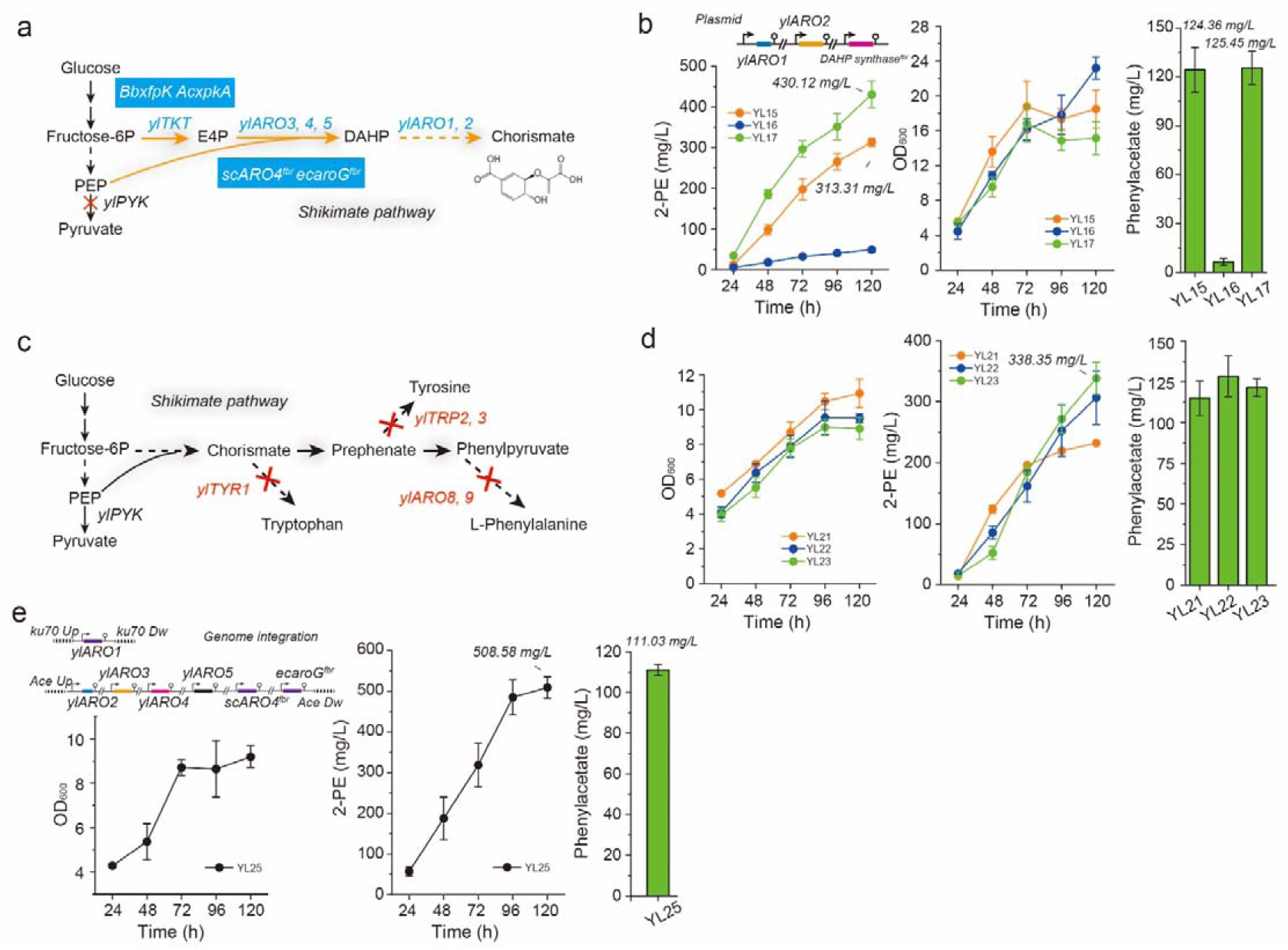
Relieving bottlenecks in shikimate pathway. (a) The shikimate pathway of *Y. lipolytica*; (b) Phenylacetate, 2-PE and cell growth profile of strains coexpressing ylARO1, ylARO2 and the feedback-resistant DAHP synthases encoding genes; (c) Deletion of amino acids, including phenylalanine, tryptophan, and tyrosine in *Y. lipolytica*; (d) Phenylacetate, 2-PE and cell growth profile of mutant strains deficient in phenylalanine, tryptophan, and tyrosine formation; (e) Phenylacetate, 2-PE and cell growth profile of mutant strains (deficient in phenylalanine, tryptophan, and tyrosine formation) expressing the shikimate pathway. All experiments were performed in triplicate and error bars represent standard deviations (SD).

On the other hand, we sought to eliminate the synthesis of aromatic amino acids byproducts, including phenylalanine, tryptophan, and tyrosine. For this purpose, five genes (Figure 3c) were chosen as knockout targets, including *ylTYR1* (*YALI0F17644g*, encoding prephenate dehydrogenase), *ylTRP2* (*YALI0D11110g*, encoding anthranilate synthase), *ylTRP3* (*YALI0E14751g*, encoding anthranilate synthase), *ylARO8* (*YALI0E20977g*, encoding aromatic amino acid aminotransferase), and *ylARO9* (*YALI0C05258g*, encoding aromatic amino acid aminotransferase). By sequentially deleting *ylTYR1, ylTRP2, ylTRP3, ylARO8*, and *ylARO9* in strain po1fk (which is Ku70-deficient), we got strains **YL18** (auxotroph of tryptophan), **YL19** (auxotroph of tryptophan and tyrosine), and **YL20** (auxotroph of tryptophan, tyrosine and phenylalanine). Then the *de novo* 2-PE pathway (*ylPAR4-ylARO10-ylARO7-ylPHA2-scARO7* ^*G141S*^) was introduced into these auxotrophic strains (YL18, 19 and 20), leading to strains YL21, YL22 and YL23. In comparison to the parental strain **YL6** (87.63 ± 4.67 mg/L of 2-PE titer), strains, **YL21, YL22**and **YL23** led to 2-PE production (Figure 3d) increased by 2.65-fold (231.84 ± 1.46 mg/L), 3.49-fold (306.30 ± 43.57mg/L), and 3.86-fold (338.35 ± 26.41 mg/L), respectively, despite that negative effect was observed in the cell growth. This result suggests that blocking of aromatic amino acids formation may further relieve allosteric regulation of DAHP synthase by L-Phe, L-Tyr and L-Trp, and simultaneously mitigate precursor competition.

Taken together, we have proved that introduction of feedback-resistant heterologous DAHP synthases and blocking aromatic amino acids formation are both effective to debottlenecking the shikimate pathway in *Y. lipolytica*. We next combined these two strategies to boost the 2-PE yield. Genes encoding *ylARO1* and *ylARO2-ylARO3-ylARO4-ylARO5-scARO4* ^*K229L*^ *-aroG* ^*S180F*^ were sequentially integrated at genome loci of *ku70* and *YALI0E30965g*, to generate the chassis strain **YL24**. By expression of the 2-PE pathway (*ylPAR4-ylARO10-ylARO7-ylPHA2-scARO7* ^*G141S*^) in strain **YL24**, the engineered strain produced 508.58 ± 26.52 mg/L of 2-PE (strain **YL25**, Figure 3e), which was 1.18-fold, 1.50-fold, and 59.97-fold increase over the **YL17, YL23** and the starting strain **YL0**, respectively.

On the other hand, the operation of shikimate pathway is driven by the upstream precursors erythrose-4-phosphate (E4P, that is derived from pentose phosphate pathway) and phosphoenolpyruvate (PEP, that is derived from glycolysis). To further unlock the shikimate pathway, we next turned to rewire the carbon flows in the central carbon metabolism to optimize the supply of both E4P and PEP.

### Removing pentose phosphate pathway bottleneck to boost precursor E4P

Metabolic flux analysis indicates that the available carbon flux toward E4P is as least one order of magnitude lower than the flux to PEP in yeast ^34^, which suggested the availability of E4P is the key step to maximizing flux to shikimate pathway. Overexpression of gene *ylTKT* (*YALI0E06479g*, encoding transketolase) improved 2-PE titer to 598.36 ± 0.46 mg/L (strain **YL26**, Supplementary Figure S4), a 1.18-fold increase over the strain **YL25**. This result indicates that the supply of E4P indeed was a bottleneck.

In the pentose phosphate pathway, phosphoketolase splits fructose-6-phosphate into E4P and acetyl-phosphate (Figure 4a) ^34, 53-55^. In a recent study, overexpression of heterologous phosphoketolase BbxfpK from *Bifidobacterium breve* led to a 5.4-fold increase of intracellular E4P concentration in *S. cerevisiae* ^53^. We therefore introduced the codon-optimized phosphoketolase BbxfpK and AcxpkA (originating from *Acidobacterium capsulatum*) into our yeast chassis to generate strains **YL27** and **YL28**, respectively. However, we observed declined 2-PE production (Supplementary Figure S4) in both strains **YL27** (191.72 ± 7.97mg/L of 2PE) and **YL28** (146.76 ± 9.64 mg/L of 2PE). As plasmids genetic instability (i.e. unequal distribution/propagation of plasmid) was observed in our previous research, we speculated that a similar effect was occurred in strains **YL27** and **YL28**. To resolve this issue, linearized gene fragments (see **Methods**) containing *pYLXP’-BbxfpK-ylPAR4-ylARO10-ylARO7-ylPHA2-scARO7* ^*G141S*^ and *pYLXP’-AcxpkA-ylPAR4-ylARO10-ylARO7-ylPHA2-scARO7* ^*G141S*^, were integrated at the pBR docking site of strain **YL24**, generating strains **YL29** and **YL30**, respectively. As expected, shake flask cultivation of strains **YL29** and **YL30** led to a 133.64% (679.66 ± 1.14 mg/L) and 139.33% (708.58 ± 56.52 mg/L) improvement in 2-PE titer (Supplementary Figure S4), compared to the parental strain **YL25**. This result clearly demonstrates that manipulation of the phosphoketolase pathway is a feasible strategy to channeling carbon flux toward E4P. After this, we integrated genes *ylTKT, BbxfpK* and *AcxpkA* at 26s rDNA site in strain **YL24** with the helper plasmid prDNAloxP-*ylTKT-BbxfpK-Acxpk*, obtaining strain **YL31**. However, expression of the 2-PE pathway (*ylPAR4-ylARO10-ylARO7-ylPHA2-scARO7* ^*G141S*^) in strain **YL31** gave almost same amount of 2-PE (strain **YL32**, 719.67 ± 51.14 mg/L, Figure 4b) compared to strain **YL30**. We speculate that the engineered yeast chassis has sufficient E4P flux to maintain the shikimate pathway.

**Figure 4.**
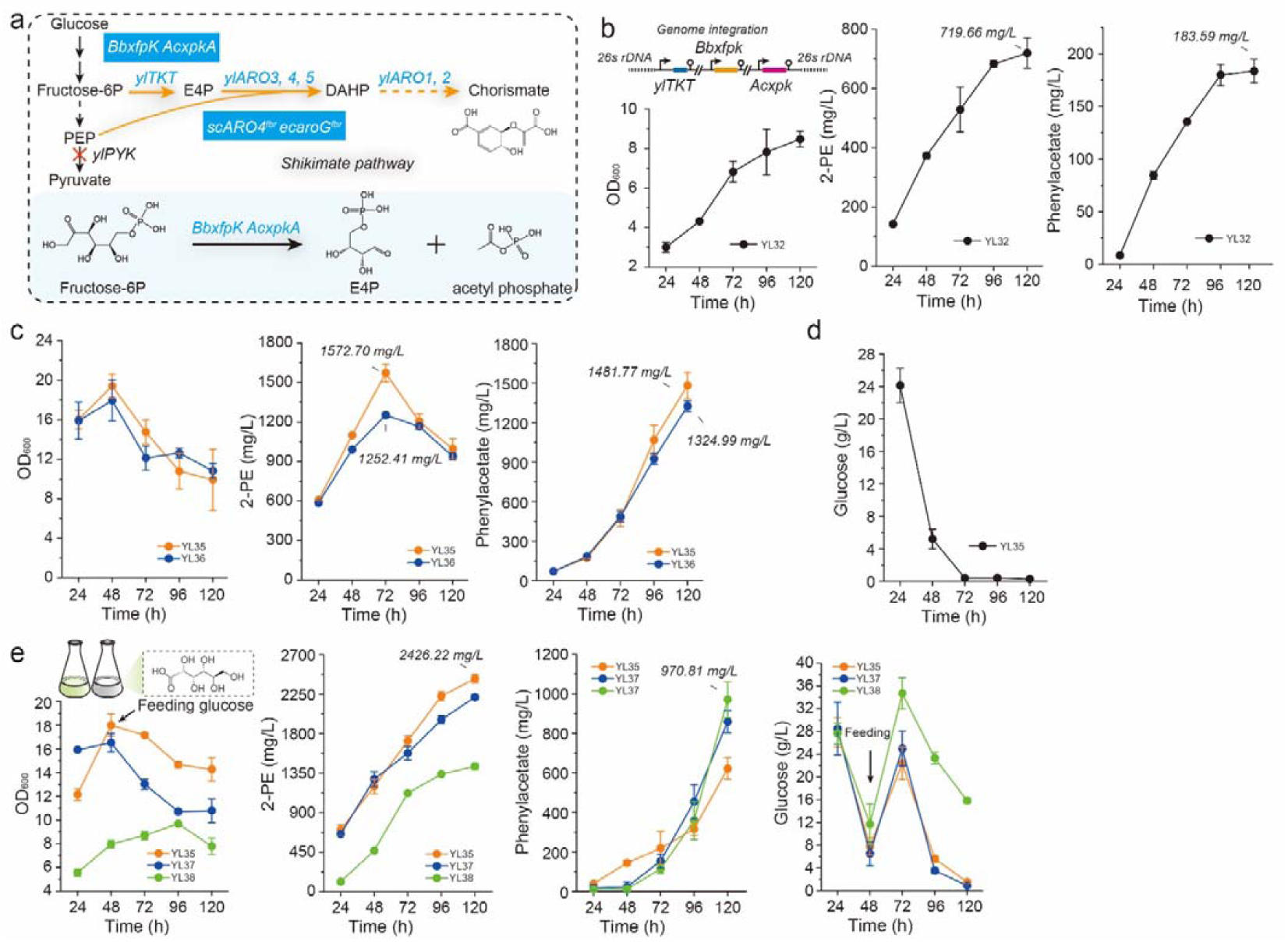
Rewiring carbon distribution toward precursors E4P and PEP. (a) Phosphoketolases provide precursor E4P to drive shikimate pathway; (b) Time profiles of 2-PE, cell growth, and phenylacetate of strains with chromosomal integration of transketolase ylTKT, phosphoketolases BbxfpK and AcxpkA at genomic 26s rDNA sites; (c) Time profile of 2-PE, cell growth, and phenylacetate of strains with pyruvate kinase ylPYK deletion in YPD medium; (d) Time profiles of glucose consumption of strain YL35; (e) Improving 2-PE titer by feeding glucose and minimizing byproduct phenylacetate by deleting aldehyde dehydrogenase. All experiments were performed in triplicate and error bars represent standard deviations (SD).

### Removing glycolytic pathway bottleneck to boost precursor PEP

To shift the metabolic equilibrium and further improve carbon flux toward shikimate pathway, we moved to boost the precursor availability of PEP. Previous study ^56^ found that blocking the reaction of PEP to pyruvate by deleting pyruvate kinase could dramatically increase the intracellular pools of PEP. To validate this strategy in *Y. lipolytica*, we attempted to delete pyruvate kinase encoding gene *ylPYK* (*YALI0F09185g*). Reports found that yeast with pyruvate kinase deficiency could not grow in the complete synthetic media (CSM) with glucose as sole carbon source ^57^. To successfully delete *ylPYK* (*YALI0F09185g*) in YL31 background chassis, we replenished 0.5 g/L of acetate in the selective CSM-plate (see **Methods**) to rescue the growth phenotype and obtained stain **YL33**. However, strain **YL33** carrying the 2-PE pathway (*ylPAR4-ylARO10-ylARO7-ylPHA2-scARO7* ^*G141S*^) showed a remarkable decrease in both biomass and 2-PE production in the synthetic minimal media, even with the feeding of acetate ^10^ (stain **YL34**, Supplementary Figure S5). Interestingly, when the same strain (**YL33**) was cultivated in YPD medium with 5 g/L sodium acetate, we observed robust cell growth (YL33 can also grow in YPD medium without acetate, as shown in Supplementary Figure S6). To this end, we integrated the 2-PE pathways (*ylPAR4-ylARO10-ylARO7-ylPHA2-scARO7* ^*G141S*^) into chassis **YL33** and cultivated YL33 with YPD medium containing 5 g/L sodium acetate (YPA medium). The generated strain **YL35** significantly improved 2-PE production to 1572.70 ± 67.82 mg/L in YPA medium (Figure 4c). The control strain **YL36** (integration of 2-PE pathway *ylPAR4-ylARO10-ylARO7-ylPHA2-scARO7* ^*G141S*^ in strain **YL31**) produced 1252.41 ± 19.02 mg/L of 2-PE with YPA media under the same cultivation conditions. This result confirmed that blocking PEP consumption by deletion of yl*PYK* is a promising strategy for enlarging flux toward shikimate pathway.

Interestingly, the highest 2-PE titer of strain **YL35** (1572.70 ± 67.82 mg/L) was obtained at 72 h, and subsequently, 2-PE was gradually oxidized to PEA (phenylacetic acid), which reached 1481.8 mg/L at 120 h (Figure 4c). We speculate that the metabolic bottleneck may be shifted from E4P/PEP to NADH due to the exhaustion of glucose (Figure 4d): the engineered cells need to oxidize 2-PE to generate NADH and maintain metabolite homeostasis. To resolve this issue, feeding with 40 g/L of glucose effectively mitigated the synthesis of PEA (Figure 5e). As a result, we obtained a final 2-PE titer at 2426.22 ± 48.33 mg/L (Figure 5e). To further improve 2-PE titer, we next sought to eliminate byproducts formation in strain **YL35** by knocking out *ylALD2, ylALD3*, and *ylHPD* (encoding aldehyde dehydrogenases), resulting in strains **YL37** and **YL38**. Deletion of genes *ylALD2* and *ylALD3* (strain **YL37**) did not significantly improve 2-PE production (2214.14 ± 26.48 mg/L, Figure 4e), in addition, further deletion of gene *ylHPD* (strain **YL38**) led a remarkable decrease in both cell growth and 2-PE production. This is possibly due to the reversibility of *ylHPD* that contributes to the reduction of PEA to 2-PE.

**Figure 5.**
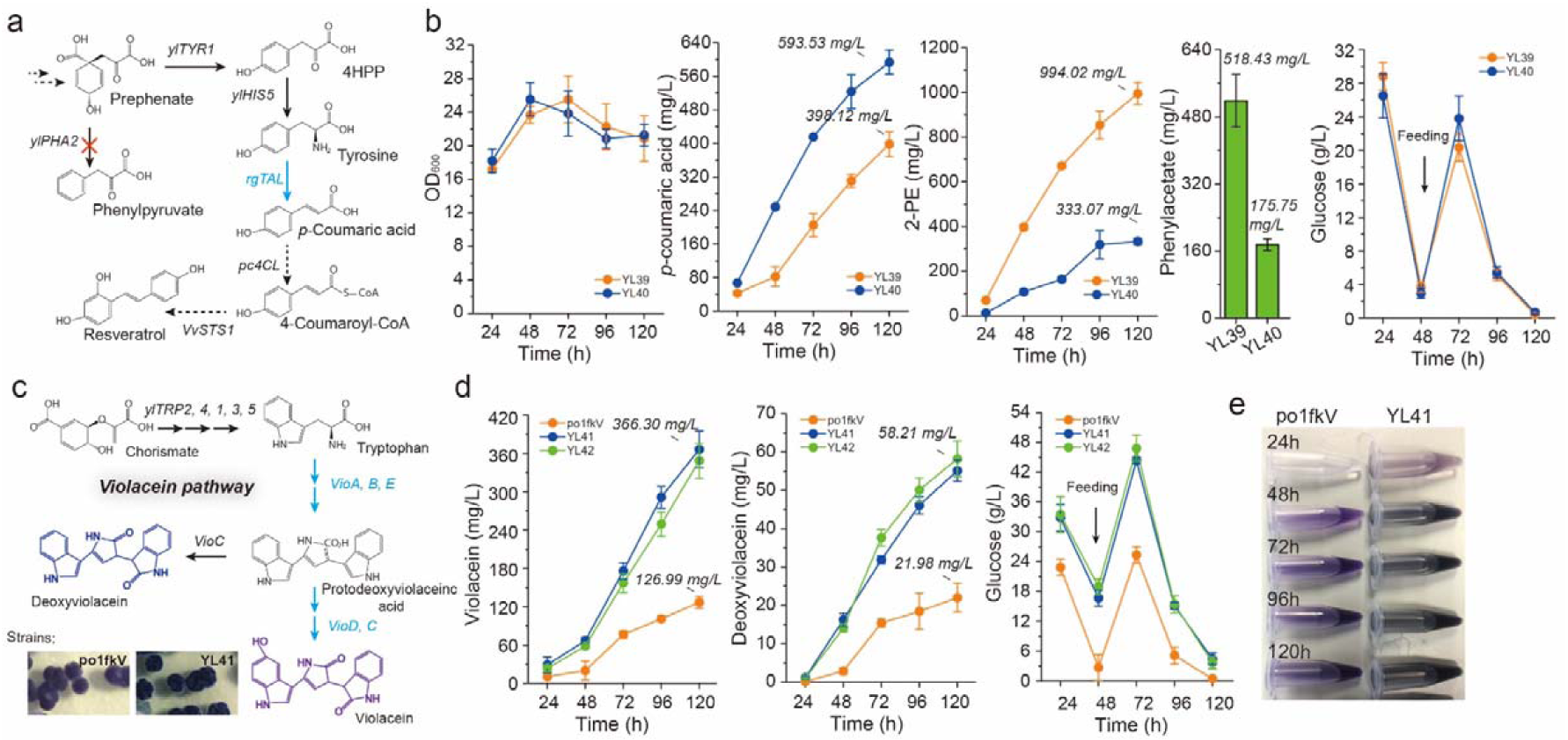
Harnessing *Y. lipolytica* chassis strain for p-coumaric acid and violacein production. (a) The de novo pathway for *p*-coumaric acid synthesis; (b) Phenylacetate, cell growth, *p*-coumaric acid, 2-PE and glucose consumption profile of the engineered chassis from the *de novo* pathway; (c) Violacein biosynthetic pathway and yeast colonies with violacein accumulation; (d) Time profiles of violacein, deoxyviolacein and glucose consumption of the violacein-producing strain. (e) Violacein cell culture harvested from synthetic minimal media (strain po0fkV) and YPD media (YL41). All experiments were performed in triplicate and error bars represent standard deviations (SD).

In summary, we obtained the *Y. lipolytica* platform strain **YL33** with optimized metabolic flux toward shikimate pathway. This chassis strain was validated for *de novo* production of 2-PE at 2426.22 ± 48.33 mg/L (strain **YL35**), a 286-fold increase over the initial strain (8.48 ± 0.50 mg/L). This is the highest 2-PE titer from the *de novo* pathway reported to date. To demonstrate the utility of this 2-PE-producing platform, we turned to synthesize other aromatic-derived compounds, including *p*-coumaric acid (the derivative of tyrosine), resveratrol (the derivatives of *p*-coumaric acid) and violacein (the derivative of tryptophan) in the following section.

### Extend the 2-PE platform for production of aromatic derivatives

In line with the optimization of 2-PE production, we decided to extend the yeast chassis to produce other aromatic derivatives, namely *p*-coumaric acid and violacein. *p*-Coumaric acid is the deamination product of tyrosine (Figure 5a), which is the universal precursor in the synthesis of flavonoids, aromatic polyketides, and lignin polyphenols ^36, 58^. By simply overexpressing a codon-optimized tyrosine ammonia lyase *RgTAL* (originating from *Rhodotorula toruloides*) ^34^and *ylTYR1* (that has been deleted in the platform strain) in the yeast chassis **YL33**, the resulting strain **YL39** produced 398.12 ± 29.39 mg/L of *p*-coumaric acid with glucose supplementation (Figure 5b). However, 994.02 ± 49.68 mg/L of 2-PE was detected, indicating a strong competing flux from the endogenous Ehrlich pathway. Thus, we next blocked 2-PE synthesis by deleting gene *ylPHA2* (encoding prephenate dehydratase), the obtained strain **YL40** produced 593.53 ± 28.75 mg/L of *p*-coumaric acid with yield at 7.42 ± 0.36 mg/g glucose (Figure 5b), a 1.49-fold increase compared to the control strain. We further extend the *p*-coumaric acid pathway for resveratrol synthesis by introducing genes *pc4CL* (4-coumarate-CoA ligase from *Petroselinum crispum*) and *VvSTS1* (resveratrol synthase from *Vitis vinifera*) ^59^. Unexpected, only 12.67 (± 2.23) mg/L of resveratrol was detected in shake flasks (Supplementary Figure S7), indicating the existence of other rate-limiting precursors such as malonyl-CoA in the chassis strain.

Violacein is a naturally-occurring purple pigment with proven chemotherapeutic activity against tumors and cancers ^60-61^. Both violacein and its derivate deoxyviolacein have demonstrated a broad range of biological activities ^60^. By introducing violacein biosynthetic genes (VioA, VioB, VioC, VioD, and VioE, Figure 5c), we previously obtained violacein production around 31 mg/L in *Y. lipolytica* ^30, 62^. Surprisingly, overexpression of genes *ylTRP2, ylTRP3, VioA, VioB, VioC, VioD*, and *VioE* in strain **YL33** led to the accumulation of black pigment in the fermentation broth (Supplementary Figure S8). Violacein and deoxyviolacein reached 366.30 ± 28.99 mg/L and 55.12 ± 2.81 mg/L (Figure 5d), which were 2.88-fold and 2.51-fold increase over the control strain (**po1fkV**, 126.99 ± 9.08 mg/L of violacein and 21.98 ± 3.75 mg/L of deoxyviolacein) and 5.23-fold and 10.44-fold higher than the production reported in our previous work ^62^. To further improve the violacein and deoxyviolacein production, we enhanced tryptophan availability by overexpressing genes *ylTRP5, ylTRP4, ylTRP3, ylTRP2*, and *ylTRP1*. However, no significant improvements of violacein (348.43 ± 27.21 mg/L) and deoxyviolacein (58.21 ± 4.62 mg/L) titer were observed in strain **YL42**, suggesting that the supply of tryptophan was sufficient in strain **YL41** and there might be other limiting steps in the violacein pathway. Besides, we also attempted to block the chorismite-competing pathway by deleting *ylARO7*. No positive colonies were obtained in the background of strain **YL41**with several attempts, but *ylARO7* could be deleted in po1fk. The specific reason of our inability to delete *ylARO7* is not clear, which might be due to the metabolic burden of violacein accumulated in the cell negatively impacting cell fitness. However, the colonies with genome integration of violacein pathway (po0fkV) in YPD plate showed that the using of same promoters and terminators may lead to the occurrence of genetic instability: a minor fraction of the re-streaked colonies lost the ability to produce violacein (Supplementary Figure 12). Thus, the frequency of using multiple identical promoters and terminators should be minimized when integrating several genes in one genome locus in *Y. lipolytica*.

Interestingly, when the violacein-producing strain **YL33** was tested in the complete synthetic media (CSM-Leu), we obtained less than 50% (180 mg/L) of violacein, compared to the violacein production with YPD media (366.3 mg/L). We speculated that the presence of tryptophan in the CSM-leu media (consisting of 50 mg/L tryptophan) might feedback inhibit the expression of critical enzymes in the violacein biosynthetic pathway. Switching to a CSM-Leu media with reduced tryptophan (20 mg/L) indeed improved violacein production to 271 mg/L, indicating the presence or buildup of tryptophan strongly represses the enzyme activity in the violacein pathway. The high violacein titer (366.3 mg/L) obtained in the YPD media might be due to the slow release of tryptophan from yeast extract and peptone, which may not render feedback inhibition effect on the violacein biosynthetic pathway. While exact inhibition constants (*K*i) of tryptophan to critical enzymes in the violacein biosynthetic pathway will be important for us to optimize amino acid composition in the CSM media, the use of chromosomally integrated strain cultivated in YPD media may bypass this feedback inhibition effect and improve aromatic compounds production.

Additionally, future investigations of strains **YL25, YL40**, and **YL42** should be performed under fed-batch cultivation with bench-top bioreactors and pilot-scale tests to improve the titer, yield and productivity of 2-PE, *p*-coumaric acid, and violacein. Major bioprocess considerations should be targeted at optimizing the culture medium, relieving 2-PE toxicity and promoting cell fitness, as well as reducing the separation and purification cost for 2-PE, *p*-coumaric acid, and violacein. Biphasic fermentation with end-product stripping might be an effective way to remove product inhibition. Considering that lignin is widely distributed in nature and it is made up with a variety of aromatic monomers, co-culture of *Y. lipolytica* with lignin-degradation microbes, including *Pseudomonas putida* and *Rhodococcus jostii et al*, with solid-state or liquid co-cultivation, might provide an attractive approach to valorize lignin and produce other complex aromatic derivatives.

In summary, by circumventing the intrinsic limitations of the endogenous shikimate pathway, we successfully engineered an oleaginous yeast chassis that produces 2426.22 ± 48.33 mg/L of 2-PE, 593.53 ± 28.75 mg/L of *p*-coumaric acid, 366.30 ± 28.99 mg/L of violacein, and 55.12 ± 2.81 mg/L of deoxyviolacein. To the best of our knowledge (Supplementary Table4), this result represents the highest 2-PE, violacein and deoxyviolacein titer from the *de novo* shikimate pathway. This report highlights the prominent metabolic characteristics of *Y. lipolytica* as chassis for production of various aromatic derivatives.

## Conclusions

*Y. lipolytica* is an oleaginous yeast with superior metabolic capability to produce a large portfolio of fuels, oleochemicals and natural products. In this work, we systematically overcame the rate-limiting steps of the endogenous shikimate pathway and build a sustainable biorefinery chassis strain for *de novo* synthesis of aromatics. We determined that relieving the feedback-inhibition of DAHP synthases is critical to channel flux to shikimate pathway. We also demonstrated that eliminating by-product formation (L-phe, L-Tyr and L-Trp) plays an important role in mitigating the feedback regulation of DAHP synthase. Overexpression of phosphoketolase and deletion of pyruvate kinase provided a sustained metabolic source for E4P and PEP, which forms the driving force to lead carbon flux through shikimate pathway. To demonstrate the utility of our engineered *Y. lipolytica* chassis strain, three natural products, 2-PE, *p*-coumaric acid and violacein, which were respectively derived from phenylalanine, tyrosine and tryptophan, were chosen to test the chassis performance. With the engineered chassis *Y. lipolytica*, we obtained 2426.22 ± 48.33 mg/L of 2-PE, 593.53 ± 28.75 mg/L of *p*-coumaric acid, 12.67 ± 2.23 mg/L of resveratrol, 366.30 ± 28.99 mg/L of violacein, and 55.12 ± 2.81 mg/L of deoxyviolacein. These results highlight the metabolic versatility of *Y. lipolytica* and the native shikimate pathway could be unlocked to produce a variety of aromatics and natural products with economic values.

## Material and Methods

### Strains, plasmid, primers, and chemicals

All stains of engineered *Y. lipolytica*, including the genotypes, recombinant plasmids, and primers have been listed in Supplementary Table1 and 2. Chemicals used in this study were all purchased from Sigma-Aldrich. Codon-optimized heterologous synthetic genes, including genes *BbxfpK, AcxpkA, RgTAL, VvSTS1*, and *Pc4CL2*, were ordered from GENEWIZ (Suzhou, China). Codon-optimization was performed with the IDT website.

### Shake flask cultivations

For performing shake flask cultivations, seed culture was carried out in the shaking tube with 2 mL seed culture medium at 30 °C and 250 r.p.m. for 48 h. Then, 0.8 mL of seed culture was inoculated into the 250 mL flask containing 30 mL of fermentation medium and grown under the conditions of 30 °C and 250 r.p.m. for 120 h. One milliliter of cell suspension was sampled every 24h for OD_600_, glucose, and desired metabolisms measurements.

Seed culture medium used in this study included the yeast complete synthetic media regular media (CSM, containing glucose 20.0 g/L, yeast nitrogen base without ammonium sulfate 1.7 g/L, ammonium sulfate 5.0 g/L, and CSM-Leu 0.74 g/L) and complex medium (YPD, containing glucose 20.0 g/L, yeast extract 10.0 g/L, and peptone 20.0 g/L). Fermentation medium used in this study also included the yeast complete synthetic media regular media (CSM, containing glucose 40.0 g/L, yeast nitrogen base without ammonium sulfate 1.7 g/L, ammonium sulfate 5.0 g/L, and CSM-Leu 0.74 g/L) and complex acetate medium (YPA, containing glucose 40.0 g/L, yeast extract 10.0 g/L, peptone 20.0 g/L, and sodium acetate 5.0 g/L).

### Yeast transformation and screening of high-producing strains

The standard protocols of *Y. lipolytica* transformation by the lithium acetate method were described as previously reported ^63-64^. In brief, one milliliter cells was harvested during the exponential growth phase (16-24 h) from 2 mL YPD medium (yeast extract 10 g/L, peptone 20 g/L, and glucose 20 g/L) in the 14-mL shake tube, and washed twice with 100 mM phosphate buffer (pH 7.0). Then, cells were resuspended in 105 uL transformation solution, containing 90 uL 50% PEG4000, 5 uL lithium acetate (2M), 5 uL boiled single stand DNA (salmon sperm, denatured) and 5 uL DNA products (including 200-500 ng of plasmids, lined plasmids or DNA fragments), and incubated at 39 °C for 1 h, then spread on selected plates. It should be noted that the transformation mixtures needed to be vortexed for 15 seconds every 15 minutes during the process of 39 °C incubation. The selected markers, including leucine, uracil and hygromycin, were used in this study. All engineering strains after genetic manipulations were performed optimized screening by the shaking tube cultivations, and the optimal strain was used to perform shaking flask (these data have been shown in Supplementary Materials).

### Single-gene and multi-genes expression vectors construction

In this work, the YaliBrick plasmid pYLXP’ was used as the expression vector ^30^. The process of plasmid constructions also has been reported ^65^. In brief, recombinant plasmids of pYLXP’-*xx* (a single gene) were built by Gibson Assembly of linearized pYLXP’ (digested by *SnaBI* and *KpnI*) and the appropriate PCR-amplified or synthetic DNA fragments. Multi-genes expression plasmids were constructed based on restriction enzyme subcloning with the isocaudamers *AvrII* and *NheI* ^66-67^. All genes were respectively expressed by the TEF promoter with intron sequence and XPR2 terminator, and the modified DNA fragments and plasmids were sequenced by Quintarabio.

### Gene knockout

A marker-free gene knockout method based on Cre-*lox* recombination system was used as previously reported ^68^. For performing gene knockout, the upstream and downstream sequences (both 1000 bp) flanking the deletion targets were PCR-amplified. These two fragments, the *loxP-Ura/Hyr-loxP* cassette (digested from plasmid *pYLXP’-loxP-Ura/Hyr by AvrII* and *salI*), and the gel-purified plasmid backbone of pYLXP’(linearized by *AvrII* and *salI*) were joined by Gibson Assembly, giving the knockout plasmids pYLXP’-*loxP-Ura/Hyr-xx* (xx is the deletion target). Next, the knockout plasmids were sequence-verified by Quintarabio. Then, the gene knockout cassettes were PCR-amplified from the knockout plasmids pYLXP’-*loxP-Ura/Hyr-xx*, and further transformed into *Y. lipolytica*. The positive transformants were determined by colony PCR. Ku70 (Po1f background) was knocked out by screening more than 200 yeast colonies. Other knockout strains were built on top of the Ku70-deficient strains. Subsequently, plasmid pYLXP’-*Cre* was introduced into the positive transformants and promoted the recombination of *loxP* sites, which recycle the selected marker. Finally, the intracellular plasmid pYLXP’-*Cre* was evicted by incubation at 30°C in YPD media for 48h. Here, *Ura* is the uracil marker, and *Hyr* is hygromycin marker. Specifically, for deletion of gene *ylPYK*, final concentration of 0.5, 1, 1.5, 2, 2.5, 5 g/L acetate was added into the selective CSM-Ura plate to rescue the cell growth. Besides integration of linearized plasmid, other genomic manipulations in platform strain **YL33**, including gene knockout and integration of desired genes, hygromycin resistance gene was used as the selective marker grown on YPD plate containing 200 mg/L hygromycin.

### Genomic integration of desired genes

In this work, genomic integration of desired genes were performed in two different ways: site-specific genomic integration plasmids or application of pBR docking platform by linearizing the plasmid pYLXP’ with digested enzyme *NotI*. Here, to meet specific requirements, we constructed three genomic integration plasmids pURLK, pURLA, and pHyLD, corresponding to the *Ku70, YALI0E30965g* (encoding acetyl-CoA hydrolase, named as Ace), and *YALI0E03212g* (encoding lactate dehydrogenase, named as LDH) genomic sites, respectively. The procedure of using these three plasmids was similar as that of gene knockout protocol. Plasmid maps and sequences for pURLK, pURLA, and pHyLD have been uploaded in the Supplementary Materials. Specifically, desired genes could be assembled into pURLX based on restriction enzyme subcloning by using the multiple cloning sites (see plasmid maps in SI files), and the integration cassettes of desired genes were retrieved by digesting plasmid pURLX-*xx* with enzyme *AvrII*. Additionally, the standard protocol of 26s rDNA genomic integration by plasmid prDNAloxP was used in this work, with detailed protocol described in previous work ^6^.

The application of pBR docking platform was achieved by linearizing the plasmid pYLXP’ with NotI restriction enzymes. It should be noted that, plasmids pYLXP’-*ylPAR4-ylARO10-ylARO7-ylPHA2-scARO7*^*G141S*^ and pYLXP’-*VioDCBAEI* have more than 2 *NotI* digestion sites, thus, we performed site-directed mutagenesis to change the NotI site to SnaBI site in plasmid pYLXP’, generating plasmid pYLXPs’. Subsequently, gene fragments in pYLXPs’-*VioDCBAE*, pYLXPs’-*ylPAR4-ylARO10-ylARO7-ylPHA2-scARO7* ^*G141S*^ and its derivates were linearized by *SnaBI* digestion, and then the linearized fragments were integrated at genomic pBR docking site.

### Quantification of biomass, glucose, 2-PE, p-coumaric acid, resveratrol, violacein and deoxyviolacein

Cell densities were monitored by measuring the optical density at 600 nm (OD_600_). The concentrations of 2-PE, phenylacetate, glucose, p-coumaric acid, resveratrol, violacein, and deoxyviolacein were all measured by high-performance liquid chromatography (HPLC) through Agilent HPLC 1220. In detail, 2-PE and penylacetate were measured at 215 nm under 40 °C (column oven temperature) with a mobile phase containing 50% (v/v) methanol in water at a flow rate of 0.5 mL/min equipped with a ZORBAX Eclipse Plus C18 column (4.6 × 100 mm, 3.5 μm, Agilent) and the VWD detector. The concentrations of glucose were measured by a Supelcogel−Carbohydrate column (Sigma, USA) and a refractive index detector with H_2_ SO_4_ (5 mM) as the mobile phase at a flow rate of 0.6 mL/min at 40 °C.

To quantify the concentration of *p*-coumaric acid, 0.1 mL of fermentation culture was mixed with 9-fold volume of absolute methanol (100% v/v), vortexed thoroughly, and centrifuged at 12000 r.p.m. for 10 min. The supernatants were analyzed at 304 nm under 40 °C (column oven temperature) with a mobile phase containing 45% (v/v) methanol in water at a flow rate of 0.5 mL/min equipped with a ZORBAX Eclipse Plus C18 column (4.6 × 100 mm, 3.5 μm, Agilent) and the VWD detector. To quantify the concentration of resveratrol, 0.25 mL of fermentation culture was mixed with an equal volume of ethyl acetate and appropriate glass beads, vortexed at 30 °C for 24 h, and centrifuged at 12000 r.p.m. for 10 min. Then, 100 uL supernatants of top organic layer was transferred to glass vial and evaporated to dryness, then resolubilized with 100 uL of methanol. Resveratrol samples were analyzed with the same HPLC protocols as *p*-coumaric acid.

To quantify the concentration of violacein and deoxyviolacein, 0.20 mL of fermentation culture was mixed with 5-fold volume of ethyl acetate and appropriate glass beads, vortexed at 30 °C for 24 h, and centrifuged at 12000 r.p.m. for 10 min. The supernatants of top organic layer were analyzed at 570 nm under 40 °C (column oven temperature) with a gradient method with two solvents, water (A) and methanol (B), at a flow rate of 0.4 mL/min equipped with a ZORBAX Eclipse Plus C18 column (4.6 × 100 mm, 3.5 μm, Agilent) and the VWD detector. The elution started with 100% of solvent A, the fraction of solvent A was decreased linearly from 100% to 20% (0-5 min), and maintained at 20% for 3 min (5-8 min), then the fraction of solvent A was increased from 20% to 100% (8-12 min), and maintained at 100% for 1 min (12-13 min).

## Supporting Information

Supporting information could be found in the online version of this article. The supporting information contains 4 supplementary Tables, supplementary methods on genomic integration and 12 supplementary figures. Supplementary Table S1, Strains and plasmids. Supplementary Table S2, Primers and Oligos. Supplementary Table S3, Strain performance with product titer. Supplementary Table S4, Comparison of 2-PE titer by various microbial hosts. Supplementary notes: theoretical yield of the 2-PE pathway. Supplementary Figure S1. Mathematical models of 2-PE yield. Supplementary Figures S2-S12 contain byproducts (phenylacetic acid) and end-product (2-PE, *p*-coumaric acid, resveratrol, violacein and deoxyviolacein) titer, cell growth profile, yeast colony morphology of the PYK-deficient strain, yeast colony with deep purple color for violacein-producing cells *et al*.

## Author contributions

PX conceived the topic and designed the study. YG performed genetic engineering and fermentation experiments with input from JM and YL. GY and PX wrote the manuscript. PX revised the manuscript.

## Acknowledgments

We would like to acknowledge Bill & Melinda Gates Foundation (grant number OPP1188443) and National Science Foundation (CBET-1805139) for financially supporting this project. The authors would also like to acknowledge the Department of Chemical, Biochemical and Environmental Engineering at University of Maryland Baltimore County for funding support. YG would like to thank the China Scholarship Council for funding support.

---

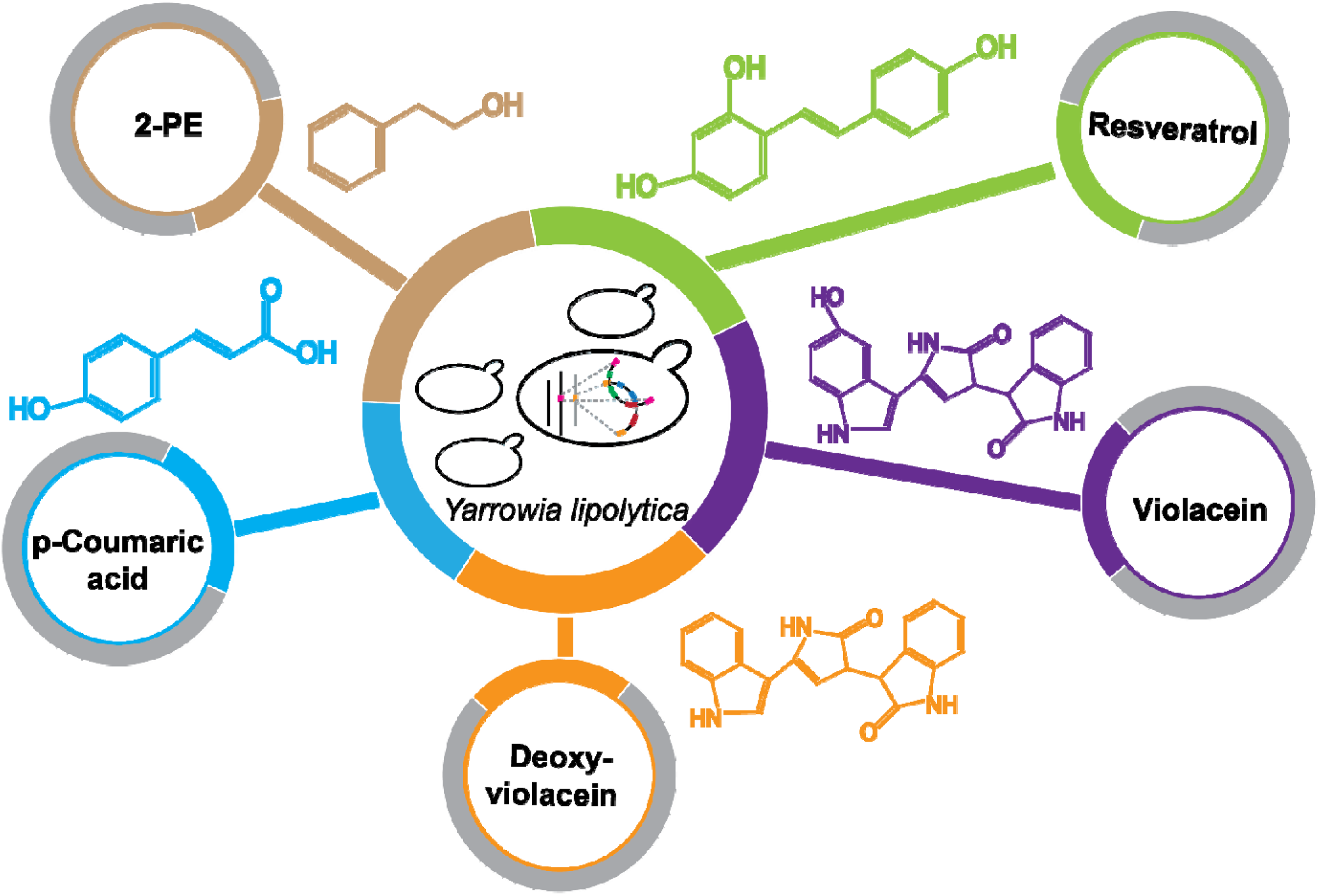
Graphic Table of Content.

## References

(1) Groenewald, M., Boekhout, T., Neuveglise, C., Gaillardin, C., Dijck, P. W., Wyss, M. (2014) Yarrowia lipolytica: safety assessment of an oleaginous yeast with a great industrial potential. Crit. Rev. Microbiol. 40, 187–206.

(2) Xu, P., Qiao, K., Stephanopoulos, G. (2017) Engineering oxidative stress defense pathways to build a robust lipid production platform in Yarrowia lipolytica. Biotechnol. Bioeng. 114 (7), 1521–1530.

(3) Xu, P., Qiao, K., Ahn, W. S., Stephanopoulos, G. (2016) Engineering Yarrowia lipolytica as a platform for synthesis of drop-in transportation fuels and oleochemicals. Proc. Nati. Acad. Sci. U. S. A. 113 (39), 10848–10853.

(4) Qiao, K., Wasylenko, T. M., Zhou, K., Xu, P., Stephanopoulos, G. (2017) Lipid production in Yarrowia lipolytica is maximized by engineering cytosolic redox metabolism. Nat. Biotechnol. 35 (2), 173–177.

(5) Abdel-Mawgoud, A. M., Markham, K. A., Palmer, C. M., Liu, N., Stephanopoulos, G., Alper, H. S. (2018) Metabolic engineering in the host Yarrowia lipolytica. Metab. Eng. 50, 192–208.

(6) Lv, Y., Edwards, H., Zhou, J., Xu, P. (2019) Combining 26s rDNA and the Cre-loxP System for Iterative Gene Integration and Efficient Marker Curation in Yarrowia lipolytica. ACS Synth. Biol. 8 (3), 568–576.

(7) Lv, Y., Marsafari, M., Koffas, M., Zhou, J., Xu, P. (2019) Optimizing oleaginous yeast cell factories for flavonoids and hydroxylated flavonoids biosynthesis. ACS Synth. Biol. 8 (11), 2514–2523.

(8) Palmer, C. M., Miller, K. K., Nguyen, A., Alper, H. S. (2020) Engineering 4-coumaroyl-CoA derived polyketide production in Yarrowia lipolytica through a β-oxidation mediated strategy. Metab. Eng. 57, 174–181.

(9) Lv, Y., Gu, Y., Xu, J., Zhou, J., Xu, P. (2020) Coupling metabolic addiction with negative autoregulation to improve strain stability and pathway yield. Metab. Eng. 61, 79–88.

(10) Liu, H., Marsafari, M., Wang, F., Deng, L., Xu, P. (2019) Engineering acetyl-CoA metabolic shortcut for eco-friendly production of polyketides triacetic acid lactone in Yarrowia lipolytica. Metab. Eng. 56, 60–68.

(11) Markham, K. A., Palmer, C. M., Chwatko, M., Wagner, J. M., Murray, C., Vazquez, S., Swaminathan, A., Chakravarty, I., Lynd, N. A., Alper, H. S. (2018) Rewiring Yarrowia lipolytica toward triacetic acid lactone for materials generation. Proc. Nati. Acad. Sci. U. S. A. 115 (9), 2096.

(12) Xue, Z., Sharpe, P. L., Hong, S. P., Yadav, N. S., Xie, D., Short, D. R., Damude, H. G., Rupert, R. A., Seip, J. E., Wang, J., Pollak, D. W., Bostick, M. W., Bosak, M. D., Macool, D. J., Hollerbach, D. H., Zhang, H., Arcilla, D. M., Bledsoe, S. A., Croker, K., McCord, E. F., Tyreus, B. D., Jackson, E. N., Zhu, Q. (2013) Production of omega-3 eicosapentaenoic acid by metabolic engineering of Yarrowia lipolytica. Nat. Biotechnol. 31 (8), 734–40.

(13) Xie, D., Jackson, E. N., Zhu, Q. (2015) Sustainable source of omega-3 eicosapentaenoic acid from metabolically engineered Yarrowia lipolytica: from fundamental research to commercial production. Appl. Microbiol. Biotechnol. 99 (4), 1599–610.

(14) Schwartz, C., Frogue, K., Misa, J., Wheeldon, I. (2017) Host and pathway engineering for enhanced lycopene biosynthesis in Yarrowia lipolytica. Front. Microbiol. 8, 2233.

(15) Zhang, R., Zhang, Y., Wang, Y., Yao, M., Zhang, J., Liu, H., Zhou, X., Xiao, W., Yuan, Y. (2019) Pregnenolone Overproduction in Yarrowia lipolytica by Integrative Components Pairing of the Cytochrome P450scc System. ACS Synth. Biol. 8, 2666–2678.

(16) Larroude, M., Celinska, E., Back, A., Thomas, S., Nicaud, J.-M., Ledesma-Amaro, R. (2018) A synthetic biology approach to transform Yarrowia lipolytica into a competitive biotechnological producer of β-carotene. Biotechnol. Bioeng. 115 (2), 464–472.

(17) Marsafari, M., Xu, P. (2020) Debottlenecking mevalonate pathway for antimalarial drug precursor amorphadiene biosynthesis in Yarrowia lipolytica. Metab. Eng. Commun. 10, e00121.

(18) Celińska, E., Ledesma-Amaro, R., Larroude, M., Rossignol, T., Pauthenier, C., Nicaud, J.-M. (2017) Golden Gate assembly system dedicated to complex pathway manipulation in Yarrowia lipolytica. Microb. Biotechnol. 10 (2), 450–455.

(19) Larroude, M., Park, Y. K., Soudier, P., Kubiak, M., Nicaud, J. M., Rossignol, T. (2019) A modular Golden Gate toolkit for Yarrowia lipolytica synthetic biology. Microb. Biotechnol. 12 (6), 1249–1259.

(20) Egermeier, M., Sauer, M., Marx, H. (2019) Golden Gate-based metabolic engineering strategy for wild-type strains of Yarrowia lipolytica. FEMS Microbiol. Lett. 366 (4), fnz022.

(21) Gao, S., Han, L., Zhu, L., Ge, M., Yang, S., Jiang, Y., Chen, D. (2014) One-step integration of multiple genes into the oleaginous yeast Yarrowia lipolytica. Biotechnol. Lett. 36 (12), 2523–2528.

(22) Gao, S., Tong, Y., Zhu, L., Ge, M., Zhang, Y., Chen, D., Jiang, Y., Yang, S. (2017) Iterative integration of multiple-copy pathway genes in Yarrowia lipolytica for heterologous beta-carotene production. Metab. Eng. 41, 192–201.

(23) Schwartz, C. M., Hussain, M. S., Blenner, M., Wheeldon, I. (2016) Synthetic RNA polymerase III promoters facilitate high-efficiency CRISPR-Cas9-mediated genome editing in Yarrowia lipolytica. ACS Synth. Biol. 5 (4), 356–9

(24) Yang, Z., Edwards, H., Xu, P. (2020) CRISPR-Cas12a/Cpf1-assisted precise, efficient and multiplexed genome-editing in Yarrowia lipolytica. Metab. Eng. Commun. 10, e00112.

(25) Gao, S., Tong, Y., Wen, Z., Zhu, L., Ge, M., Chen, D., Jiang, Y.,Yang, S. (2016) Multiplex gene editing of the Yarrowia lipolytica genome using the CRISPR-Cas9 system. J. Ind. Microbiol. Biotechnol. 43 (8), 1085–1093.

(26) Wagner, J. M., Williams, E. V.,Alper, H. S. (2018) Developing a piggyBac transposon system and compatible selection markers for insertional mutagenesis and genome engineering in Yarrowia lipolytica. Biotechnol. J. 13 (5), 1800022.

(27) Bredeweg, E. L., Pomraning, K. R., Dai, Z., Nielsen, J., Kerkhoven, E. J., Baker, S. E. (2017) A molecular genetic toolbox for Yarrowia lipolytica. Biotechnol. Biofuels 10, 2.

(28) Blazeck, J., Liu, L., Redden, H., Alper, H. (2011) Tuning gene expression in Yarrowia lipolytica by a hybrid promoter approach. Appl. Environ. Microbiol. 77 (22), 7905–7914.

(29) Liu, H., Marsafari, M., Deng, L.,Xu, P. (2019) Understanding lipogenesis by dynamically profiling transcriptional activity of lipogenic promoters in Yarrowia lipolytica. Appl. Microbiol. Biotechnol. 103 (7), 3167–3179

(30) Wong, L., Engel, J., Jin, E., Holdridge, B., Xu, P. (2017) YaliBricks, a versatile genetic toolkit for streamlined and rapid pathway engineering in Yarrowia lipolytica. Metab. Eng. Commun. 5, 68–77.

(31) Cui, Z., Gao, C., Li, J., Hou, J., Lin, C. S. K.,Qi, Q. (2017) Engineering of unconventional yeast Yarrowia lipolytica for efficient succinic acid production from glycerol at low pH. Metab. Eng. 42, 126–133.

(32) Spagnuolo, M., Shabbir Hussain, M., Gambill, L.,Blenner, M. (2018) Alternative substrate metabolism in Yarrowia lipolytica. Front. Microbiol. 9, 1077.

(33) Ledesma-Amaro, R., Nicaud, J.-M. (2016) Metabolic engineering for expanding the substrate range of Yarrowia lipolytica. Trends Biotechnol. 34 (10), 798–809.

(34) Liu, Q., Yu, T., Li, X., Chen, Y., Campbell, K., Nielsen, J., Chen, Y. (2019) Rewiring carbon metabolism in yeast for high level production of aromatic chemicals. Nat. Commun. 10 (1), 4976.

(35) Huccetogullari, D., Luo, Z. W.,Lee, S. Y. (2019) Metabolic engineering of microorganisms for production of aromatic compounds. Microb. Cell Fact. 18 (1), 41.

(36) Wang, J., Shen, X., Rey, J., Yuan, Q.,Yan, Y. (2018) Recent advances in microbial production of aromatic natural products and their derivatives. Appl. Microbiol. Biotechnol. 102 (1), 47–61.

(37) Xu, P., Marsafari, M., Zha, J., Koffas, M. (2020) Microbial coculture for flavonoid synthesis. Trends Biotechnol. 38 (7), 686–688.

(38) Furuya, T., Arai, Y., Kino, K. (2012) Biotechnological production of caffeic acid by bacterial cytochrome P450 CYP199A2. Appl. Environ. Microbiol. 78 (17), 6087.

(39) Lv, Y., Xu, S., Lyu, Y., Zhou, S., Du, G., Chen, J., Zhou, J. (2019) Engineering enzymatic cascades for the efficient biotransformation of eugenol and taxifolin to silybin and isosilybin. Green Chemistry 21 (7), 1660–1667.

(40) Nthangeni, M. B., Urban, P., Pompon, D., Smit, M. S., Nicaud, J. M. (2004) The use of Yarrowia lipolytica for the expression of human cytochrome P450 CYP1A1. Yeast 21 (7), 583–592.

(41) Hassing, E.-J., de Groot, P. A., Marquenie, V. R., Pronk, J. T., Daran, J.-M. G. (2019) Connecting central carbon and aromatic amino acid metabolisms to improve de novo 2-phenylethanol production in Saccharomyces cerevisiae. Metab. Eng. 56, 165–180.

(42) Luttik, M. A., Vuralhan, Z., Suir, E., Braus, G. H., Pronk, J. T., Daran, J. M. (2008) Alleviation of feedback inhibition in Saccharomyces cerevisiae aromatic amino acid biosynthesis: quantification of metabolic impact. Metab. Eng. 10 (3-4), 141–53.

(43) Li, Z., Wang, X.,Zhang, H. (2019) Balancing the non-linear rosmarinic acid biosynthetic pathway by modular co-culture engineering. Metab. Eng. 54, 1–11.

(44) Wang, R., Zhao, S., Wang, Z.,Koffas, M. A. G. (2020) Recent advances in modular co-culture engineering for synthesis of natural products. Curr. Opin. Biotechnol. 62, 65–71.

(45) Zhang, H., Pereira, B., Li, Z., Stephanopoulos, G. (2015) Engineering Escherichia coli coculture systems for the production of biochemical products. Proc. Nati. Acad. Sci. U. S. A. 112 (27), 8266.

(46) Celińska, E., Kubiak, P., Białas, W., Dziadas, M., Grajek, W. (2013) Yarrowia lipolytica: the novel and promising 2-phenylethanol producer. J. Ind. Microbiol. Biotechnol. 40 (3), 389–392.

(47) Lu, X., Wang, Y., Zong, H., Ji, H., Zhuge, B., Dong, Z. (2016) Bioconversion of L-phenylalanine to 2-phenylethanol by the novel stress-tolerant yeast Candida glycerinogenes WL2002-5. Bioengineered 7 (6), 418–423.

(48) Wang, Z., Jiang, M., Guo, X., Liu, Z., He, X. (2018) Reconstruction of metabolic module with improved promoter strength increases the productivity of 2-phenylethanol in Saccharomyces cerevisiae. Microb. Cell Fact. 17 (1), 60.

(49) Stark, A. H., Crawford, M. A., Reifen, R. (2008) Update on alpha-linolenic acid. Nutr. Rev. 66.

(50) Kim, B., Cho, B. R., Hahn, J. S. (2014) Metabolic engineering of Saccharomyces cerevisiae for the production of 2-phenylethanol via Ehrlich pathway. Biotechnol. Bioeng. 111 (1), 115–24.

(51) Rodriguez, A., Kildegaard, K. R., Li, M., Borodina, I., Nielsen, J. (2015) Establishment of a yeast platform strain for production of p-coumaric acid through metabolic engineering of aromatic amino acid biosynthesis. Metab. Eng. 31, 181–8.

(52) Gu, Y., Ma, J., Zhu, Y., Xu, P. (2020) Refactoring Ehrlich pathway for high-yield 2-phenylethanol production in Yarrowia lipolytica. ACS Synth Biol 9 (3), 623–633.

(53) Bergman, A., Hellgren, J., Moritz, T., Siewers, V., Nielsen, J., Chen, Y. (2019) Heterologous phosphoketolase expression redirects flux towards acetate, perturbs sugar phosphate pools and increases respiratory demand in Saccharomyces cerevisiae. Microb. Cell Fact. 18 (1), 25.

(54) Meadows, A. L., Hawkins, K. M., Tsegaye, Y., Antipov, E., Kim, Y., Raetz, L., Dahl, R. H., Tai, A., Mahatdejkul-Meadows, T., Xu, L., Zhao, L. S., Dasika, M. S., Murarka, A., Lenihan, J., Eng, D., Leng, J. S., Liu, C. L., Wenger, J. W., Jiang, H. X., Chao, L. L., Westfall, P., Lai, J., Ganesan, S., Jackson, P., Mans, R., Platt, D., Reeves, C. D., Saija, P. R., Wichmann, G., Holmes, V. F., Benjamin, K., Hill, P. W., Gardner, T. S., Tsong, A. E. (2016) Rewriting yeast central carbon metabolism for industrial isoprenoid production. Nature 537 (7622), 694–697.

(55) Bogorad, I. W., Lin, T. S., Liao, J. C. (2013) Synthetic non-oxidative glycolysis enables complete carbon conservation. Nature 502 (7473), 693–697.

(56) Gu, Y., Lv, X., Liu, Y., Li, J., Du, G., Chen, J., Rodrigo, L. A., Liu, L. (2019) Synthetic redesign of central carbon and redox metabolism for high yield production of N-acetylglucosamine in Bacillus subtilis. Metab. Eng. 51, 59–69.

(57) Xu, Y. F., Zhao, X., Glass, D. S., Absalan, F., Perlman, D. H., Broach, J. R., Rabinowitz, J. D. (2012) Regulation of yeast pyruvate kinase by ultrasensitive allostery independent of phosphorylation. Mol. Cell 48 (1), 52–62.

(58) Brey, L. F., Wlodarczyk, A. J., Bang Thofner, J. F., Burow, M., Crocoll, C., Nielsen, I., Zygadlo Nielsen, A. J., Jensen, P. E. (2019) Metabolic engineering of Synechocystis sp. PCC 6803 for the production of aromatic amino acids and derived phenylpropanoids. Metab. Eng. 57, 129–139.

(59) Lim, C., Fowler, Z., Hueller, T., Schaffer, S., Koffas, M. (2011) High-yield resveratrol production in engineered Escherichia coli. Appl. Environ. Microbiol. 3451–3460.

(60) Rodrigues, A. L., Gocke, Y., Bolten, C., Brock, N. L., Dickschat, J. S., Wittmann, C. (2012) Microbial production of the drugs violacein and deoxyviolacein: analytical development and strain comparison. Biotechnol. Lett. 34 (4), 717–20.

(61) Sun, H., Zhao, D., Xiong, B., Zhang, C., Bi, C. (2016) Engineering Corynebacterium glutamicum for violacein hyper production. Microb. Cell Fact. 15 (1), 148.

(62) Tong, Y., Zhou, J., Zhang, L., Xu, P. (2019) Engineering oleaginous yeast Yarrowia lipolytica for violacein production: extraction, quantitative measurement and culture optimization. bioRxiv, 687012.

(63) Gietz, R. D., Woods, R. A. (2002) Transformation of yeast by lithium acetate/single-stranded carrier DNA/polyethylene glycol method. Methods Enzymol. 350, 87–96.

(64) Chen, D. C., Beckerich, J. M.,Gaillardin, C. (1997) One-step transformation of the dimorphic yeast Yarrowia lipolytica. Appl. Microbiol.Biotechnol. 48 (2), 232–235.

(65) Wong, L., Holdridge, B., Engel, J., Xu, P., Genetic tools for streamlined and accelerated pathway engineering in Yarrowia lipolytica. In Microbial Metabolic Engineering: Methods and Protocols, Santos, C. N. S.; Ajikumar, P. K., Eds. Springer New York: New York, NY, 2019; pp 155–177.

(66) Xu, P., Koffas, M. A. G., Polizzi, K., Kontoravdi, C. (2013) Assembly of Multi-gene Pathways and Combinatorial Pathway Libraries Through ePathBrick Vectors. Methods Mol. Bio. 1073, 107–129.

(67) Xu, P., Vansiri, A., Bhan, N., Koffas, M. (2012) ePathBrick: A synthetic biology platform for engineering metabolic pathways in E. coli. ACS Synth. Biol. 1 (7), 256–266.

(68) Fickers, P., Le Dall, M. T., Gaillardin, C., Thonart, P., Nicaud, J. M. (2003) New disruption cassettes for rapid gene disruption and marker rescue in the yeast Yarrowia lipolytica. J. Microbiol. Methods 55 (3), 727–37.

