## Supplementary tables and figures for "Engineering *Yarrowia lipolytica* as a chassis for *de novo* synthesis of five aromatic-derived natural products and chemicals"

### **Supplementary Table**

***Table 1. Strains and plasmids used in this study***

| Names | Characteristics | | Reference |
| --- | --- | --- | --- |
| **Strains** | | | |
| po1g | Wild-type strain W29 (ATCC20460) derivate; W29 *ΔmatA* *Δxpr*2-332 *Δaxp*-2 *Δleu*2-270 pBR platform | | ^1^ |
| po1f | po1g derivate; Further deletion of gene *ura*; po1g *Δura*3 | | ^1^ |
| po1fk | po1f derivate; Further deletion of gene *ku70*; po1f *Δku70::loxP* | | ^2^ |
| YL0 | po1fk pYLXP’ | | This work |
| YL1 | po1fk pYLXP’-*ylPAR4* | | This work |
| YL2 | po1fk pYLXP’-*ylARO10* | | This work |
| YL3 | po1fk pYLXP’-*ylPHA2* | | This work |
| YL4 | po1fk pYLXP’-*ylARO7* | | This work |
| YL5 | po1fk pYLXP’-*ylPAR4-ylARO10-ylPHA2-ylARO7* | | This work |
| YL6 | po1fk pYLXP’-*ylPAR4-ylARO10-ylPHA2-ylARO7-ScARO7* *^G141S^* | | This work |
| YL7 | po1fk derivate; Further integration of genes *ylPAR4*, *ylARO10*, *ylPHA2*, *ylARO7*,and *ScARO7^G141S^* *at* *YALI0E30965g* site; po1fk *ylPAR4* *ylARO10* *ylPHA2* *ylARO7 ScARO7^G141S^::loxP* | | This work |
| YL8 | YL7 pYLXP’-*ylARO1* | | This work |
| YL9 | YL7 pYLXP’-*ylARO2* | | This work |
| YL10 | YL7 pYLXP’-*ylARO3* | | This work |
| YL11 | YL7 pYLXP’-*ylARO4* | | This work |
| YL12 | YL7 pYLXP’-*ylARO5* | | This work |
| YL13 | YL7 pYLXP’-*ylARO1-ylARO2* | | This work |
| YL14 | YL7 pYLXP’-*ylARO1-ylARO2-ylARO3-ylARO4-ylARO5* | | This work |
| YL15 | YL7 pYLXP’-*ylARO1-ylARO2-scARO4^K229L^* | | This work |
| YL16 | YL7 pYLXP’-*ylARO1-ylARO2-ecaroG^L175D^* | | This work |
| YL17 | YL7 pYLXP’-*ylARO1-ylARO2-ecaroG^S180F^* | | This work |
| YL18 | po1fk derivate; Further deletion of genes *ylTYR1*; po1fk *ΔylTYR1::loxP* | | This work |
| YL19 | YL18 derivate; Further Deletion of genes *ylTRP2*, and *ylTRP3*; po1fk *ΔylTYR1 ΔylTRP2 ΔylTRP3::loxP* | | This work |
| YL20 | YL19 derivate; Further Deletion of genes *ylARO8*, and *ylARO9*; po1fk *ΔylTYR1 ΔylTRP2 ΔylTRP3 ΔylARO8 ΔylARO9::loxP* | | This work |
| YL21 | YL18 pYLXP’-*ylPAR4-ylARO10-ylPHA2-ylARO7-ScARO7* *^G141S^* | | This work |
| YL22 | YL19 pYLXP’-*ylPAR4-ylARO10-ylPHA2-ylARO7-ScARO7* *^G141S^* | | This work |
| YL23 | YL20 pYLXP’-*ylPAR4-ylARO10-ylPHA2-ylARO7-ScARO7* *^G141S^* | | This work |
| YL24 | YL20 derivate; Further integration of genes *ylARO1, ylARO2, ylARO3, ylARO4, ylARO5, scARO4^K229L^, aroG^S180F^* at *YALI0E30965g* and *ku70* sites; po1fk *ΔylTYR1 ΔylTRP2 ΔylTRP3 ΔylARO8 ΔylARO9 ylARO1 ylARO2 ylARO3 ylARO4 ylARO5 scARO4^K229L^ aroG^S180F^::loxP* | | This work |
| YL25 | YL24 pYLXP’-*ylPAR4-ylARO10-ylPHA2-ylARO7-ScARO7* *^G141S^* | | This work |
| YL26 | YL24 pYLXP’-*ylTKT*-*ylPAR4-ylARO10-ylPHA2-ylARO7-ScARO7* *^G141S^* | | This work |
| YL27 | YL24 pYLXP’-*bbxfpK*-*ylPAR4-ylARO10-ylPHA2-ylARO7-ScARO7* *^G141S^* | | This work |
| YL28 | YL24 pYLXP’-*acxpkA*-*ylPAR4-ylARO10-ylPHA2-ylARO7-ScARO7* *^G141S^* | | This work |
| YL29 | YL24 derivate; Further integration of genes *bbxfpK*, *ylPAR4*, *ylARO10*, *ylPHA2*, *ylARO7*, *ScARO7^G141S^* at pBR platform; po1fk *ΔylTYR1 ΔylTRP2 ΔylTRP3 ΔylARO8 ΔylARO9 ylARO1 ylARO2 ylARO3 ylARO4 ylARO5 scARO4^K229L^ aroG^S180F^* *bbxfpK* *ylPAR4* *ylARO10* *ylPHA2* *ylARO7* *ScARO7^G141S^::Leu* | | This work |
| YL30 | YL24 derivate; Further integration of genes *acxpkA*, *ylPAR4*, *ylARO10*, *ylPHA2*, *ylARO7*, *ScARO7^G141S^* at pBR platform; po1fk *ΔylTYR1 ΔylTRP2 ΔylTRP3 ΔylARO8 ΔylARO9 ylARO1 ylARO2 ylARO3 ylARO4 ylARO5 scARO4^K229L^ aroG^S180F^ acxpkA ylPAR4 ylARO10* *ylPHA2* *ylARO7* *ScARO7^G141S^::Leu* | | This work |
| YL31 | YL24 derivate; Further integration of genes *ylTKT*, *BbxfpK*, and *Acxpk* at 26s rDNA site; po1fk *ΔylTYR1 ΔylTRP2 ΔylTRP3 ΔylARO8 ΔylARO9 ylARO1 ylARO2 ylARO3 ylARO4 ylARO5 scARO4^K229L^ aroG^S180F^ ylTKT* *bbxfpK acxpk::loxP* | | This work |
| YL32 | YL31 pYLXP’-*ylPAR4-ylARO10-ylPHA2-ylARO7-ScARO7* *^G141S^* | | This work |
| YL33 | YL31 derivate; Further deletion of gene *ylPYK*; po1fk *ΔylTYR1 ΔylTRP2 ΔylTRP3 ΔylARO8 ΔylARO9 ΔylPYK ylARO1 ylARO2 ylARO3 ylARO4 ylARO5 scARO4^K229L^ aroG^S180F^ ylTKT* *bbxfpK acxpk::loxP* | | This work |
| YL34 | YL33 pYLXP’-*ylPAR4-ylARO10-ylPHA2-ylARO7-ScARO7* *^G141S^* | | This work |
| YL35 | YL33 derivate; Further integration of genes *ylPAR4*, *ylARO10*, *ylPHA2*, *ylARO7*, *ScARO7^G141S^* at pBR platform; po1fk *ΔylTYR1 ΔylTRP2 ΔylTRP3 ΔylARO8 ΔylARO9 ΔylPYK ylARO1 ylARO2 ylARO3 ylARO4 ylARO5 scARO4^K229L^ aroG^S180F^ ylTKT* *bbxfpK acxpk ylPAR4* *ylARO10* *ylPHA2* *ylARO7* *ScARO7^G141S^::Leu* | | This work |
| YL36 | YL31 derivate; Further integration of genes *ylPAR4*, *ylARO10*, *ylPHA2*, *ylARO7*, *ScARO7^G141S^* at pBR platform; po1fk *ΔylTYR1 ΔylTRP2 ΔylTRP3 ΔylARO8 ΔylARO9 ylARO1 ylARO2 ylARO3 ylARO4 ylARO5 scARO4^K229L^ aroG^S180F^ ylTKT* *bbxfpK acxpk ylPAR4* *ylARO10* *ylPHA2* *ylARO7* *ScARO7^G141S^::Leu* | | This work |
| YL37 | YL35 derivate; Further deletion of genes *ylALD2* and *ylALD3*; po1fk *ΔylTYR1 ΔylTRP2 ΔylTRP3 ΔylARO8 ΔylARO9 ΔylPYK ΔylALD2 ΔylADL3 ylARO1 ylARO2 ylARO3 ylARO4 ylARO5 scARO4^K229L^ aroG^S180F^ ylTKT* *bbxfpK acxpk ylPAR4* *ylARO10* *ylPHA2* *ylARO7* *ScARO7^G141S^::Leu* | | This work |
| YL38 | YL37 derivate; Further deletion of genes *ylHPD*; po1fk *ΔylTYR1 ΔylTRP2 ΔylTRP3 ΔylARO8 ΔylARO9 ΔylPYK ΔylALD2 ΔylADL3 ΔylHPD ylARO1 ylARO2 ylARO3 ylARO4 ylARO5 scARO4^K229L^ aroG^S180F^ ylTKT* *bbxfpK acxpk ylPAR4* *ylARO10* *ylPHA2* *ylARO7* *ScARO7^G141S^::Leu* | | This work |
| YL39 | YL33 derivate; Further integration of genes *rgTAL* and *ylTYR1* at pBR platform; po1fk *ΔylTYR1 ΔylTRP2 ΔylTRP3 ΔylARO8 ΔylARO9 ΔylPYK ylARO1 ylARO2 ylARO3 ylARO4 ylARO5 scARO4^K229L^ aroG^S180F^ ylTKT* *bbxfpK acxpk rgTAL ylTYR1::Leu* | | This work |
| YL40 | YL39 derivate; Further deletion of genes *ylPHA2*; po1fk *ΔylTYR1 ΔylTRP2 ΔylTRP3 ΔylARO8 ΔylARO9 ΔylPYK ΔylPHA2 ylARO1 ylARO2 ylARO3 ylARO4 ylARO5 scARO4^K229L^ aroG^S180F^ ylTKT* *bbxfpK acxpk rgTAL ylTYR1::Leu* | | This work |
| YL41 | YL33 derivate; Further integration of genes *VioA*, *VioB*, *VioC*, *VioD*, and *VioE* at pBR platform; po1fk *ΔylTYR1 ΔylTRP2 ΔylTRP3 ΔylARO8 ΔylARO9 ΔylPYK ylARO1 ylARO2 ylARO3 ylARO4 ylARO5 scARO4^K229L^ aroG^S180F^ ylTKT* *bbxfpK ylTRP2* *ylTRP3 acxpk VioA* *VioB* *VioC* *VioD VioE::Leu* | | This work |
| YL42 | YL41 derivate; Further integration of genes *ylTRP5*, *ylTRP4*, *ylTRP3*, *ylTRP2*, and *ylTRP1* at *YALI0E03212g* site; po1fk *ΔylTYR1 ΔylTRP2 ΔylTRP3 ΔylARO8 ΔylARO9 ΔylPYK ylARO1 ylARO2 ylARO3 ylARO4 ylARO5 scARO4^K229L^ aroG^S180F^ ylTKT* *bbxfpK acxpk ylTRP5* *ylTRP4 ylTRP3* *ylTRP2 ylTRP1 ylTRP2* *ylTRP3 VioA* *VioB* *VioC* *VioD VioE::Leu* | | This work |
| pof1kV | po1fk derivate; Further integration of genes *VioA*, *VioB*, *VioC*, *VioD*, and *VioE* at pBR | | This work |
| ****************************************************************************************** | | | |
| **Plasmids** | | | |
| pYLXP’ | | YaliBrick plasmid | ^3^ |
| pYLXP’*-loxP-ura* | | pYLXP’ containing the *loxP-URA-loxP* cassette | ^4^ |
| pYLXP’*-loxP-hygr* | | pYLXP’ containing the *loxP-hygr-loxP* cassette | ^4^ |
| pYLXP’-*Cre* | | pYLXP’ containing gene *Cre* | ^4^ |
| pYLXPs’ | | pYLXP’ derivate; *NotI* site was mutated to *SnaBI* site | This work |
| pURLA | | *Ku70* site integration plasmid | This work |
| pURLK | | *YALI0E30965g* site integration plasmid | This work |
| pURLD | | *YALI0E03212g* site integration plasmid | This work |
| prDNAloxP | | 26s rDNA site integration plasmid | ^4^ |
| pYLXP’-*ylPAR4* | | pYLXP’ containing gene *ylPAR4* |  |
| pYLXP’-*ylARO10* | | pYLXP’ containing gene *ylARO10* |  |
| pYLXP’-*ylPHA2* | | pYLXP’ containing gene *ylPHA2* | This work |
| pYLXP’-*ylARO7* | | pYLXP’ containing gene *ylARO7* | This work |
| pYLXP’-*ylPAR4-ylARO10-ylPHA2-ylARO7* | | pYLXP’ containing gene *ylPAR4*, *ylARO10*, *ylPHA2*, and *ylARO7* | This work |
| pYLXP’-*ylPAR4-ylARO10-ylPHA2-ylARO7-ScARO7* *^G141S^* | | pYLXP’ containing gene *ylPAR4*, *ylARO10*, *ylPHA2*, *ylARO7*, and *ScARO7* *^G141S^* | This work |
| pURLA-*ylPAR4-ylARO10-ylARO7-ylPHA2-scARO7^G141S^* | | pURLA containing gene *ylPAR4*, *ylARO10*, *ylPHA2*, *ylARO7*, and *ScARO7* *^G141S^* | This work |
| pYLXP’-*ylARO1* | | pYLXP’ containing gene *ylARO1* | ^5^ |
| pYLXP’-*ylARO2* | | pYLXP’ containing gene *ylARO2* | This work |
| pYLXP’-*ylARO3* | | pYLXP’ containing gene *ylARO3* | This work |
| pYLXP’-*ylARO4* | | pYLXP’ containing gene *ylARO4* | This work |
| pYLXP’-*ylARO5* | | pYLXP’ containing gene *ylARO5* | This work |
| pYLXP’-*ylARO1-ylARO2* | | pYLXP’ containing gene *ylARO1* and *ylARO2* | This work |
| pYLXP’-*ylARO1-ylARO2-ylARO3-ylARO4-ylARO5* | | pYLXP’ containing gene *ylARO1, ylARO2, ylARO3, ylARO4,* and *ylARO5* | This work |
| pYLXP’-*ylARO1-ylARO2-scARO4^K229L^* | | pYLXP’ containing gene *ylARO1, ylARO2, scARO4^K229L^* | This work |
| pYLXP’-*ylARO1-ylARO2-ecaroG^L175D^* | | pYLXP’ containing gene *ylARO1, ylARO2, ecaroG^L175D^* | This work |
| pYLXP’-*ylARO1-ylARO2-ecaroG^S180F^* | | pYLXP’ containing gene *ylARO1, ylARO2, ecaroG^S180F^* | This work |
| pYLXP’*-loxP-hygr-ΔylTYR1* | | pYLXP’*-loxP-hygr* containing gene *ylTYR1 deletion cassette* | This work |
| pYLXP’*-loxP-hygr-ΔylTRP2* | | pYLXP’*-loxP-hygr* containing gene *ylTRP2 deletion cassette* | This work |
| pYLXP’*-loxP-hygr-ΔylTRP3* | | pYLXP’*-loxP-hygr* containing gene *ylTRP3 deletion cassette* | This work |
| pYLXP’*-loxP-hygr-ΔylARO8* | | pYLXP’*-loxP-hygr* containing gene *ylARO8 deletion cassette* | This work |
| pYLXP’*-loxP-hygr-ΔylARO9* | | pYLXP’*-loxP-hygr* containing gene *ylARO9 deletion cassette* | This work |
| pURLA-*ylARO2-ylARO3-ylARO4-ylARO5-scARO4^K229L^-aroG^S180F^* | | pURLA containing gene *ylARO2, ylARO3, ylARO4, ylARO5, scARO4^K229L^, and aroG^S180F^* | This work |
| pURLK-*ylARO1* | | pURLK containing gene *ylARO1* | This work |
| pYLXP’-*ylTKT*-*ylPAR4-ylARO10-ylPHA2-ylARO7-ScARO7* *^G141S^* | | pYLXP’ containing gene *ylTKT*, *ylPAR4*, *ylARO10*, *ylPHA2*, and *ylARO7* | This work |
| pYLXP’-*bbxfpK*-*ylPAR4-ylARO10-ylPHA2-ylARO7-ScARO7* *^G141S^* | | pYLXP’ containing gene *bbxfpK*, *ylPAR4*, *ylARO10*, *ylPHA2*, and *ylARO7* | This work |
| pYLXP’-*acxpkA*-*ylPAR4-ylARO10-ylPHA2-ylARO7-ScARO7* *^G141S^* | | pYLXP’ containing gene *acxpkA*, *ylPAR4*, *ylARO10*, *ylPHA2*, and *ylARO7* | This work |
| prDNAloxP-*ylTKT*-*bbxfpK-acxpk* | | prDNAloxP containing gene *ylTKT*, *bbxfpK, and acxpk* | This work |
| pYLXP’*-loxP-ura-ylPYK* | | pYLXP’*-loxP-ura* containing gene *ylPYK* deletion cassette | This work |
| pYLXP’*-loxP-hygr-ylALD2* | | pYLXP’*-loxP-hygr* containing gene *ylALD2* deletion cassette | This work |
| pYLXP’*-loxP-hygr-ylALD3* | | pYLXP’*-loxP-hygr* containing gene *ylALD3* deletion cassette | This work |
| pYLXP’*-loxP-hygr-**ylHPD* | | pYLXP’*-loxP-ura* containing gene *ylHPD* deletion cassette | This work |
| pYLXP’-*rgTAL-ylTYR1* | | pYLXP’ containing gene *rgTAL* and *ylTYR1* | This work |
| pYLXP’-*rgTAL-ylTYR1-VvSTS1-Pc4CL2* | | pYLXP’ containing gene *rgTAL*, *ylTYR1*, *VvSTS1*, and *Pc4CL2* | This work |
| pYLXP’*-loxP-hygr-ylPHA2* | | pYLXP’*-loxP-hygr* containing gene *ylPHA2* deletion cassette | ^3^ |
| pYLXP’*-VioDCBAE* | | pYLXP’ containing gene *VioA, VioB, VIoC, VioD, and VioE* | This work |
| pYLXP’*-ylTRP2-ylTRP3-VioDCBAE* | | pYLXP’ containing gene *ylTRP2, ylTRP3, VioA, VioB, VIoC, VioD, and VioE* | This work |
| pYLXP’*-loxP-hygr-ylARO7* | | pYLXP’*-loxP-hygr* containing gene *ylARO7* deletion cassette | This work |

***Table 2. Primers used in this study***

| **Primers** | **Sequence** |
| --- | --- |
| TRP2_Dw-F | tagcgagacaataacggaggaTTGGAGAGTGTGAGCTCTCGTTC |
| TRP2_Dw-R | gttacatccttttatcagacataGAGTAGGAATGCTTCCGATGTACGC |
| TRP2_Up-F | ggcatccctaaatttgatgaaagATCCCATTGTTGGTTGATGCCC |
| TRP2_Up-R | taatgtatgctatacgaagttatGGTGGTGTAGTTCGGGGGTG |
| TRP3_Dw-F | gctagcgagacaataacggaggaAGGCGATGAAGATGCACTTCAT |
| TRP3_Dw-R | ttacatccttttatcagacataAATTAACAGGGTCACACGAGCTCT |
| TRP3_Up-F | gcatccctaaatttgatgaaagGCTGCCAGAGTGCATTTCTCG |
| TRP3_Up-R | taatgtatgctatacgaagttatTGTGGAGTAAGTGAAGCCGTTGAG |
| TYP1_Dw-F | gctagcgagacaataacggaggaGACACACTTGCAGGTCTAAAAGTTCC |
| TYP1_Dw-R | ttacatccttttatcagacataCCTCCGAAGAGGCTCTCAAAATGA |
| TYP1_Up-F | ggcatccctaaatttgatgaaagGGACAGAGTGTCCAACAAGCCAAT |
| TYP1_Up-R | aatgtatgctatacgaagttatcGTTGTAGAGCGTGGCGAAAGT |
| ALD3_Dw-F | gctagcgagacaataacggaggaAGGCCGTCCACATTAACCTGG |
| ALD3_Dw-R | gttacatccttttatcagacataCTGCTGCAACCAGCCCTACAAA |
| ALD3_Up-F | gcatccctaaatttgatgaaagCAAGAAGGGATAAAAATGGAAACTCGGTCT |
| ALD3_Up-R | taatgtatgctatacgaagttatACTTGCACTAGGTTAGCAGCGAC |
| ALD2_Dw-F | gctagcgagacaataacggaggaACGATGAGCGAACGAATCGTCT |
| ALD2_Dw-R | gttacatccttttatcagacataTCTGTTGGATTCTAGGGAACTGTTTCTG |
| ALD2_Up-F | ggcatccctaaatttgatgaaagCCGCTCTCAAGTGTCTGAAAGTTGAAT |
| ALD2_Up-R | taatgtatgctatacgaagttatATATTTAGAGTTCGGGATAAAGTTCAATGT |
| ALD2_Cas-F | CCGCTCTCAAGTGTCTGAAAGTTGAAT |
| ALD2_Cas-R | TCTGTTGGATTCTAGGGAACTGTTTCTG |
| TYP1_Cas-F | GGACAGAGTGTCCAACAAGCCAAT |
| TYP1_Cas-R | CTCCGAAGAGGCTCTCAAAATGA |
| TRP3_Cas-F | GCTGCCAGAGTGCATTTCTCG |
| TRp3_Cas-R | AATTAACAGGGTCACACGAGCTCT |
| TRP2_Cas-F | ATCCCATTGTTGGTTGATGCCC |
| TRP2_Cas_R | GAGTAGGAATGCTTCCGATGTACGC |
| ALD3_Cas-F | CAAGAAGGGATAAAAATGGAAACTCGGTCT |
| ALD3_Cas-R | CTGCTGCAACCAGCCCTACAAA |
| TYP1_DwChk-R | AGGATAAGAAGGCCAAGGCTTCT |
| TYP1_UpChk-F | ATGGATCTAAACGCTGGCGTCT |
| TRP2_DwChk-R | CATTGATGCACGCATCATTCCC |
| TRP2_UpChk-F | AAGTAGTAGGACAAGGGGTTGGC |
| ALD2_DwChk-R | ACTCCTCTCTAGACTCCTCCTGTTC |
| ALD2_UpChk-F | GTGTCTCCATCACATGACCACAATC |
| TRP3_DwChk-R | ACGGTAAACCTCACCTGATCCG |
| TRP3_UpChk-F | CTCTTCGACTGTTGGCTCTGTCTC |
| ALD3_DwChk-R | TAGCCTCGTTAATGCACCGAGT |
| ALD3_UpChk-F | GGGAATGCTCCATTGAGATGATGGA |
| ARO1_Ing_R(Ku70) | CCTAGtcctccgttattgtctcgggacacgggcatctcacttgc |
| ARO1_Int_F(ku70) | tagcatacattatacgaagttattgaaagcctagggacgacagagac |
| ku70-Dw-R(NotI) | acatccttttatcagacataggcggccgcAGTGAACGACCAAGACTAAAGGGTG |
| ku70_Up-F(NotI) | ggcatccctaaatttgatgaaaggcggccgcTGTACCATTCTACCCGGGGTCTG |
| PHA2_DwChkR | AGAATACCTTGATTCTGGCCACCG |
| PHA2_UpChkF | TCTGGCCGAGTTCAAGCTCCA |
| PHA2_CasF | TACCATCACCGACCCAGAGACCA |
| PHA2_CasR | TGCCGCATGCCATTCAGCTA |
| PHA2_DwF | gctagcgagacaataacggaggaCTTAGACGGTTCAGCGTTTCTGT |
| PHA2_DwR | tacatccttttatcagacataTGCCGCATGCCATTCAGCTA |
| PHA2_UpF | atccctaaatttgatgaaagTACCATCACCGACCCAGAGACCA |
| PHA2_UpR | atgtatgctatacgaagttatGATGTGTAATGTGTGTGATCAAGTGTGC |
| ARO3_F | gcactttttgcagtactaaccgcagcccgctatgcacaacgcttctaa |
| ARO3_R | aggccatggaactagtcggtaccctaacctcgtcgagtcttgacgg |
| ARO4-F | gcactttttgcagtactaaccgcagccacccaaagtcgttattaccgat |
| ARO4-R | ggccatggaactagtcggtaccctacttctgggctctcatgctgt |
| ARO5-F | cactttttgcagtactaaccgcagtcccgttcctcctctcccaac |
| ARO5-R | ggccatggaactagtcggtaccttagttcttgtttcgtcgctccttgac |
| ARO2_F | cagcactttttgcagtactaaccgcagAGCACTTTCGGCACGCTT |
| ARO2_R | acaggccatggaactagtcggtaccTTAGAAAAAGGACTTTGCGCCCTTTCT |
| pTEF-Rvs | ccggatggccagacaaagaaaca |
| XPR2-Fw | gtaaatagaaaatctggcttgtaggtggcaaaat |
| CEN1_F | atttcacagtctgaacttttgcagattacc |
| ORI1001_R | tggatctaaggttcgtactcaacactcac |
| HPD_CheckF | GCTACTTGTTGTAGACGTAGACATACACT |
| HPD_CheckR | TATTCGCACGACACTGACATTTAAGGC |
| HPD_CasF | CCGTACGTAAGAGACTGCCATAAGT |
| HPD_CasR | CTTCCCCTCCTGCTCACCCC |
| HPD_Dw_F | cgagacaataacggaggaTCATCTCTAGAGACGAGGCGTGC |
| HPD_Dw_R | agcttgcctatgttacatccttttatcagacataCTTCCCCTCCTGCTCACCCC |
| HPD_Up_F | ccctaaatttgatgaaagCCGTACGTAAGAGACTGCCATAAGT |
| HPD_Up_R | atgctatacgaagttatGTTGTTGGTGGTTATTTGTTTGTGTGTCA |
| EcAroG(L175D)_1F | ccagcactttttgcagtactaaccgcagaattatcagaacgacgatttacgcatcaaag |
| EcAroG(L175D)_2R | ggacaggccatggaactagtcggtaccttacccgcgacgcgcttttac |
| EcAroG(L175G)_1R | ctgatgcatcttcgcggtgc |
| EcAroG(L175G)_2F | gcaccgcgaagatgcatcag |
| SaARO7(G141S)_2R | aggccatggaactagtcggtaccttactcttccaaccttcttagcaagtattccac |
| ScARO7(G141S)_1F | gcactttttgcagtactaaccgcaggatttcacaaaaccagaaactgttttaaatctac |
| ScARO7(G141S)_1R | aacagaagagaagttattcttatcatcaccatctc |
| ScARO7(G141S)_2F | agagatggtgatgataagaataacttctcttctgtt |
| EcAroG(S180F)_1F | gcactttttgcagtactaaccgcagaattatcagaacgacgatttacgcatcaaag |
| EcAroG(S180F)_1R | cggacagaaaagccctgatgcca |
| EcAroG(S180F)_2F | tggcatcagggcttttctgtccg |
| EcAroG(S180F)_2R | gggacaggccatggaactagtcggtaccttacccgcgacgcgcttttac |
| ScARo4(K229L)_1F | ccagcactttttgcagtactaaccgcagagtgaatctccaatgttcgctgc |
| ScARO4(K229L)_1R | accatgcaaagtaacacccatgaaat |
| ScARo4(K229L)_2F | atttcatgggtgttactttgcatggt |
| ScARO4(K229L)_2R | ggccatggaactagtcggtaccctatttcttgttaacttctcttctttgtctgacagc |
| ylgapN_DwR(2) | acatccttttatcagacataAGCTCAATAGCCCACGACTATAGTC |
| ARO7_DwChkR | CAAAGACACCCGGCTCAGC |
| ARO7_UpChkF | CTTCTGGTTTACTCCGATACGGGGA |
| 26s rDNA_DwF | ctttaccgcagcagatccgcggccgcagatcttggtggtagtagcaaatattcaaatg |
| 26s rDNA_DwR | agatccactattggcctatcctaggccgcgggtccggctgccag |
| ylTYR1_F | cactttttgcagtactaaccgcagtctattgaggaatggaagaaaaccaagctag |
| ylTYR1_R | ggggacaggccatggaactagtcggtaccttaagtggttgacagaatggtgttgatcat |
| ARO7_CasF | CACACGCTTGTATATATATTTATCAAGTTTTTTC |
| ARO7_CasR | TCTACCGAGGCATCATGCCCAC |
| ARO7_DwF | gctagcgagacaataacggaggagtcgacAGTAGCGTGTGTTTTTTTAGCGAAGC |
| ARO7_DwR | catccttttatcagacatagcggccgcTCTACCGAGGCATCATGCCCAC |
| ARO7_UpF | ctaaatttgatgaaaggcggccgcCACACGCTTGTATATATATTTATCAAGTTTTTTC |
| ARO7_UpR | aatgtatgctatacgaagttatTTCCGGCGAATTTGGGCAGA |
| ylTRP5_F | ccagcactttttgcagtactaaccgcagtctgctcatctggctgccac |
| ylTRP5_R | cgtggggacaggccatggaactagtcggtaccttactcaaatcgcaggtcccagtc |
| scTRP1_F | agcactttttgcagtactaaccgcagtctgttattaatttcacaggtagttctggtcc |
| scTRP1_R | ggggacaggccatggaactagtcggtaccctatttcttagcatttttgacgaaatttgc |
| ylTRP2_F | accagcactttttgcagtactaaccgcaggcttccaagaccaaagttcttctgatc |
| ylTRP2_R | tggggacaggccatggaactagtcggtaccctaaccaatgagctccttgacaaaagc |
| ylTRP4_F | agcactttttgcagtactaaccgcagtctctgcctctcaacacccagc |
| ylTRP4_R | caacgtggggacaggccatggaactagtcggtaccttatgcctccttcggggccg |
| pUrLp_1F | gcatacattatacgaagttatgctagcgtccggagcggccgcGCATGCaagtcgactatgtctgataaaaggatg |
| pUrLp_1R | gcagcatcctgcgatgcagatccgaaacataatggtgcagggc |
| pUrLp_2F | gccctgcaccattatgtttcggatctgcatcgcaggatgctgc |
| pUrLp_2R | cagatccactattggcctatgggg |
| pUrLp_SeqF | accgatacgcgagcgaacg |
| pUrLp_SeqR | gggcttgtctgctcccgg |
| Ku70_DwF | gtccggagcggccgcGCATGCaagtcgacaCTAGGGAGGCACATCTAAACGAATAACG |
| Ku70_DwR | gttacatccttttatcagacatacctaggAGTGAACGACCAAGACTAAAGGGTG |
| ku70_UpF | ccctaaatttgatgaaagcctaggCGACTTGATGTTTAGAGTGTCCAGATCC |
| ku70_UpR | taatgtatgctatacgaagttatTTTCAAAAAGCGGCGGTTCGTG |
| Ace_DwF | cggccgcGCATGCaagtcgacaACACTATATAAGAATGTATTTATTTTCTCTCTTAACC |
| Ace_DwR | acatccttttatcagacatacctaggGACGCGAGTAGGTTGACTCATACA |
| Ace_UpF | gcatccctaaatttgatgaaagcctaggGCTATTCTTACGGTGTACAGTTACGAGCA |
| Ace_UpR | gtatgctatacgaagttatTGTGTGAGATGGGGTAGTACGGAA |
| Lac_DwF | gagcggccgcGCATGCaagtcgacaGTACATACGGGTATTTTGGGAAGACACA |
| Lac_DwR | catccttttatcagacatacctaggTTCCAAAACAGGGCTCCAAATGCG |
| Lac_UpF | atccctaaatttgatgaaagcctaggCTTACGACGAGCACGCTTCTGAC |
| Lac_UpR | tatgctatacgaagttatGGTCGTAGTGGTTTGTGGAGGT |
| Ace_DwChkR | CTCCTTTGTTGGCGAAGATTCCG |
| Ace_UpChkF | GTTGTTTGACGGCGTTTGACAAG |
| ylTRP1_F | accagcactttttgcagtactaaccgcaggactttctctactcttcgacatgtctacat |
| ylTRP1_R | cgtggggacaggccatggaactagtcggtaccttaccccctggcgtttttgacaaac |
| NotI/SnaBI_R | cagatccactattggcctattacgtaggatctgctgcggtaaagctc |
| NotI/SnBI_F | ggaagtcagcgccctgcaccattatgttccggatctgcatcgcagg |
| TRP3_F | ccagcactttttgcagtactaaccgcagatccaactcagacccactctagagg |
| TRP3_R | gcaacgtggggacaggccatggaactagtcggtaccctactcacccttctgctgtccag |
| Ace_DwChkR | CTCCTTTGTTGGCGAAGATTCCG |
| Ace_UpChkF | GTTGTTTGACGGCGTTTGACAAG |
| ARO34a_ChR | gtcatggatggaacaggggcca |
| aroG_ChkF | tcaaaaatggcaccgacggtacga |
| scARO7_ChkR | ctcctactcaagctttgcaaacattctat |
| ylARO7_ChkF | GATGTCATCAACAACCTGGCTCTTGATACTAA |
| RgTAL_ChkF | gtgtcgctgatcgatcaacacttc |
| VvSTS1_ChkR | ggaacttctgcagtaataatctcttgacgaatg |
| VioC_ChkR | cctggatcagcaccgacttgc |
| VioD_ChkF | ttcatgaccctgagccacgacc |
| ylTRP3_ChkF | tcggctacaacggagaggcc |
| ylTRP5_ChkR | gaggcttgcctccagtgtactttc |

***Table 3. Screening of high-performance strains***

| **Strains** | **Clone numbers** | | | | | |
| --- | --- | --- | --- | --- | --- | --- |
|  | 1 | 2 | 3 | 4 | 5 | 6 |
| **2PE (mg/L)** | | | | | | |
| YL0 | 3.12 | 1.25 | 3.52 | 4.31 | 1.28 | 2.42 |
| YL1 | 6.01 | 4.91 | 4.69 | 4.21 | 4.77 | 6.07 |
| YL2 | 16.84 | 15.01 | 7.35 | 7.10 | 9.64 | 7.02 |
| YL3 | 9.42 | 6.59 | 4.95 | 9.83 | 7.05 | 10.15 |
| YL4 | 4.95 | 2.80 | 1.96 | 7.84 | 8.61 | 4.95 |
| YL5 | 19.61 | 13.44 | 5.10 | 13.52 | 5.98 | 52.78 |
| YL6 | 45.91 | 52.52 | 8.09 | 7.27 | 5.55 | 3.57 |
| YL7 | 104.73 | 81.06 | 79.70 | 32.71 | 71.87 | 31.50 |
| YL8 | 27.91 | 17.36 | 33.02 | 31.25 | 36.47 | 36.82 |
| YL9 | 18.11 | 16.79 | 30.81 | 17.79 | 21.56 | 21.48 |
| YL10 | 77.92 | 65.63 | 28.93 | 56.42 | 32.85 | 62.82 |
| YL11 | 73.92 | 29.68 | 69.43 | 42.72 | 19.54 | 43.14 |
| YL12 | 19.72 | 35.18 | 18.27 | 47.19 | 22.55 | 33.85 |
| YL13 | 51.58 | 81.08 | 38.18 | 49.33 | 68.51 | 89.08 |
| YL14 | 50.46 | 70.93 | 51.87 | 44.12 | 14.01 | 17.48 |
| YL15 | 191.54 | 259.54 | 151.93 | 171.49 | 317.40 | 310.66 |
| YL16 | 17.21 | 15.55 | 31.19 | 57.69 | 19.64 | 15.98 |
| YL17 | 155.96 | 124.23 | 161.78 | 168.41 | 181.59 | 171.66 |
| YL21 | 80.20 | 44.95 | 25.87 | 46.62 | 49.24 | 55.69 |
| YL22 | 299.98 | 328.41 | 317.85 | 310.65 | 280.67 | 323.90 |
| YL23 | 19.00 | 400.26 | 340.45 | 411.12 | 417.22 | 360.08 |
| YL25 | 290.53 | 470.77 | 540.94 | 283.13 | 297.29 | 366.72 |
| YL26 | 436.87 | 336.87 | 334.81 | 302.29 | 389.76 | 377.62 |
| YL27 | 100.08 | 70.98 | 154.52 | 128.54 | 63.19 | 175.35 |
| YL28 | 90.79 | 18.46 | 47.54 | 116.80 | 66.47 | 166.55 |
| YL29 | 368.29 | 61.30 | 52.03 | 125.03 | 33.88 | 57.65 |
| YL30 | 402.60 | 282.74 | 278.75 | 609.87 | 353.52 | 311.73 |
| YL32 | 371.70 | 11.87 | 205.22 | 282.22 | 298.26 | 140.18 |
| YL34 | 36.23 | 64.83 | 9.48 | 37.47 | 16.25 | 11.91 |
| YL35 | 452.16 | 796.15 | 620.45 | 771.47 | 313.86 | 605.25 |
| YL36 | 163.99 | 492.29 | 454.35 | 201.74 | 380.01 | 609.48 |
| YL37 | 667.31 | 1156.35 | 853.28 | 418.57 | 247.00 | 768.40 |
| YL38 | 741.23 | 102.09 | 939.73 | 1156.93 | 577.68 | 499.90 |
| ***p*-Coumaric acid (mg/L)** | | | | | | |
| YL39 | 61.23 | 121.14 | 153.39 | 218.98 | 81.33 | 214.99 |
| YL40 | 21.27 | 6.28 | 4.93 | 189.55 | 236.40 | 323.41 |
| **Resveratrol (mg/L)** | | | | | | |
| YLRes | 8.39 | 3.54 | 2.36 | 7.31 | 5.93 | 2.54 |
| Violacein (mg/L) | | | | | | |
| YL41 | 69.18 | 59.85 | 169.91 | 15.48 | 53.98 | 202.04 |
| YL42 | 121.93 | 61.80 | 56.75 | 157.30 | 54.61 | 291.17 |
| po1fkV | 89.87 | 65.14 | 71 | 62 | 77.22 | 95.11 |

Data in red color indicates the highest production titer screened from the pooled yeast colonies.

***Table 4*** The comparison of 2PE and other aromatic production by *de novo* pathway in microorganisms

| Strains | Substrate | Titer | Cultivation | Reference |
| --- | --- | --- | --- | --- |
| **2-PE production by *de novo* pathway in microorganisms** | | | | |
| *S. cerevisiae* BY4741 | glucose | 0.10 g/L | Shake flask | ^6^ |
| *S. cerevisiae* BY4741 | glucose | 0.41 g/L | Shake flask | ^7^ |
| *S. cerevisiae* | glucose | 1.59 g/L | 2-L Bioreactor | ^8^ |
| *E. coli* DG02 | glucose | 1.02 g/L | Shake flask | ^9^ |
| *E. coli* NST74 | glucose | 1.94 g/L | Shake flask | ^10^ |
| *E. coli* MG1655 | glucose | 0.18 g/L | Shake flask | ^11^ |
| *Enterobacter sp.* CGMCC 508 | glucose | 0.34 g/L | Shake flask | ^12^ |
| *E. coli* BP-42 | glucose | 0.29 g/L | Shake flask | ^13^ |
| *E. coli* BW25113(DE3) | glucose | 0.94 g/L | Shake flask | ^14^ |
| **YL25** | glucose | 2.43 g/L | Shake flask | This work |
| ***p*-coumaric acid production by *de novo* pathway in microorganisms** | | | | |
| *S. cerevisiae* | glucose | 12.5 g/L | 1-L Bioreactor | ^15^ |
| *E. coli* | glucose | 0.97 g/L | Shake flask | ^16^ |
| *S. cerevisiae* W303-1A | glucose | 0.01 g/L | Shake flask | ^17^ |
| *S. cerevisiae* | glucose | 1.93 g/L | 96-deep well plate | ^18^ |
| **YL40** | glucose | 0.59 g/L | Shake flask | This work |
| **Violacein production by de novo pathway in microorganisms** | | | | |
| *Janthinobacterium lividum* | glycerol | 1.83 g/L | 2-L Bioreactor | ^19^ |
| *Chromobacterium violaceum* | - | 0.15 g/L | Shake flask | ^20^ |
| *Y. lipolytica* | glucose | 0.07 g/L | Shake flask | ^21^ |
| **YL42** | glucose | 0.37 g/L | Shake flask | This work |

### **Supplementary Methods**

**Integrative plasmid constructed in this work**

Compared to *S. cerevisiae*, the genetic toolbox in *Y. lipolytica* is still less developed, due to the unclear genetic backgrounds and complexity of the non-homologous end joining mechanism ^5^. To overcome these limitations, we constructed three genomic integration plasmids, namely pUrLA, pUrLK, and pHyLD, to assemble and deposit very long gene fragments to the chromosome. Plasdmid maps and sequence for pUrLA, pUrLK, and pHyLD have been appended in this supplementary file.

### **Supplementary Notes**

***Note1. Deduction of the stoichiometry of 2PE biosynthesis and the theoretical yield calculation***

For deducing the stoichiometry of 2PE biosynthesis by *de novo* pathway from glucose, we divided the 2PE synthesis pathway to three parts, including (i) glucose to PEP; (ii) glucose to E4P; and (iii) PEP and E4P to 2PE. According to the reactions annotated by KEGG (https://www.kegg.jp/), we got three stoichiometry equations:

(i) Glucose to PEP: Glc 🡺 2 PEP + NADH

(ii) Glucose to E4P: 1.5 Glc + ATP 🡺 2 E4P +2 NAD(P)H + CO_2_

(iii) PEP and E4P to 2PE: 2 PEP + E4P + NADPH + NADH + ATP 🡺 2 CO_2_ + 2PE

Thus, the stoichiometry of 2PE biosynthesis by De Novo pathway from glucose is (3.5 Glc + 3 ATP 🡺 5 CO_2_ + 2 2PE).

Subsequently, we established a global stoichiometric model of 2PE biosynthesis and assumed that production of 1 mol of 2PE needs x mol of glucose. Thus, the overall stoichiometrics will be:

x Glucose (C_6_H_12_O_6_) + (6x-10) O_2_ 🡺 2PE (C_8_H_10_O) + (6x-8) CO_2_ + (6x-5) H_2_O

Thus, the yield of 2-PE (g/g _glucose_) is

Y*_2-PE_*=$\frac{122}{180x}$ (1)

Furthermore, to assess the carbon conversion efficiency, we introduced the respiratory quotient (RQ):

RQ=$\frac{6x-8}{6x-10}$ (2)

Thus, the yield of 2-PE could be solved as

Y*_2-PE_*=$\frac{122}{180x}$=$\frac{122}{180}$*$\frac{3n-3}{5n-4}$(3)

As Shown in the metabolic model (SFigure1), the stoichiometrics (Eqn. 3) suggest that the theoretically maximum Y*_2-PE_* is 0.4436 g/g _glucose_.

### **Supplementary Figures**

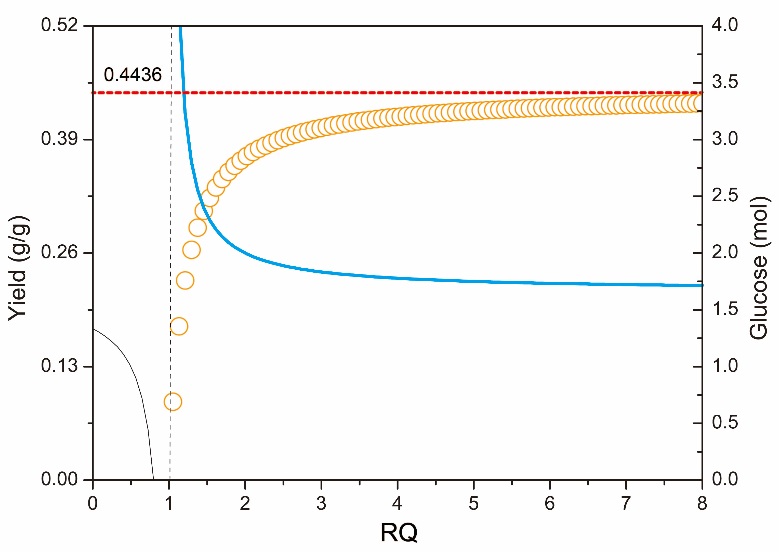

**Figure S1. Mathematical models of 2-PE yield**

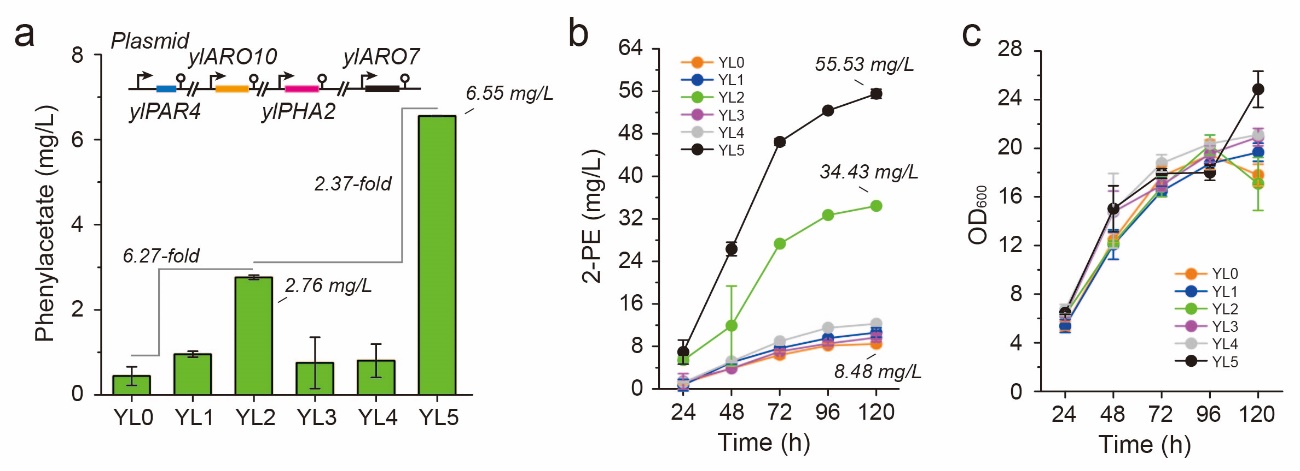

**Figure S2.** (a-c) Phenylacetate titer, time profiles of 2-PE titer and cell growth of strains carrying the 2-PE pathway, including genes *ylPAR4* (*YALI0D07062g*, encoding phenylacetaldehyde reductase), *ylARO10* (*YALI0D06930g*, encoding phenylpyruvate decarboxylase), *ylPHA2* (*YALI0B17336g*, encoding prephenate dehydratase) and *ylARO7* (*YALI0E17479g*, encoding chorismate mutase). All experiments were performed in triplicate and error bars represent standard deviations (SD).

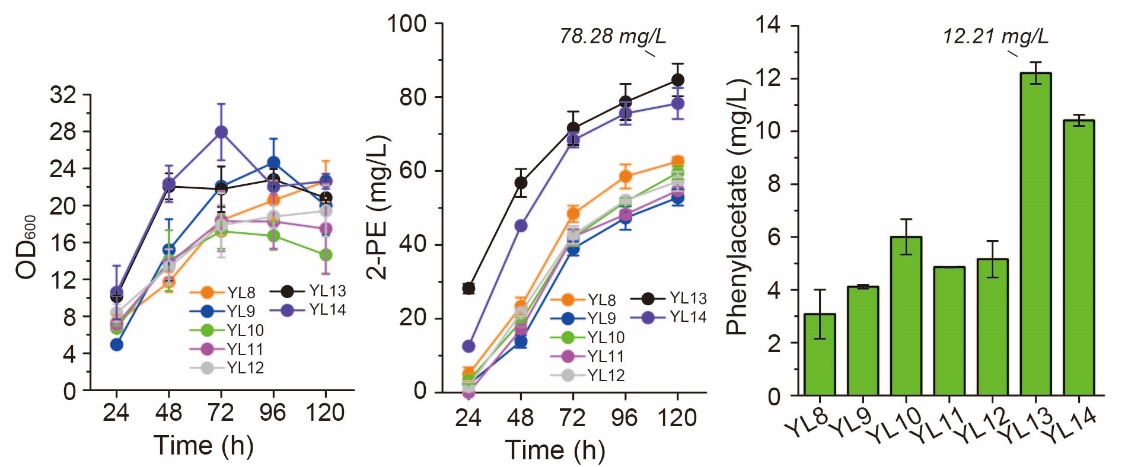

**Figure S3.** Phenylacetate, 2-PE and cell growth profile of strains overexpressing genes *ylARO1* (*YALI0F12639g*, encoding pentafunctional protein), *ylARO2* (*YALI0D17930g*, encoding bifunctional chorismate synthase), *ylARO3* (*YALI0B20020g*, encoding DAHP synthase), *ylARO4* (*YALI0B22440g*, encoding DAHP synthase), and *ylARO5* (*YALI0C06952g*, encoding DAHP synthase); All experiments were performed in triplicate and error bars represent standard deviations (SD).

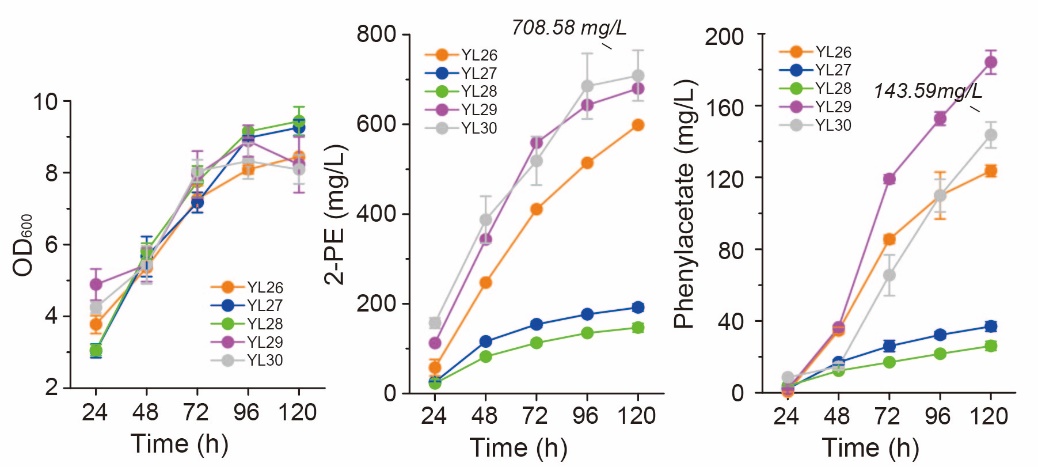

**Figure S4.** Time profiles of 2-PE, cell growth, and phenylacetate of strains overexpressing transketolase *ylTKT*, phosphoketolases *BbxfpK* and *AcxpkA*; All experiments were performed in triplicate and error bars represent standard deviations (SD).

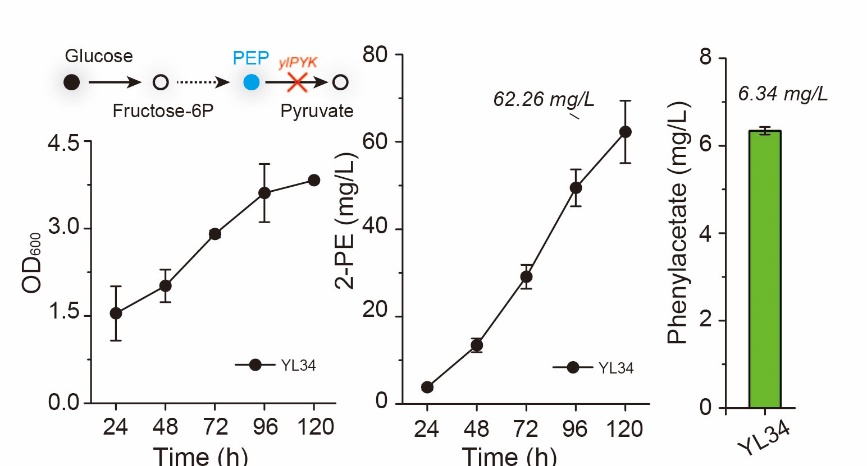

**Figure S5.** Time profile of 2-PE, cell growth, and phenylacetate of strains with pyruvate kinase ylPYK deletion in CSM medium with feeding of acetate. All experiments were performed in triplicate and error bars represent standard deviations (SD).

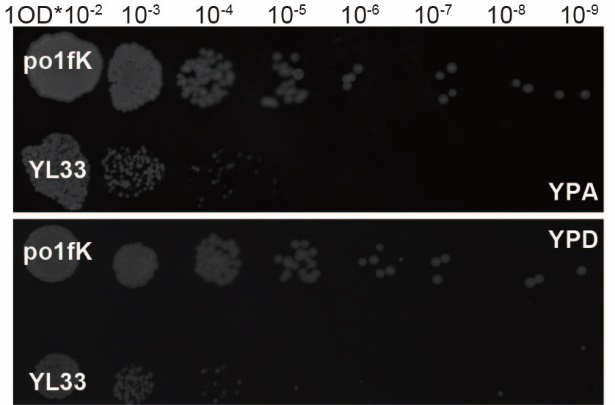

**Figure S6. Cell growth on plate (24h) of *Y. lipolytica* with gene *ylPYK* deletion.** po1fK, po1fk *Δku70::loxP*; YL33, po1fk *ΔylTYR1 ΔylTRP2 ΔylTRP3 ΔylARO8 ΔylARO9 ΔylPYK ylARO1 ylARO2 ylARO3 ylARO4 ylARO5 scARO4K229L aroGS180F ylTKT bbxfpK acxpk::loxP*; YPD medium, containing glucose 40.0 g/L, yeast extract 10.0 g/L, peptone 20.0 g/L; YPA medium, YPD with sodium acetate 5.0 g/L.

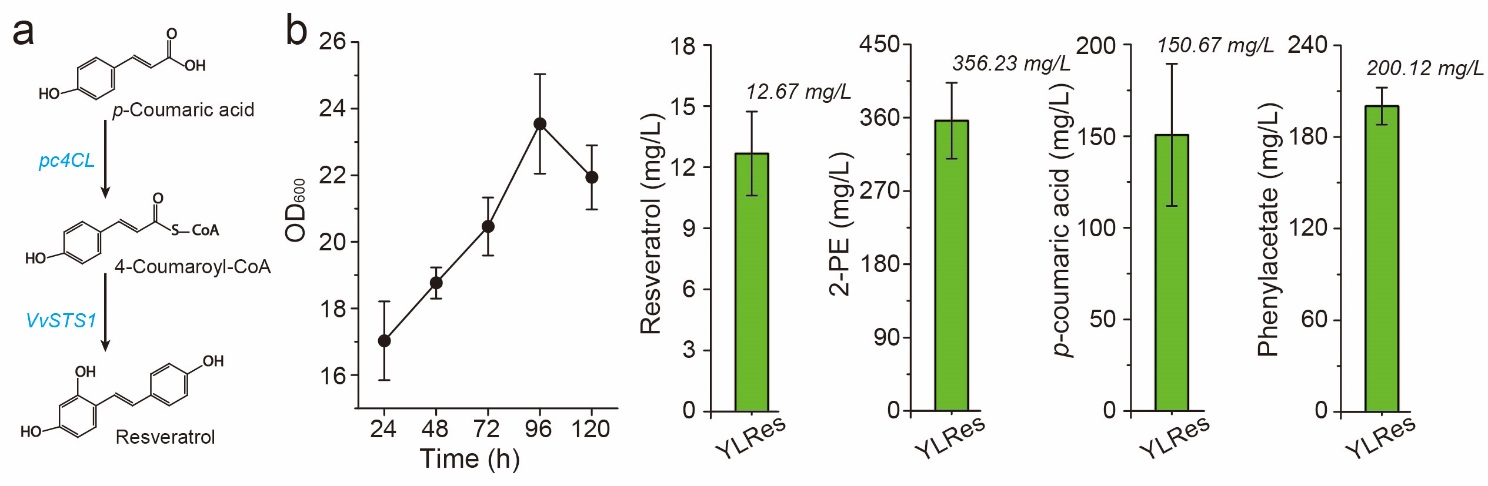

**Figure S7. Resveratrol production by strain YLRes.** (a) The catalytic steps form *p*-coumaric acid to resveratrol; pc4CL, 4-coumarate-CoA ligase from *Petroselinum crispum*; VvSTS1, resveratrol synthase from *Vitis vinifera*; (b) Cell growth, resveratrol titer, 2-PE titer, *p*-coumaric acid titer and phenylacetate titer of strain YLRes. All experiments were performed in triplicate and error bars show standard deviation (SD). All experiments were performed in triplicate and error bars represent standard deviations (SD).

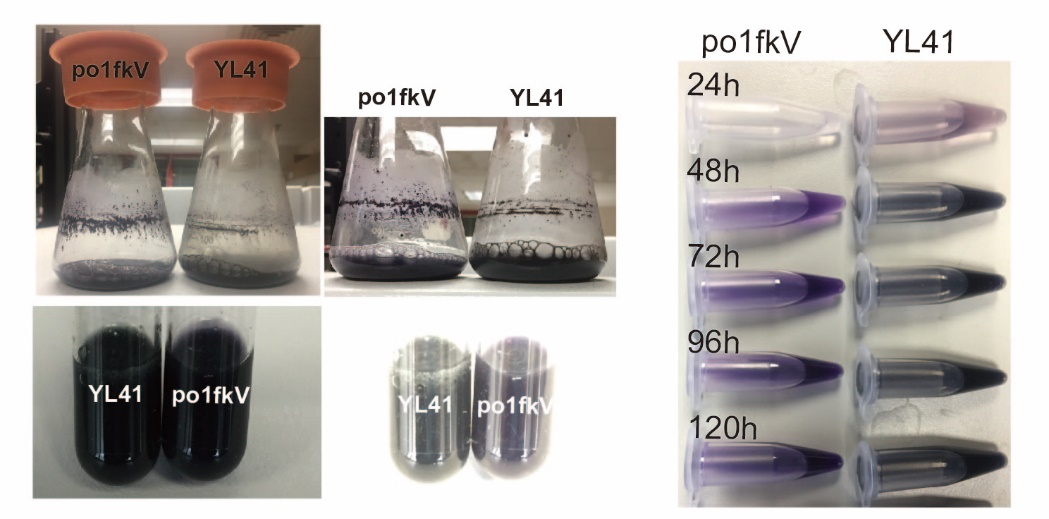

**Figure S8. Shaking flask cultivation and extraction of violacein produced by strains YL41 and po1fkV.** (a) Cell cultural of shaking flask cultivation. The extraction process is that 0.20 mL of fermentation culture was mixed with 5-fold volume of ethyl acetate and appropriate glass beads, vortexed at 30 ^o^C for 24 h.

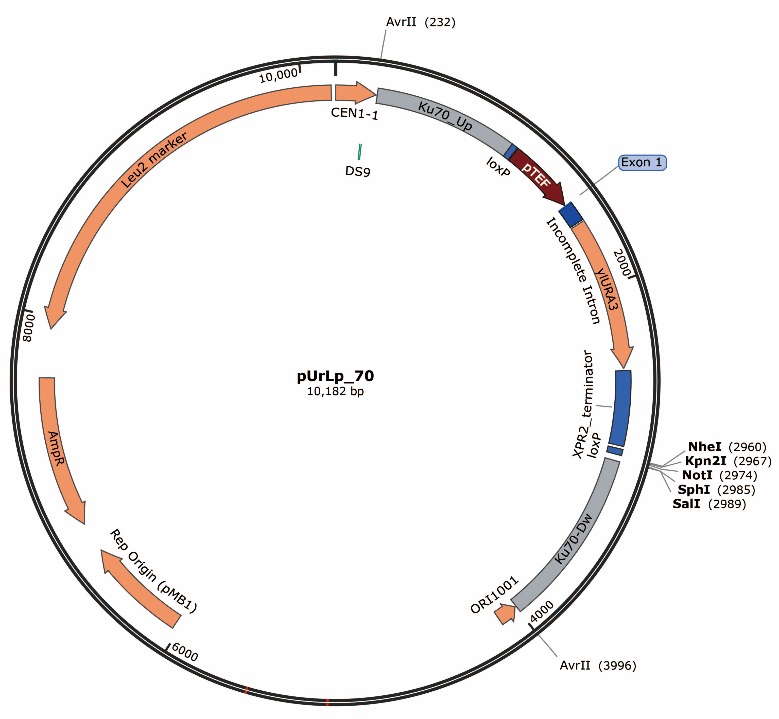

**Figure S9. Plasmid pUrLp70 map.** The multiple clone sites include *NheI, Kpn2I, NotI, SphI*, and *SalI*. After assembling the desired genes, the integration cassettes could be obtained by digested plasmid with enzyme *AvrII*.

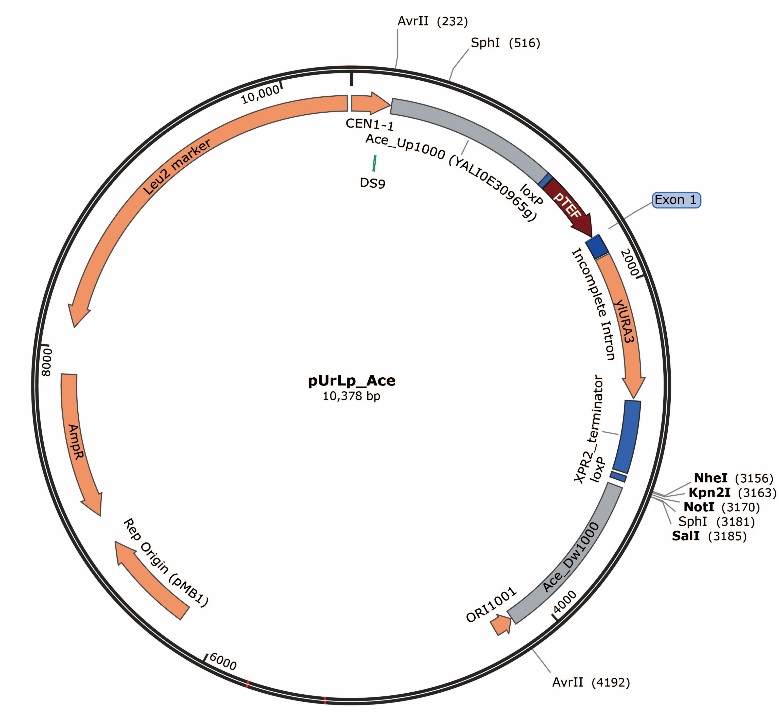

**Figure S10. Plasmid pUrLAce map.** The multiple clone sites include *NheI, Kpn2I, NotI,* and *SalI*. After assembling the desired genes, the integration cassettes could be obtained by digested plasmid with enzyme *AvrII*.

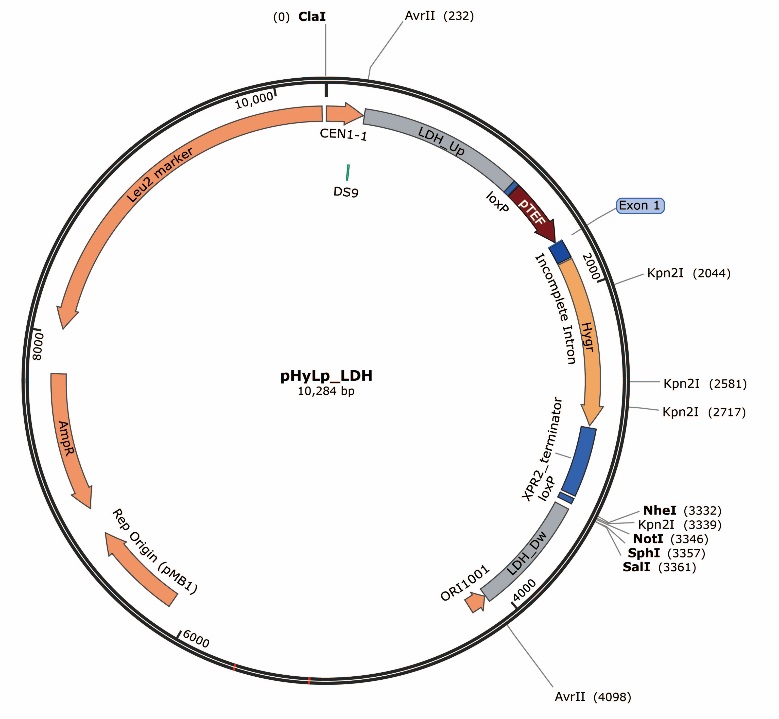

**Figure S11. Plasmid pUrLDH map.** The multiple clone sites include *NheI, NotI, SphI*, and *SalI*. After assembling the desired genes, the integration cassettes could be obtained by digested plasmid with enzyme *AvrII*.

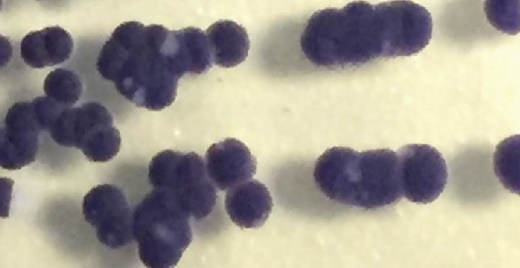

**Figure S12.** Yeast colonies (po0fkV) with integrating violacein pathway in genome

### **DNA Sequences**

Plasmid pUrLA sequences:

1	CGATGCTTTT CGTAGATAAT GGAATACAAA TGGATATCCA GAGTATACAC ATGGATAGTA

61	TACACTGACA CGACAATTCT GTATCTCTTT ATGTTAACTA CTGTGAGGCG TTAAATAGAG

121	CTTGATATAT AAAATGTTAC ATTTCACAGT CTGAACTTTT GCAGATTACC TAATTTGGTA

181	AGATATTAAT TATGAACTGA AAGTTGATGG CATCCCTAAA TTTGATGAAA GCCTAGGGAG

241	AACTGCTCCT GTGAATCTCT TAACGAACAC AGTCGCTCAA CCAATGTTGG TGATGAAGAT

301	GTCAAAAACA ACGCGGAGCC TACCCCCGAA GTGGGGGGTC AAACCTGAGA CAGATCGCGT

361	ACCGCCTGCA TACCTGCAAT GCAACCATCA TGCTACTTGT AGTGTTGCAG GGCCCGATGT

421	TCCACCGAAG CATTTCATTG GTGGATCACC CACTAGTTTG AACTGGTATG ATCTCTGTTT

481	TTGTTTTGTC AATTGCATAA CCATGTTGGA GGGCCATGTT TATTTACCCC CCACGCCCCT

541	GCTCTCACAG CTATTTTTCA GCCCGTGTCT CACACCGTCG GGGTGGTTTT AGTTTGTCAT

601	TCAATTACTG CTGACCGGCG TGTTCGTCCC CACTCGCTAC TCAACACACC CACCGCCACT

661	GCTACACTGC GCCACTAGTC TGACGGATTT GCTTGCATTC CTCCAATATG TAGCTTACAA

721	CACGTTTTTA AGCGCCGTCA CATATAATTA ATTTAATCTG CATTTTTCTA TCTCTCGCTT

781	GCCCGTAGTA TTACGTGAAT CAACTAGAAC ATATGGGGAG CTTCTGTTGC TGTTTCTCCA

841	ACTGCAATTA TGTCTACTAC AAGTAGTATA TATTTGACCA ACCGGTTATT GAACCCTACT

901	GGCTGTATTC TGGGGGAATA CATTGCATAG ATATGCACGT AAGTGGGGGT GTCTTTCTCG

961	CGCCATTGTA CTAAGAATGA CCCCCACGTC TCATTCTGGG GCAACCAGAG TCACAGCGAA

1021	GGGATATATA GCTCAAGCTA GTCTTTAATA CACTTCCTCT TTTCTGACAT TTGATCTTTC

1081	ACAACCGTCC TCGCTGAGCG CTTTTGACTT ACTGTAAGGG ACTCCTCTCT GTGCAACTCT

1141	ATTACTCACT CCGCAAGCCA CGAATCTACA ACTACCGATA CTCTGAATTG GCATAGGGTC

1201	TCTTTCCATT CTTGATACAT TCACACATCC AGTTTCGCCA TAACTTCGTA TAGCATACAT

1261	TATACGAAGT TATGACGACA GAGACCGGGT TGGCGGCGCA TTTGTGTCCC AAAAAACAGC

1321	CCCAATTGCC CCAATTGACC CCAAATTGAC CCAGTAGCGG GCCCAACCCC GGCGAGAGCC

1381	CCCTTCTCCC CACATATCAA ACCTCCCCCG GTTCCCACAC TTGCCGTTAA GGGCGTAGGG

1441	TACTGCAGTC TGGAATCTAC GCTTGTTCAG ACTTTGTACT TGTTTCTTTG TCTGGCCATC

1501	CGGGTAACCC ATGCCGGACG CAAAATAGAC TACTGAAAAT TTTTTTGCTT TGTGGTTGGG

1561	ACTTTAGCCA AGGGTATAAA AGACCACCGT CCCCGAATTA CCTTTCCTCT TCTTTTCTCT

1621	CTCTCCTTGT CAACTCACAC CCGAAATCGT TAAGCATTTC CTTCTGAGTA TAAGAATCAT

1681	TCAAATCTAG AATGGTGAGT TTCAGAGGCA GCAGCAATTG CCACGGGCTT TGAGCACACG

1741	GCCGGGTGTG GTCCCATTCC CATCGACACA AGACGCCACG TCATCCGACC AGCACTTTTT

1801	GCAGTACTAA CCGCAGTAAC CGCAGAAAAA GCCTGAACTC ACCGCCACGT CTGTCGAGAA

1861	GTTTCTGATC GAAAAGTTCG ACAGCGTCTC CGACCTGATG CAGCTCTCGG AGGGCGAAGA

1921	ATCTCGTGCT TTCAGCTTCG ATGTAGGAGG GCGTGGATAT GTCCTGCGGG TAAATAGCTG

1981	CGCCGATGGT TTCTACAAAG ATCGTTATGT TTATCGGCAC TTTGCATCGG CCGCGCTCCC

2041	GATTCCGGAA GTGCTTGACA TTGGGGAATT TAGCGAGAGC CTGACCTATT GCATCTCCCG

2101	CCGTGCACAG GGTGTCACGT TGCAAGACCT GCCTGAAACC GAACTGCCCG CTGTTCTGCA

2161	GCCGGTCGCG GAGGCCATGG ATGCGATCGC TGCGGCCGAT CTTAGCCAGA CGAGCGGGTT

2221	CGGCCCATTC GGACCGCAAG GAATCGGTCA ATACACTACA TGGCGTGATT TCATATGCGC

2281	GATTGCTGAT CCCCATGTGT ATCACTGGCA AACTGTGATG GACGACACCG TCAGTGCGTC

2341	CGTCGCGCAG GCTCTCGATG AGCTGATGCT TTGGGCCGAG GACTGCCCCG AAGTCCGGCA

2401	CCTCGTGCAC GCGGATTTCG GCTCCAACAA TGTCCTGACG GACAATGGCC GCATAACAGC

2461	GGTCATTGAC TGGAGCGAGG CGATGTTCGG GGATTCCCAA TACGAGGTCG CCAACATCTT

2521	CTTCTGGAGG CCGTGGTTGG CTTGTATGGA GCAGCAGACG CGCTACTTCG AGCGGAGGCA

2581	TCCGGAGCTT GCAGGATCGC CGCGGCTCCG GGCGTATATG CTCCGCATTG GTCTTGACCA

2641	ACTCTATCAG AGCTTGGTTG ACGGCAATTT CGATGATGCA GCTTGGGCGC AGGGTCGATG

2701	CGACGCAATC GTCCGATCCG GAGCCGGGAC TGTCGGGCGT ACACAAATCG CCCGCAGAAG

2761	CGCGGCCGTC TGGACCGATG GCTGTGTAGA AGTACTCGCC GATAGTGGAA ACCGACGCCC

2821	CAGCACTCGT CCGAGGGCAA AGGAATAGGG TACCGACTAG TTCCATGGCC TGTCCCCACG

2881	TTGCCGGTCT TGCCTCCTAC TACCTGTCCA TCAATGACGA GGTTCTCACC CCTGCCCAGG

2941	TCGAGGCTCT TATTACTGAG TCCAACACCG GTGTTCTTCC CACCACCAAC CTCAAGGGCT

3001	CTCCCAACGC TGTTGCCTAC AACGGTGTTG GCATTTAGGC AATTAACAGA TAGTTTGCCG

3061	GTGATAATTC TCTTAACCTC CCACACTCCT TTGACATAAC GATTTATGTA ACGAAACTGA

3121	AATTTGACCA GATATTGTTG TAAATAGAAA ATCTGGCTTG TAGGTGGCAA AATGCGGCGT

3181	CTTTGTTCAT CAATTCCCTC TGTGACTACT CGTCATCCCT TTATGTTCGA CTGTCGTATT

3241	TCTTATTTTC CATACATATG CAAGTGAGAT GCCCGTGTCC GTTATCAAAT CTAGTTAATA

3301	ACTTCGTATA GCATACATTA TACGAAGTTA TGCTAGCGTC CGGAGCGGCC GCGCATGCAA

3361	GTCGACAACC TGCAAAGTCT CTCACGTACG TCTGTAATGA TACAAAATGT AAGCCTTGAA

3421	ATGTATGGGC TGCCTGTATT CGTAAAATTA CAGTACTTGT ACGCCATATA TGGGGTCCAT

3481	ATGTTGTTTA ACTATGTCAC CAATTCTGAC AGTCATTCGA TCTAATTATG CAATTTGTAT

3541	TCAAAGGGCA TGACTCCCTC TTTTCACTCA CTATTGTTTT TTATTATTAT TATATTTTTA

3601	TAATTTGATT GCCATCCTGT CTGCACCTAT TCCAACTACT GTATGTACTG TTCAAGTGCT

3661	ACAAGTAGTA CGATACTTGC ACTGTAACTA CAAGTCCAAG TAGGTTTTGT GAGCTAAATT

3721	GGACTCTATC CATAGACGAG TTTGTTGAGC GGCTATCGAC AGGAAAGTGT TACAGTGATG

3781	TACTTGTATG TACGTGTATC GTACTGTTTA GTAAACATAT ATATAGACAA CTTGACACAC

3841	CCAAATCAAT AGGCAATGAG GCTAGTTTCC ATTTATACAT CTCCGTGACC TTTGATTGTC

3901	ATGTTGGACT GATAAGATGT GCACATTGAC GCGCCATAAA TGGATACTGG GTCCGAGATT

3961	TATGCATCCC CCACATGTAT AGTCAACCTT AAGGGCAGGT CACATCTCCC ATCACCATCA

4021	AATCTGACCA CCAAAACAAT TGGATCTGAG CGATGATCCT CATCAAAAAT AAAGTGGCGC

4081	TGAAGCTGCT ACAGCTACCT AGGTATGTCT GATAAAAGGA TGTAACATAG GCAAGCTGCT

4141	CGTGAGTGTT GAGTACGAAC CTTAGATCCA AATCACCCGC ACCCACGGAT ATACTTGCTT

4201	GAATATACAG TAGTATGCTC GACCGATGCC CTTGAGAGCC TTCAACCCAG TCAGCTCCTT

4261	CCGGTGGGCG CGGGGCATGA CTATCGTCGC CGCACTTATG ACTGTCTTCT TTATCATGCA

4321	ACTCGTAGGA CAGGTGCCGG CAGCGCTCTG GGTCATTTTC GGCGAGGACC GCTTTCGCTG

4381	GAGCGCGACG ATGATCGGCC TGTCGCTTGC GGTATTCGGA ATCTTGCACG CCCTCGCTCA

4441	AGCCTTCGTC ACTGGTCCCG CCACCAAACG TTTCGGCGAG AAGCAGGCCA TTATCGCCGG

4501	CATGGCGGCC GACGCGCTGG GCTACGTCTT GCTGGCGTTC GCGACGCGAG GCTGGATGGC

4561	CTTCCCCATT ATGATTCTTC TCGCTTCCGG CGGCATCGGG ATGCCCGCGT TGCAGGCCAT

4621	GCTGTCCAGG CAGGTAGATG ACGACCATCA GGGACAGCTT CAAGGATCGC TCGCGGCTCT

4681	TACCAGCCTA ACTTCGATCA CTGGACCGCT GATCGTCACG GCGATTTATG CCGCCTCGGC

4741	GAGCACATGG AACGGGTTGG CATGGATTGT AGGCGCCGCC CTATACCTTG TCTGCCTCCC

4801	CGCGTTGCGT CGCGGTGCAT GGAGCCGGGC CACCTCGACC TGAATGGAAG CCGGCGGCAC

4861	CTCGCTAACG GATTCACCAC TCCAAGAATT GGAGCCAATC AATTCTTGCG GAGAACTGTG

4921	AATGCGCAAA CCAACCCTTG GCAGAACATA TCCATCGCGT CCGCCATCTC CAGCAGCCGC

4981	ACGCGGCGCA TCTCGGGCAG CGTTGGGTCC TGGCCACGGG TGCGCATGAT CGTGCTCCTG

5041	TCGTTGAGGA CCCGGCTAGG CTGGCGGGGT TGCCTTACTG GTTAGCAGAA TGAATCACCG

5101	ATACGCGAGC GAACGTGAAG CGACTGCTGC TGCAAAACGT CTGCGACCTG AGCAACAACA

5161	TGAATGGTCT TCGGTTTCCG TGTTTCGTAA AGTCTGGAAA CGCGGAAGTC AGCGCCCTGC

5221	ACCATTATGT TTCGGATCTG CATCGCAGGA TGCTGCTGGC TACCCTGTGG AACACCTACA

5281	TCTGTATTAA CGAAGCGCTG GCATTGACCC TGAGTGATTT TTCTCTGGTC CCGCCGCATC

5341	CATACCGCCA GTTGTTTACC CTCACAACGT TCCAGTAACC GGGCATGTTC ATCATCAGTA

5401	ACCCGTATCG TGAGCATCCT CTCTCGTTTC ATCGGTATCA TTACCCCCAT GAACAGAAAT

5461	CCCCCTTACA CGGAGGCATC AGTGACCAAA CAGGAAAAAA CCGCCCTTAA CATGGCCCGC

5521	TTTATCAGAA GCCAGACATT AACGCTTCTG GAGAAACTCA ACGAGCTGGA CGCGGATGAA

5581	CAGGCAGACA TCTGTGAATC GCTTCACGAC CACGCTGATG AGCTTTACCG CAGCAGATCC

5641	GCGGCCCCAT AGGCCAATAG TGGATCTGCT GCCTCGCGCG TTTCGGTGAT GACGGTGAAA

5701	ACCTCTGACA CATGCAGCTC CCGGAGACGG TCACAGCTTG TCTGTAAGCG GATGCCGGGA

5761	GCAGACAAGC CCGTCAGGGC GCGTCAGCGG GTGTTGGCGG GTGTCGGGGC GCAGCCATGA

5821	CCCAGTCACG TAGCGATAGC GGAGTGTATA CTGGCTTAAC TATGCGGCAT CAGAGCAGAT

5881	TGTACTGAGA GTGCACCATA TGCGGTGTGA AATACCGCAC AGATGCGTAA GGAGAAAATA

5941	CCGCATCAGG CGCTCTTCCG CTTCCTCGCT CACTGACTCG CTGCGCTCGG TCGTTCGGCT

6001	GCGGCGAGCG GTATCAGCTC ACTCAAAGGC GGTAATACGG TTATCCACAG AATCAGGGGA

6061	TAACGCAGGA AAGAACATGT GAGCAAAAGG CCAGCAAAAG GCCAGGAACC GTAAAAAGGC

6121	CGCGTTGCTG GCGTTTTTCC ATAGGCTCCG CCCCCCTGAC GAGCATCACA AAAATCGACG

6181	CTCAAGTCAG AGGTGGCGAA ACCCGACAGG ACTATAAAGA TACCAGGCGT TTCCCCCTGG

6241	AAGCTCCCTC GTGCGCTCTC CTGTTCCGAC CCTGCCGCTT ACCGGATACC TGTCCGCCTT

6301	TCTCCCTTCG GGAAGCGTGG CGCTTTCTCA TAGCTCACGC TGTAGGTATC TCAGTTCGGT

6361	GTAGGTCGTT CGCTCCAAGC TGGGCTGTGT GCACGAACCC CCCGTTCAGC CCGACCGCTG

6421	CGCCTTATCC GGTAACTATC GTCTTGAGTC CAACCCGGTA AGACACGACT TATCGCCACT

6481	GGCAGCAGCC ACTGGTAACA GGATTAGCAG AGCGAGGTAT GTAGGCGGTG CTACAGAGTT

6541	CTTGAAGTGG TGGCCTAACT ACGGCTACAC TAGAAGGACA GTATTTGGTA TCTGCGCTCT

6601	GCTGAAGCCA GTTACCTTCG GAAAAAGAGT TGGTAGCTCT TGATCCGGCA AACAAACCAC

6661	CGCTGGTAGC GGTGGTTTTT TTGTTTGCAA GCAGCAGATT ACGCGCAGAA AAAAAGGATC

6721	TCAAGAAGAT CCTTTGATCT TTTCTACGGG GTCTGACGCT CAGTGGAACG AAAACTCACG

6781	TTAAGGGATT TTGGTCATGA GATTATCAAA AAGGATCTTC ACCTAGATCC TTTTAAATTA

6841	AAAATGAAGT TTTAAATCAA TCTAAAGTAT ATATGAGTAA ACTTGGTCTG ACAGTTACCA

6901	ATGCTTAATC AGTGAGGCAC CTATCTCAGC GATCTGTCTA TTTCGTTCAT CCATAGTTGC

6961	CTGACTCCCC GTCGTGTAGA TAACTACGAT ACGGGAGGGC TTACCATCTG GCCCCAGTGC

7021	TGCAATGATA CCGCGAGACC CACGCTCACC GGCTCCAGAT TTATCAGCAA TAAACCAGCC

7081	AGCCGGAAGG GCCGAGCGCA GAAGTGGTCC TGCAACTTTA TCCGCCTCCA TCCAGTCTAT

7141	TAATTGTTGC CGGGAAGCTA GAGTAAGTAG TTCGCCAGTT AATAGTTTGC GCAACGTTGT

7201	TGCCATTGCT GCAGGCATCG TGGTGTCACG CTCGTCGTTT GGTATGGCTT CATTCAGCTC

7261	CGGTTCCCAA CGATCAAGGC GAGTTACATG ATCCCCCATG TTGTGCAAAA AAGCGGTTAG

7321	CTCCTTCGGT CCTCCGATCG TTGTCAGAAG TAAGTTGGCC GCAGTGTTAT CACTCATGGT

7381	TATGGCAGCA CTGCATAATT CTCTTACTGT CATGCCATCC GTAAGATGCT TTTCTGTGAC

7441	TGGTGAGTAC TCAACCAAGT CATTCTGAGA ATAGTGTATG CGGCGACCGA GTTGCTCTTG

7501	CCCGGCGTCA ACACGGGATA ATACCGCGCC ACATAGCAGA ACTTTAAAAG TGCTCATCAT

7561	TGGAAAACGT TCTTCGGGGC GAAAACTCTC AAGGATCTTA CCGCTGTTGA GATCCAGTTC

7621	GATGTAACCC ACTCGTGCAC CCAACTGATC TTCAGCATCT TTTACTTTCA CCAGCGTTTC

7681	TGGGTGAGCA AAAACAGGAA GGCAAAATGC CGCAAAAAAG GGAATAAGGG CGACACGGAA

7741	ATGTTGAATA CTCATACTCT TCCTTTTTCA ATATTATTGA AGCATTTATC AGGGTTATTG

7801	TCTCATGAGC GGATACATAT TTGAATGTAT TTAGAAAAAT AAACAAATAG GGGTTCCGCG

7861	CACATTTCCC CGAAAAGTGC CACCTGACGT CTAAGAAACC ATTATTATCA TGACATTAAC

7921	CTATAAAAAT AGGCGTATCA CGAGGCCCTT TCGTCTTCAA GAATTCATGT CACACAAACC

7981	GATCTTCGCC TCAAGGAAAC CTAATTCTAC ATCCGAGAGA CTGCCGAGAT CTGTTCGGAA

8041	ATCAACGGAT GCTCAACCGA TTTCGACAGT AATAATTTGA ATCGAATCGG AGCCTAAAAT

8101	GAACCCGAGT ATATCTCATA AAATTCTCGG TGAGAGGTCT GTGACTGTCA GTACAAGGTG

8161	CCTTCATTAT GCCCTCAACC TTACCATACC TCACTGAATG TAGTGTACCT CTAAAAATGA

8221	AATACAGTGC CAAAAGCCAT GGCACTGAGC TCGTCTAACG GACTTGATAT ACAACCAATT

8281	AAAACAAATG AAAAGAAATA CAGTTCTTTG TATCATTTGT AACAATTACC CTGTACAAAC

8341	TAAGGTATTG AAATCCCACA ATATTCCCAA AGTCCACCCC TTTCCAAATT GTCATGCCTA

8401	CAACTCATAT ACCAAGCACT AACCTACCAA ACACCACTAA AACCCCACAA AATATATCTT

8461	ACCGAATATA CAGTAACAAG CTACCACCAC ACTCGTTGGG TGCAGTCGCC AGCTTAAAGA

8521	TATCTATCCA CATCAGCCAC AACTCCCTTC CTTTAATAAA CCGACTACAC CCTTGGCTAT

8581	TGAGGTTATG AGTGAATATA CTGTAGACAA GACACTTTCA AGAAGACTGT TTCCAAAACG

8641	TACCACTGTC CTCCACTACA AACACACCCA ATCTGCTTCT TCTAGTCAAG GTTGCTACAC

8701	CGGTAAATTA TAAATCATCA TTTCATTAGC AGGGCAGGGC CCTTTTTATA GAGTCTTATA

8761	CACTAGCGGA CCCTGCCGGT AGACCAACCC GCAGGCGCGT CAGTTTGCTC CTTCCATCAA

8821	TGCGTCGTAG AAACGACTTA CTCCTTCTTG AGCAGCTCCT TGACCTTGTT GGCAACAAGT

8881	CTCCGACCTC GGAGGTGGAG GAAGAGCCTC CGATATCGGC GGTAGTGATA CCAGCCTCGA

8941	CGGACTCCTT GACGGCAGCC TCAACAGCGT CACCGGCGGG CTTCATGTTA AGAGAGAACT

9001	TGAGCATCAT GGCGGCAGAC AGAATGGTGG CAATGGGGTT GACCTTCTGC TTGCCGAGAT

9061	CGGGGGCAGA TCCGTGACAG GGCTCGTACA GACCGAACGC CTCGTTGGTG TCGGGCAGAG

9121	AAGCCAGAGA GGCGGAGGGC AGCAGACCCA GAGAACCGGG GATGACGGAG GCCTCGTCGG

9181	AGATGATATC GCCAAACATG TTGGTGGTGA TGATGATACC ATTCATCTTG GAGGGCTGCT

9241	TGATGAGGAT CATGGCGGCC GAGTCGATCA GCTGGTGGTT GAGCTCGAGC TGGGGGAATT

9301	CGTCCTTGAG GACTCGAGTG ACAGTCTTTC GCCAAAGTCG AGAGGAGGCC AGCACGTTGG

9361	CCTTGTCAAG AGACCACACG GGAAGAGGGG GGTTGTGCTG AAGGGCCAGG AAGGCGGCCA

9421	TTCGGGCAAT TCGCTCAACC TCAGGAACGG AGTAGGTCTC GGTGTCGGAA GCGACGCCAG

9481	ATCCGTCATC CTCCTTTCGC TCTCCAAAGT AGATACCTCC GACGAGCTCT CGGACAATGA

9541	TGAAGTCGGT GCCCTCAACG TTTCGGATGG GGGAGAGATC GGCGAGCTTG GGCGACAGCA

9601	GCTGGCAGGG TCGCAGGTTG GCGTACAGGT TCAGGTCCTT TCGCAGCTTG AGGAGACCCT

9661	GCTCGGGTCG CACGTCGGTT CGTCCGTCGG GAGTGGTCCA TACGGTGTTG GCAGCGCCTC

9721	CGACAGCACC GAGCATAATA GAGTCAGCCT TTCGGCAGAT GTCGAGAGTA GCGTCGGTGA

9781	TGGGCTCGCC CTCCTTCTCA ATGGCAGCTC CTCCAATGAG TCGGTCCTCG AACACAAACT

9841	CGGTGCCGGA GGCCTCAGCA ACAGACTTGA GCACCTTGAC GGCCTCGGCA ATCACCTCGG

9901	GGCCACAGAA GTCGCCGCCG AGAAGAACAA TCTTCTTGGA GTCAGTCTTG GTCTTCTTAG

9961	TTTCGGGTTC CATTGTGGAT GTGTGTGGTT GTATGTGTGA TGTGGTGTGT GGAGTGAAAA

10021	TCTGTGGCTG GCAAACGCTC TTGTATATAT ACGCACTTTT GCCCGTGCTA TGTGGAAGAC

10081	TAAACCTCCG AAGATTGTGA CTCAGGTAGT GCGGTATCGG CTAGGGACCC AAACCTTGTC

10141	GATGCCGATA GCGCTATCGA ACGTACCCAG CCGGCCGGGA GTATGTCGGA GGGGACATAC

10201	GAGATCGTCA AGGGTTTGTG GCCAACTGGT AAATAAATGA TGACTCAGGC GACGACGGAA

10261	TTCTCATGTT TGACAGCTTA TCAT

Plasmid pUrLK sequences:

1	CGATGCTTTT CGTAGATAAT GGAATACAAA TGGATATCCA GAGTATACAC ATGGATAGTA

61	TACACTGACA CGACAATTCT GTATCTCTTT ATGTTAACTA CTGTGAGGCG TTAAATAGAG

121	CTTGATATAT AAAATGTTAC ATTTCACAGT CTGAACTTTT GCAGATTACC TAATTTGGTA

181	AGATATTAAT TATGAACTGA AAGTTGATGG CATCCCTAAA TTTGATGAAA GCCTAGGCGA

241	CTTGATGTTT AGAGTGTCCA GATCCGCAAG ATCGGCTCGC ACTTGTGTTG TGTTGTTTCA

301	AATCAGCCTG TCGTTTTGTG TCGTTTGAGA TCATTCTGTC TCACTCTTAG GCTCGCTTAG

361	AACCGACAAC GGAGAATCCG GGCTCGGTTT TTCGGTCGGC CTTGATCTGG GCCTTGGACT

421	TGTACTGGTC GGCCATCTCC ACGTTGACCA GCTCCTTGAC CTTGTAGAGC TGACCGGCGA

481	TACCAGGAGA CACCTTGTAG TACTTCTGGG AGCCGACCTT GCCCAGACCG AGGGTCTTGA

541	GCACGTCACG TGTTCTCCAC GGCATTCGCA GGATAGATCG GACCTGTGTG ACTTTGTAGA

601	ACATGGCGTT TCAGGTGGTT GCGTGAGTGT GTAAAATCGT GTCTTTCAGA AGTTACAAAT

661	TTCACCGCAT TTAGAGTTTA TGCAGATGGG CGGTGTGTGG TTGGGAGTTC GATTTCCGTG

721	CGTGCATTTG ATCTTGATGA ATTGGATTTG TACATGAGGA AGAGCACGTC AAGCACCGCC

781	TACTGCAAAC TCGTGAATAT TGAGATTATT GAGGAAATTC AAGGAAAATT CAGATCAGAT

841	TTGAGAGCAA AGTCCAACAA TACTACACAA TCCCTTTCCT GTATTCTTCC ACCATCGTCA

901	TCGTCGTCTG TCTTCTCTTC AGCTTTTTAA TTTCACTCCC CACAAACCCA AATTTAGCTG

961	CATCATTCAT CAACCTCCAA TTATAACTAT ACATCGCGAC ACGAACACGA AACACGAACC

1021	ACGAACCGCC GCTTTTTGAA AATAACTTCG TATAGCATAC ATTATACGAA GTTATGACGA

1081	CAGAGACCGG GTTGGCGGCG CATTTGTGTC CCAAAAAACA GCCCCAATTG CCCCAATTGA

1141	CCCCAAATTG ACCCAGTAGC GGGCCCAACC CCGGCGAGAG CCCCCTTCTC CCCACATATC

1201	AAACCTCCCC CGGTTCCCAC ACTTGCCGTT AAGGGCGTAG GGTACTGCAG TCTGGAATCT

1261	ACGCTTGTTC AGACTTTGTA CTTGTTTCTT TGTCTGGCCA TCCGGGTAAC CCATGCCGGA

1321	CGCAAAATAG ACTACTGAAA ATTTTTTTGC TTTGTGGTTG GGACTTTAGC CAAGGGTATA

1381	AAAGACCACC GTCCCCGAAT TACCTTTCCT CTTCTTTTCT CTCTCTCCTT GTCAACTCAC

1441	ACCCGAAATC GTTAAGCATT TCCTTCTGAG TATAAGAATC ATTCAAATCT AGAATGGTGA

1501	GTTTCAGAGG CAGCAGCAAT TGCCACGGGC TTTGAGCACA CGGCCGGGTG TGGTCCCATT

1561	CCCATCGACA CAAGACGCCA CGTCATCCGA CCAGCACTTT TTGCAGTACT AACCGCAGCC

1621	CTCCTACGAA GCTCGAGCTA ACGTCCACAA GTCCGCCTTT GCCGCTCGAG TGCTCAAGCT

1681	CGTGGCAGCC AAGAAAACCA ACCTGTGTGC TTCTCTGGAT GTTACCACCA CCAAGGAGCT

1741	CATTGAGCTT GCCGATAAGG TCGGACCTTA TGTGTGCATG ATCAAGACCC ATATCGACAT

1801	CATTGACGAC TTCACCTACG CCGGCACTGT GCTCCCCCTC AAGGAACTTG CTCTTAAGCA

1861	CGGTTTCTTC CTGTTCGAGG ACAGAAAGTT CGCAGATATT GGCAACACTG TCAAGCACCA

1921	GTACAAGAAC GGTGTCTACC GAATCGCCGA GTGGTCCGAT ATCACCAACG CCCACGGTGT

1981	ACCCGGAACC GGAATCATTG CTGGCCTGCG AGCTGGTGCC GAGGAAACTG TCTCTGAACA

2041	GAAGAAGGAG GACGTCTCTG ACTACGAGAA CTCCCAGTAC AAGGAGTTCC TGGTCCCCTC

2101	TCCCAACGAG AAGCTGGCCA GAGGTCTGCT CATGCTGGCC GAGCTGTCTT GCAAGGGCTC

2161	TCTGGCCACT GGCGAGTACT CCAAGCAGAC CATTGAGCTT GCCCGATCCG ACCCCGAGTT

2221	TGTGGTTGGC TTCATTGCCC AGAACCGACC TAAGGGCGAC TCTGAGGACT GGCTTATTCT

2281	GACCCCCGGG GTGGGTCTTG ACGACAAGGG AGACGCTCTC GGACAGCAGT ACCGAACTGT

2341	TGAGGATGTC ATGTCTACCG GAACGGATAT CATAATTGTC GGCCGAGGTC TGTACGGCCA

2401	GAACCGAGAT CCTATTGAGG AGGCCAAGCG ATACCAGAAG GCTGGCTGGG AGGCTTACCA

2461	GAAGATTAAC TGTTAGGGTA CCGACTAGTT CCATGGCCTG TCCCCACGTT GCCGGTCTTG

2521	CCTCCTACTA CCTGTCCATC AATGACGAGG TTCTCACCCC TGCCCAGGTC GAGGCTCTTA

2581	TTACTGAGTC CAACACCGGT GTTCTTCCCA CCACCAACCT CAAGGGCTCT CCCAACGCTG

2641	TTGCCTACAA CGGTGTTGGC ATTTAGGCAA TTAACAGATA GTTTGCCGGT GATAATTCTC

2701	TTAACCTCCC ACACTCCTTT GACATAACGA TTTATGTAAC GAAACTGAAA TTTGACCAGA

2761	TATTGTTGTA AATAGAAAAT CTGGCTTGTA GGTGGCAAAA TGCGGCGTCT TTGTTCATCA

2821	ATTCCCTCTG TGACTACTCG TCATCCCTTT ATGTTCGACT GTCGTATTTC TTATTTTCCA

2881	TACATATGCA AGTGAGATGC CCGTGTCCGT TATCAAATCT AGTTAATAAC TTCGTATAGC

2941	ATACATTATA CGAAGTTATG CTAGCGTCCG GAGCGGCCGC GCATGCAAGT CGACACTAGG

3001	GAGGCACATC TAAACGAATA ACGAATATTA ATGATACCAT CATATCTCAG AACATGTATG

3061	ACTGCTGCTT CCAAACGATA TGAGGATGAG TCCTCTTTCA GATTAAGATA GAGTACAAAT

3121	ATATTATCTA TATACTGGTG TCTGTGCGAT GTCGTATGAG CGGTGAATCA TGTGACTGTC

3181	ACGTGGTTTG GCCCAAGTTA CACCGTAGCT ACGCCTTTCT TGACCGTCTC CATGGTCTTC

3241	TGGGCGGGTT GACAGTTTCC ACTGGATGAG CGTCCGCCTC CTGTTCCTGT CGTTGTCCCT

3301	GCAGCTCAGC CTCAATCTTC TGACCGAGCT CGGAGTCCAG GGAAATGCCA ACAGGTTGTC

3361	CAAGCAACAT CATGGTTTGG TGGGCAGCCG TGATCTCATC GTCGTTGGAT ACCATTCGGT

3421	ACTTGGCCTC AATCTGCACA AAGTAGCGGT ACCACTGGTT TCGAGCAAAC CGCTCCAATT

3481	GAGCCTCTCC GTCGAGAGAG AGAGTAGGTG ATTGCTCCAA CTTGCGGCCA AAATGAAGTT

3541	CTCGACTCAC CTTTTTGAAG CGGTTCTTCT TGCCCATCTT GGTGGCGAAA GTAGTGGCTA

3601	GTGGTGGATG ACTTTGTATA ATGTACCGAT GAAGAGGGTT GTATTTGCTC AGTAAGAAGT

3661	AGCGAGTGAA ATCAGATGAC TTAACGAGAG CAAAGGGCAA TGGAATACCT GCTGCCTGAT

3721	TAACAACAGC TTCTGTGTCG TTTCTCTCTT GTGAATGAGT GTGTTGCTAG AGGTAGGTTG

3781	GCACTCCAAT GTTACGACAC ACAATAGTCT ATAGAGCACT ACAAAGGGCT ATATCGTCAA

3841	CTGCTCTATT GTAGCTACAG TACAGTACAT ACCATCAAGT GAACAATGGA CCACCAAACT

3901	CGGCACTAAG CCAATAGAAC CTTTGCGGCC TCCTTTATCA CGTTTCTATA TACCTTGTCC

3961	ATTTATGTGC CACCCTTTAG TCTTGGTCGT TCACTCCTAG GTATGTCTGA TAAAAGGATG

4021	TAACATAGGC AAGCTGCTCG TGAGTGTTGA GTACGAACCT TAGATCCAAA TCACCCGCAC

4081	CCACGGATAT ACTTGCTTGA ATATACAGTA GTATGCTCGA CCGATGCCCT TGAGAGCCTT

4141	CAACCCAGTC AGCTCCTTCC GGTGGGCGCG GGGCATGACT ATCGTCGCCG CACTTATGAC

4201	TGTCTTCTTT ATCATGCAAC TCGTAGGACA GGTGCCGGCA GCGCTCTGGG TCATTTTCGG

4261	CGAGGACCGC TTTCGCTGGA GCGCGACGAT GATCGGCCTG TCGCTTGCGG TATTCGGAAT

4321	CTTGCACGCC CTCGCTCAAG CCTTCGTCAC TGGTCCCGCC ACCAAACGTT TCGGCGAGAA

4381	GCAGGCCATT ATCGCCGGCA TGGCGGCCGA CGCGCTGGGC TACGTCTTGC TGGCGTTCGC

4441	GACGCGAGGC TGGATGGCCT TCCCCATTAT GATTCTTCTC GCTTCCGGCG GCATCGGGAT

4501	GCCCGCGTTG CAGGCCATGC TGTCCAGGCA GGTAGATGAC GACCATCAGG GACAGCTTCA

4561	AGGATCGCTC GCGGCTCTTA CCAGCCTAAC TTCGATCACT GGACCGCTGA TCGTCACGGC

4621	GATTTATGCC GCCTCGGCGA GCACATGGAA CGGGTTGGCA TGGATTGTAG GCGCCGCCCT

4681	ATACCTTGTC TGCCTCCCCG CGTTGCGTCG CGGTGCATGG AGCCGGGCCA CCTCGACCTG

4741	AATGGAAGCC GGCGGCACCT CGCTAACGGA TTCACCACTC CAAGAATTGG AGCCAATCAA

4801	TTCTTGCGGA GAACTGTGAA TGCGCAAACC AACCCTTGGC AGAACATATC CATCGCGTCC

4861	GCCATCTCCA GCAGCCGCAC GCGGCGCATC TCGGGCAGCG TTGGGTCCTG GCCACGGGTG

4921	CGCATGATCG TGCTCCTGTC GTTGAGGACC CGGCTAGGCT GGCGGGGTTG CCTTACTGGT

4981	TAGCAGAATG AATCACCGAT ACGCGAGCGA ACGTGAAGCG ACTGCTGCTG CAAAACGTCT

5041	GCGACCTGAG CAACAACATG AATGGTCTTC GGTTTCCGTG TTTCGTAAAG TCTGGAAACG

5101	CGGAAGTCAG CGCCCTGCAC CATTATGTTT CGGATCTGCA TCGCAGGATG CTGCTGGCTA

5161	CCCTGTGGAA CACCTACATC TGTATTAACG AAGCGCTGGC ATTGACCCTG AGTGATTTTT

5221	CTCTGGTCCC GCCGCATCCA TACCGCCAGT TGTTTACCCT CACAACGTTC CAGTAACCGG

5281	GCATGTTCAT CATCAGTAAC CCGTATCGTG AGCATCCTCT CTCGTTTCAT CGGTATCATT

5341	ACCCCCATGA ACAGAAATCC CCCTTACACG GAGGCATCAG TGACCAAACA GGAAAAAACC

5401	GCCCTTAACA TGGCCCGCTT TATCAGAAGC CAGACATTAA CGCTTCTGGA GAAACTCAAC

5461	GAGCTGGACG CGGATGAACA GGCAGACATC TGTGAATCGC TTCACGACCA CGCTGATGAG

5521	CTTTACCGCA GCAGATCCGC GGCCCCATAG GCCAATAGTG GATCTGCTGC CTCGCGCGTT

5581	TCGGTGATGA CGGTGAAAAC CTCTGACACA TGCAGCTCCC GGAGACGGTC ACAGCTTGTC

5641	TGTAAGCGGA TGCCGGGAGC AGACAAGCCC GTCAGGGCGC GTCAGCGGGT GTTGGCGGGT

5701	GTCGGGGCGC AGCCATGACC CAGTCACGTA GCGATAGCGG AGTGTATACT GGCTTAACTA

5761	TGCGGCATCA GAGCAGATTG TACTGAGAGT GCACCATATG CGGTGTGAAA TACCGCACAG

5821	ATGCGTAAGG AGAAAATACC GCATCAGGCG CTCTTCCGCT TCCTCGCTCA CTGACTCGCT

5881	GCGCTCGGTC GTTCGGCTGC GGCGAGCGGT ATCAGCTCAC TCAAAGGCGG TAATACGGTT

5941	ATCCACAGAA TCAGGGGATA ACGCAGGAAA GAACATGTGA GCAAAAGGCC AGCAAAAGGC

6001	CAGGAACCGT AAAAAGGCCG CGTTGCTGGC GTTTTTCCAT AGGCTCCGCC CCCCTGACGA

6061	GCATCACAAA AATCGACGCT CAAGTCAGAG GTGGCGAAAC CCGACAGGAC TATAAAGATA

6121	CCAGGCGTTT CCCCCTGGAA GCTCCCTCGT GCGCTCTCCT GTTCCGACCC TGCCGCTTAC

6181	CGGATACCTG TCCGCCTTTC TCCCTTCGGG AAGCGTGGCG CTTTCTCATA GCTCACGCTG

6241	TAGGTATCTC AGTTCGGTGT AGGTCGTTCG CTCCAAGCTG GGCTGTGTGC ACGAACCCCC

6301	CGTTCAGCCC GACCGCTGCG CCTTATCCGG TAACTATCGT CTTGAGTCCA ACCCGGTAAG

6361	ACACGACTTA TCGCCACTGG CAGCAGCCAC TGGTAACAGG ATTAGCAGAG CGAGGTATGT

6421	AGGCGGTGCT ACAGAGTTCT TGAAGTGGTG GCCTAACTAC GGCTACACTA GAAGGACAGT

6481	ATTTGGTATC TGCGCTCTGC TGAAGCCAGT TACCTTCGGA AAAAGAGTTG GTAGCTCTTG

6541	ATCCGGCAAA CAAACCACCG CTGGTAGCGG TGGTTTTTTT GTTTGCAAGC AGCAGATTAC

6601	GCGCAGAAAA AAAGGATCTC AAGAAGATCC TTTGATCTTT TCTACGGGGT CTGACGCTCA

6661	GTGGAACGAA AACTCACGTT AAGGGATTTT GGTCATGAGA TTATCAAAAA GGATCTTCAC

6721	CTAGATCCTT TTAAATTAAA AATGAAGTTT TAAATCAATC TAAAGTATAT ATGAGTAAAC

6781	TTGGTCTGAC AGTTACCAAT GCTTAATCAG TGAGGCACCT ATCTCAGCGA TCTGTCTATT

6841	TCGTTCATCC ATAGTTGCCT GACTCCCCGT CGTGTAGATA ACTACGATAC GGGAGGGCTT

6901	ACCATCTGGC CCCAGTGCTG CAATGATACC GCGAGACCCA CGCTCACCGG CTCCAGATTT

6961	ATCAGCAATA AACCAGCCAG CCGGAAGGGC CGAGCGCAGA AGTGGTCCTG CAACTTTATC

7021	CGCCTCCATC CAGTCTATTA ATTGTTGCCG GGAAGCTAGA GTAAGTAGTT CGCCAGTTAA

7081	TAGTTTGCGC AACGTTGTTG CCATTGCTGC AGGCATCGTG GTGTCACGCT CGTCGTTTGG

7141	TATGGCTTCA TTCAGCTCCG GTTCCCAACG ATCAAGGCGA GTTACATGAT CCCCCATGTT

7201	GTGCAAAAAA GCGGTTAGCT CCTTCGGTCC TCCGATCGTT GTCAGAAGTA AGTTGGCCGC

7261	AGTGTTATCA CTCATGGTTA TGGCAGCACT GCATAATTCT CTTACTGTCA TGCCATCCGT

7321	AAGATGCTTT TCTGTGACTG GTGAGTACTC AACCAAGTCA TTCTGAGAAT AGTGTATGCG

7381	GCGACCGAGT TGCTCTTGCC CGGCGTCAAC ACGGGATAAT ACCGCGCCAC ATAGCAGAAC

7441	TTTAAAAGTG CTCATCATTG GAAAACGTTC TTCGGGGCGA AAACTCTCAA GGATCTTACC

7501	GCTGTTGAGA TCCAGTTCGA TGTAACCCAC TCGTGCACCC AACTGATCTT CAGCATCTTT

7561	TACTTTCACC AGCGTTTCTG GGTGAGCAAA AACAGGAAGG CAAAATGCCG CAAAAAAGGG

7621	AATAAGGGCG ACACGGAAAT GTTGAATACT CATACTCTTC CTTTTTCAAT ATTATTGAAG

7681	CATTTATCAG GGTTATTGTC TCATGAGCGG ATACATATTT GAATGTATTT AGAAAAATAA

7741	ACAAATAGGG GTTCCGCGCA CATTTCCCCG AAAAGTGCCA CCTGACGTCT AAGAAACCAT

7801	TATTATCATG ACATTAACCT ATAAAAATAG GCGTATCACG AGGCCCTTTC GTCTTCAAGA

7861	ATTCATGTCA CACAAACCGA TCTTCGCCTC AAGGAAACCT AATTCTACAT CCGAGAGACT

7921	GCCGAGATCT GTTCGGAAAT CAACGGATGC TCAACCGATT TCGACAGTAA TAATTTGAAT

7981	CGAATCGGAG CCTAAAATGA ACCCGAGTAT ATCTCATAAA ATTCTCGGTG AGAGGTCTGT

8041	GACTGTCAGT ACAAGGTGCC TTCATTATGC CCTCAACCTT ACCATACCTC ACTGAATGTA

8101	GTGTACCTCT AAAAATGAAA TACAGTGCCA AAAGCCATGG CACTGAGCTC GTCTAACGGA

8161	CTTGATATAC AACCAATTAA AACAAATGAA AAGAAATACA GTTCTTTGTA TCATTTGTAA

8221	CAATTACCCT GTACAAACTA AGGTATTGAA ATCCCACAAT ATTCCCAAAG TCCACCCCTT

8281	TCCAAATTGT CATGCCTACA ACTCATATAC CAAGCACTAA CCTACCAAAC ACCACTAAAA

8341	CCCCACAAAA TATATCTTAC CGAATATACA GTAACAAGCT ACCACCACAC TCGTTGGGTG

8401	CAGTCGCCAG CTTAAAGATA TCTATCCACA TCAGCCACAA CTCCCTTCCT TTAATAAACC

8461	GACTACACCC TTGGCTATTG AGGTTATGAG TGAATATACT GTAGACAAGA CACTTTCAAG

8521	AAGACTGTTT CCAAAACGTA CCACTGTCCT CCACTACAAA CACACCCAAT CTGCTTCTTC

8581	TAGTCAAGGT TGCTACACCG GTAAATTATA AATCATCATT TCATTAGCAG GGCAGGGCCC

8641	TTTTTATAGA GTCTTATACA CTAGCGGACC CTGCCGGTAG ACCAACCCGC AGGCGCGTCA

8701	GTTTGCTCCT TCCATCAATG CGTCGTAGAA ACGACTTACT CCTTCTTGAG CAGCTCCTTG

8761	ACCTTGTTGG CAACAAGTCT CCGACCTCGG AGGTGGAGGA AGAGCCTCCG ATATCGGCGG

8821	TAGTGATACC AGCCTCGACG GACTCCTTGA CGGCAGCCTC AACAGCGTCA CCGGCGGGCT

8881	TCATGTTAAG AGAGAACTTG AGCATCATGG CGGCAGACAG AATGGTGGCA ATGGGGTTGA

8941	CCTTCTGCTT GCCGAGATCG GGGGCAGATC CGTGACAGGG CTCGTACAGA CCGAACGCCT

9001	CGTTGGTGTC GGGCAGAGAA GCCAGAGAGG CGGAGGGCAG CAGACCCAGA GAACCGGGGA

9061	TGACGGAGGC CTCGTCGGAG ATGATATCGC CAAACATGTT GGTGGTGATG ATGATACCAT

9121	TCATCTTGGA GGGCTGCTTG ATGAGGATCA TGGCGGCCGA GTCGATCAGC TGGTGGTTGA

9181	GCTCGAGCTG GGGGAATTCG TCCTTGAGGA CTCGAGTGAC AGTCTTTCGC CAAAGTCGAG

9241	AGGAGGCCAG CACGTTGGCC TTGTCAAGAG ACCACACGGG AAGAGGGGGG TTGTGCTGAA

9301	GGGCCAGGAA GGCGGCCATT CGGGCAATTC GCTCAACCTC AGGAACGGAG TAGGTCTCGG

9361	TGTCGGAAGC GACGCCAGAT CCGTCATCCT CCTTTCGCTC TCCAAAGTAG ATACCTCCGA

9421	CGAGCTCTCG GACAATGATG AAGTCGGTGC CCTCAACGTT TCGGATGGGG GAGAGATCGG

9481	CGAGCTTGGG CGACAGCAGC TGGCAGGGTC GCAGGTTGGC GTACAGGTTC AGGTCCTTTC

9541	GCAGCTTGAG GAGACCCTGC TCGGGTCGCA CGTCGGTTCG TCCGTCGGGA GTGGTCCATA

9601	CGGTGTTGGC AGCGCCTCCG ACAGCACCGA GCATAATAGA GTCAGCCTTT CGGCAGATGT

9661	CGAGAGTAGC GTCGGTGATG GGCTCGCCCT CCTTCTCAAT GGCAGCTCCT CCAATGAGTC

9721	GGTCCTCGAA CACAAACTCG GTGCCGGAGG CCTCAGCAAC AGACTTGAGC ACCTTGACGG

9781	CCTCGGCAAT CACCTCGGGG CCACAGAAGT CGCCGCCGAG AAGAACAATC TTCTTGGAGT

9841	CAGTCTTGGT CTTCTTAGTT TCGGGTTCCA TTGTGGATGT GTGTGGTTGT ATGTGTGATG

9901	TGGTGTGTGG AGTGAAAATC TGTGGCTGGC AAACGCTCTT GTATATATAC GCACTTTTGC

9961	CCGTGCTATG TGGAAGACTA AACCTCCGAA GATTGTGACT CAGGTAGTGC GGTATCGGCT

10021	AGGGACCCAA ACCTTGTCGA TGCCGATAGC GCTATCGAAC GTACCCAGCC GGCCGGGAGT

10081	ATGTCGGAGG GGACATACGA GATCGTCAAG GGTTTGTGGC CAACTGGTAA ATAAATGATG

10141	ACTCAGGCGA CGACGGAATT CTCATGTTTG ACAGCTTATC AT

Plasmid pUrLA sequences:

1	CGATGCTTTT CGTAGATAAT GGAATACAAA TGGATATCCA GAGTATACAC ATGGATAGTA

61	TACACTGACA CGACAATTCT GTATCTCTTT ATGTTAACTA CTGTGAGGCG TTAAATAGAG

121	CTTGATATAT AAAATGTTAC ATTTCACAGT CTGAACTTTT GCAGATTACC TAATTTGGTA

181	AGATATTAAT TATGAACTGA AAGTTGATGG CATCCCTAAA TTTGATGAAA GCCTAGGGCT

241	ATTCTTACGG TGTACAGTTA CGAGCACTTG TACTATAGTT TAGTTGTGGT TAAAACAACG

301	TTGTGGCACA ATTGTAGTCC TCGCAGATCC CTACAAAGGC CGGCCATTGC ACGCAAATTG

361	CACGAATTTT TCGCAGGCTT GCGCGACAAA TGCACCCAAA TACCGCACAA ATGACTCTGT

421	TTTTCACTTT CTCGTTTTCT CCATATTCTC CATATTCTCC ATTTTTTTGC CGTCCGCAAA

481	AACCCAATTC AACCCGGGGT ACAGGTTTGG TGCATGCTGT GCTAGATGAA AGTAGCCCTA

541	TTCCACTTCG GAAGGATGGG ATCCGAGATT CAAAGTTACC CTGTCTCATG ATTATCCGAA

601	CTTCCGGGTG TCCATGTGGT GCCAGGGACA GCGGCGCTAT GAGTAGAGAG ATGGAACGAG

661	AATTACAGGA AGGGGGGGAG ACACAGATCA TGGCTGATAC AGATGTACCT GCGTCTTTTA

721	CGTCTCCTGG GTCCTTCTCC ATGTGTATTT CGCACCCGCC AGGGCAATTA TATCCGCCTC

781	TCCATGTGTC CATTTTTGTG TATCAGCTGT TGGGCGCTTG TCAAGCACAC CAATGACAAA

841	CTTCATAATG ACTCTGGGGA AGGAAGCGCA GTTTGGTGGA AGTGTATGTA TGTAGTCTAA

901	GGGAGTCATG TTCCAGCATG AGCAGGAGAG GTGGGCGGAA GGGAGAAAAA ACAATCCGAT

961	ACCACTCCAA TGGTCATGTC ACCCCTCTCC TTCTATCTAC ACTCTGCCCT TCTCAACATA

1021	CGACCCTATG GACACGCCTC TACTAACCCC AACTCAGGGA CTGCGGCTCA ACATTCTGCA

1081	AGGCTTGGAA GAATTTGACT TTTGGCTTCC GTCAACTTTT TTATTTTGGA AGAAGTCAAA

1141	AAGCTAATAG CACACGCATC GCAAACAATA TCGAGAGACG CCTATATAAA GGCAGCGTGT

1201	CCTGATCCCT CCCTTCCGTA CTACCCCATC TCACACAATA ACTTCGTATA GCATACATTA

1261	TACGAAGTTA TGACGACAGA GACCGGGTTG GCGGCGCATT TGTGTCCCAA AAAACAGCCC

1321	CAATTGCCCC AATTGACCCC AAATTGACCC AGTAGCGGGC CCAACCCCGG CGAGAGCCCC

1381	CTTCTCCCCA CATATCAAAC CTCCCCCGGT TCCCACACTT GCCGTTAAGG GCGTAGGGTA

1441	CTGCAGTCTG GAATCTACGC TTGTTCAGAC TTTGTACTTG TTTCTTTGTC TGGCCATCCG

1501	GGTAACCCAT GCCGGACGCA AAATAGACTA CTGAAAATTT TTTTGCTTTG TGGTTGGGAC

1561	TTTAGCCAAG GGTATAAAAG ACCACCGTCC CCGAATTACC TTTCCTCTTC TTTTCTCTCT

1621	CTCCTTGTCA ACTCACACCC GAAATCGTTA AGCATTTCCT TCTGAGTATA AGAATCATTC

1681	AAATCTAGAA TGGTGAGTTT CAGAGGCAGC AGCAATTGCC ACGGGCTTTG AGCACACGGC

1741	CGGGTGTGGT CCCATTCCCA TCGACACAAG ACGCCACGTC ATCCGACCAG CACTTTTTGC

1801	AGTACTAACC GCAGCCCTCC TACGAAGCTC GAGCTAACGT CCACAAGTCC GCCTTTGCCG

1861	CTCGAGTGCT CAAGCTCGTG GCAGCCAAGA AAACCAACCT GTGTGCTTCT CTGGATGTTA

1921	CCACCACCAA GGAGCTCATT GAGCTTGCCG ATAAGGTCGG ACCTTATGTG TGCATGATCA

1981	AGACCCATAT CGACATCATT GACGACTTCA CCTACGCCGG CACTGTGCTC CCCCTCAAGG

2041	AACTTGCTCT TAAGCACGGT TTCTTCCTGT TCGAGGACAG AAAGTTCGCA GATATTGGCA

2101	ACACTGTCAA GCACCAGTAC AAGAACGGTG TCTACCGAAT CGCCGAGTGG TCCGATATCA

2161	CCAACGCCCA CGGTGTACCC GGAACCGGAA TCATTGCTGG CCTGCGAGCT GGTGCCGAGG

2221	AAACTGTCTC TGAACAGAAG AAGGAGGACG TCTCTGACTA CGAGAACTCC CAGTACAAGG

2281	AGTTCCTGGT CCCCTCTCCC AACGAGAAGC TGGCCAGAGG TCTGCTCATG CTGGCCGAGC

2341	TGTCTTGCAA GGGCTCTCTG GCCACTGGCG AGTACTCCAA GCAGACCATT GAGCTTGCCC

2401	GATCCGACCC CGAGTTTGTG GTTGGCTTCA TTGCCCAGAA CCGACCTAAG GGCGACTCTG

2461	AGGACTGGCT TATTCTGACC CCCGGGGTGG GTCTTGACGA CAAGGGAGAC GCTCTCGGAC

2521	AGCAGTACCG AACTGTTGAG GATGTCATGT CTACCGGAAC GGATATCATA ATTGTCGGCC

2581	GAGGTCTGTA CGGCCAGAAC CGAGATCCTA TTGAGGAGGC CAAGCGATAC CAGAAGGCTG

2641	GCTGGGAGGC TTACCAGAAG ATTAACTGTT AGGGTACCGA CTAGTTCCAT GGCCTGTCCC

2701	CACGTTGCCG GTCTTGCCTC CTACTACCTG TCCATCAATG ACGAGGTTCT CACCCCTGCC

2761	CAGGTCGAGG CTCTTATTAC TGAGTCCAAC ACCGGTGTTC TTCCCACCAC CAACCTCAAG

2821	GGCTCTCCCA ACGCTGTTGC CTACAACGGT GTTGGCATTT AGGCAATTAA CAGATAGTTT

2881	GCCGGTGATA ATTCTCTTAA CCTCCCACAC TCCTTTGACA TAACGATTTA TGTAACGAAA

2941	CTGAAATTTG ACCAGATATT GTTGTAAATA GAAAATCTGG CTTGTAGGTG GCAAAATGCG

3001	GCGTCTTTGT TCATCAATTC CCTCTGTGAC TACTCGTCAT CCCTTTATGT TCGACTGTCG

3061	TATTTCTTAT TTTCCATACA TATGCAAGTG AGATGCCCGT GTCCGTTATC AAATCTAGTT

3121	AATAACTTCG TATAGCATAC ATTATACGAA GTTATGCTAG CGTCCGGAGC GGCCGCGCAT

3181	GCAAGTCGAC AACACTATAT AAGAATGTAT TTATTTTCTC TCTTAACCTA TTGATGTTCC

3241	TGATCTTGAG CTACTTGGAC TGGTACAGTA GCTATAGATG AGCCTTATGC ATGTCTCACA

3301	CTAACTACAT AATATCTAAA AAAGCCGCTA ACGGCGTCTA CCCTTGTATA GTGCGGCTAC

3361	AACCAAAGAT ACCACTTTGT GTCCAATCTC CATTGTGGGG TGTCAATGGC GTGACGTGAA

3421	GACGCGTGCG GACAGGCTTT GTAGATAAGG GTTGTAGATA AGACGTTATT GGAATGACAT

3481	ATGTATTGTA CAGTGTAATT GTGCTGTAAG TACTCGTAAT TGCATCTGAC TGAGCTGTCT

3541	TAATGATGTA TCGGGTTGAC ATATCTGAAC GTCGAAATTT GACACCTAGT TGCTTCTGAA

3601	CACTCATCCT AATCACTGTT GATAATAGCT CAACACTCAT TGGAGCGCCG GAAATTAGTC

3661	TTAGTCAGTT GTATTGTTCG TGCGATAGCT ACTGTATGTA CTGTAATACG GCTAATAGTG

3721	TAATTTCCCC GCCAGGGGTT TTGGTTGAGT CTAGTTCCAT CATGTGTTGG GAGAGCCAGA

3781	ATACGATATT GTACAGGCAA ATAATACTCG TCTTATTTAG TCCTCGTACA GCACAAAGAA

3841	AGTTGTTATC AGTGCCGATG TGCAGTATGT CGGCCGAGTC GGGACACAAG CAGAGAGTAC

3901	GGGAGATGCA GTGGTCCGAC TCGGTTGTTC TCTGCTGTCG TACTTGTACC AGTGCAGTAT

3961	ACACCAACAA GGATTTCTCG TCATACGGTC ATTGCAGCTG TGCTTCATGC GCTGTCAGGT

4021	AATATCATAT CATGTCACCC CCATTTTGTG AGATCTCACT ATAAAGCATG TTCGTTATAC

4081	TGTTATTCAA TGTCAACCAA GCTCTGAGCA GCTACTGCTG TAGTTATTAG TTGTAGTGTA

4141	CAACAGCTAA GTATGAACCG GAAAGCATGT ATGAGTCAAC CTACTCGCGT CCCTAGGTAT

4201	GTCTGATAAA AGGATGTAAC ATAGGCAAGC TGCTCGTGAG TGTTGAGTAC GAACCTTAGA

4261	TCCAAATCAC CCGCACCCAC GGATATACTT GCTTGAATAT ACAGTAGTAT GCTCGACCGA

4321	TGCCCTTGAG AGCCTTCAAC CCAGTCAGCT CCTTCCGGTG GGCGCGGGGC ATGACTATCG

4381	TCGCCGCACT TATGACTGTC TTCTTTATCA TGCAACTCGT AGGACAGGTG CCGGCAGCGC

4441	TCTGGGTCAT TTTCGGCGAG GACCGCTTTC GCTGGAGCGC GACGATGATC GGCCTGTCGC

4501	TTGCGGTATT CGGAATCTTG CACGCCCTCG CTCAAGCCTT CGTCACTGGT CCCGCCACCA

4561	AACGTTTCGG CGAGAAGCAG GCCATTATCG CCGGCATGGC GGCCGACGCG CTGGGCTACG

4621	TCTTGCTGGC GTTCGCGACG CGAGGCTGGA TGGCCTTCCC CATTATGATT CTTCTCGCTT

4681	CCGGCGGCAT CGGGATGCCC GCGTTGCAGG CCATGCTGTC CAGGCAGGTA GATGACGACC

4741	ATCAGGGACA GCTTCAAGGA TCGCTCGCGG CTCTTACCAG CCTAACTTCG ATCACTGGAC

4801	CGCTGATCGT CACGGCGATT TATGCCGCCT CGGCGAGCAC ATGGAACGGG TTGGCATGGA

4861	TTGTAGGCGC CGCCCTATAC CTTGTCTGCC TCCCCGCGTT GCGTCGCGGT GCATGGAGCC

4921	GGGCCACCTC GACCTGAATG GAAGCCGGCG GCACCTCGCT AACGGATTCA CCACTCCAAG

4981	AATTGGAGCC AATCAATTCT TGCGGAGAAC TGTGAATGCG CAAACCAACC CTTGGCAGAA

5041	CATATCCATC GCGTCCGCCA TCTCCAGCAG CCGCACGCGG CGCATCTCGG GCAGCGTTGG

5101	GTCCTGGCCA CGGGTGCGCA TGATCGTGCT CCTGTCGTTG AGGACCCGGC TAGGCTGGCG

5161	GGGTTGCCTT ACTGGTTAGC AGAATGAATC ACCGATACGC GAGCGAACGT GAAGCGACTG

5221	CTGCTGCAAA ACGTCTGCGA CCTGAGCAAC AACATGAATG GTCTTCGGTT TCCGTGTTTC

5281	GTAAAGTCTG GAAACGCGGA AGTCAGCGCC CTGCACCATT ATGTTTCGGA TCTGCATCGC

5341	AGGATGCTGC TGGCTACCCT GTGGAACACC TACATCTGTA TTAACGAAGC GCTGGCATTG

5401	ACCCTGAGTG ATTTTTCTCT GGTCCCGCCG CATCCATACC GCCAGTTGTT TACCCTCACA

5461	ACGTTCCAGT AACCGGGCAT GTTCATCATC AGTAACCCGT ATCGTGAGCA TCCTCTCTCG

5521	TTTCATCGGT ATCATTACCC CCATGAACAG AAATCCCCCT TACACGGAGG CATCAGTGAC

5581	CAAACAGGAA AAAACCGCCC TTAACATGGC CCGCTTTATC AGAAGCCAGA CATTAACGCT

5641	TCTGGAGAAA CTCAACGAGC TGGACGCGGA TGAACAGGCA GACATCTGTG AATCGCTTCA

5701	CGACCACGCT GATGAGCTTT ACCGCAGCAG ATCCGCGGCC CCATAGGCCA ATAGTGGATC

5761	TGCTGCCTCG CGCGTTTCGG TGATGACGGT GAAAACCTCT GACACATGCA GCTCCCGGAG

5821	ACGGTCACAG CTTGTCTGTA AGCGGATGCC GGGAGCAGAC AAGCCCGTCA GGGCGCGTCA

5881	GCGGGTGTTG GCGGGTGTCG GGGCGCAGCC ATGACCCAGT CACGTAGCGA TAGCGGAGTG

5941	TATACTGGCT TAACTATGCG GCATCAGAGC AGATTGTACT GAGAGTGCAC CATATGCGGT

6001	GTGAAATACC GCACAGATGC GTAAGGAGAA AATACCGCAT CAGGCGCTCT TCCGCTTCCT

6061	CGCTCACTGA CTCGCTGCGC TCGGTCGTTC GGCTGCGGCG AGCGGTATCA GCTCACTCAA

6121	AGGCGGTAAT ACGGTTATCC ACAGAATCAG GGGATAACGC AGGAAAGAAC ATGTGAGCAA

6181	AAGGCCAGCA AAAGGCCAGG AACCGTAAAA AGGCCGCGTT GCTGGCGTTT TTCCATAGGC

6241	TCCGCCCCCC TGACGAGCAT CACAAAAATC GACGCTCAAG TCAGAGGTGG CGAAACCCGA

6301	CAGGACTATA AAGATACCAG GCGTTTCCCC CTGGAAGCTC CCTCGTGCGC TCTCCTGTTC

6361	CGACCCTGCC GCTTACCGGA TACCTGTCCG CCTTTCTCCC TTCGGGAAGC GTGGCGCTTT

6421	CTCATAGCTC ACGCTGTAGG TATCTCAGTT CGGTGTAGGT CGTTCGCTCC AAGCTGGGCT

6481	GTGTGCACGA ACCCCCCGTT CAGCCCGACC GCTGCGCCTT ATCCGGTAAC TATCGTCTTG

6541	AGTCCAACCC GGTAAGACAC GACTTATCGC CACTGGCAGC AGCCACTGGT AACAGGATTA

6601	GCAGAGCGAG GTATGTAGGC GGTGCTACAG AGTTCTTGAA GTGGTGGCCT AACTACGGCT

6661	ACACTAGAAG GACAGTATTT GGTATCTGCG CTCTGCTGAA GCCAGTTACC TTCGGAAAAA

6721	GAGTTGGTAG CTCTTGATCC GGCAAACAAA CCACCGCTGG TAGCGGTGGT TTTTTTGTTT

6781	GCAAGCAGCA GATTACGCGC AGAAAAAAAG GATCTCAAGA AGATCCTTTG ATCTTTTCTA

6841	CGGGGTCTGA CGCTCAGTGG AACGAAAACT CACGTTAAGG GATTTTGGTC ATGAGATTAT

6901	CAAAAAGGAT CTTCACCTAG ATCCTTTTAA ATTAAAAATG AAGTTTTAAA TCAATCTAAA

6961	GTATATATGA GTAAACTTGG TCTGACAGTT ACCAATGCTT AATCAGTGAG GCACCTATCT

7021	CAGCGATCTG TCTATTTCGT TCATCCATAG TTGCCTGACT CCCCGTCGTG TAGATAACTA

7081	CGATACGGGA GGGCTTACCA TCTGGCCCCA GTGCTGCAAT GATACCGCGA GACCCACGCT

7141	CACCGGCTCC AGATTTATCA GCAATAAACC AGCCAGCCGG AAGGGCCGAG CGCAGAAGTG

7201	GTCCTGCAAC TTTATCCGCC TCCATCCAGT CTATTAATTG TTGCCGGGAA GCTAGAGTAA

7261	GTAGTTCGCC AGTTAATAGT TTGCGCAACG TTGTTGCCAT TGCTGCAGGC ATCGTGGTGT

7321	CACGCTCGTC GTTTGGTATG GCTTCATTCA GCTCCGGTTC CCAACGATCA AGGCGAGTTA

7381	CATGATCCCC CATGTTGTGC AAAAAAGCGG TTAGCTCCTT CGGTCCTCCG ATCGTTGTCA

7441	GAAGTAAGTT GGCCGCAGTG TTATCACTCA TGGTTATGGC AGCACTGCAT AATTCTCTTA

7501	CTGTCATGCC ATCCGTAAGA TGCTTTTCTG TGACTGGTGA GTACTCAACC AAGTCATTCT

7561	GAGAATAGTG TATGCGGCGA CCGAGTTGCT CTTGCCCGGC GTCAACACGG GATAATACCG

7621	CGCCACATAG CAGAACTTTA AAAGTGCTCA TCATTGGAAA ACGTTCTTCG GGGCGAAAAC

7681	TCTCAAGGAT CTTACCGCTG TTGAGATCCA GTTCGATGTA ACCCACTCGT GCACCCAACT

7741	GATCTTCAGC ATCTTTTACT TTCACCAGCG TTTCTGGGTG AGCAAAAACA GGAAGGCAAA

7801	ATGCCGCAAA AAAGGGAATA AGGGCGACAC GGAAATGTTG AATACTCATA CTCTTCCTTT

7861	TTCAATATTA TTGAAGCATT TATCAGGGTT ATTGTCTCAT GAGCGGATAC ATATTTGAAT

7921	GTATTTAGAA AAATAAACAA ATAGGGGTTC CGCGCACATT TCCCCGAAAA GTGCCACCTG

7981	ACGTCTAAGA AACCATTATT ATCATGACAT TAACCTATAA AAATAGGCGT ATCACGAGGC

8041	CCTTTCGTCT TCAAGAATTC ATGTCACACA AACCGATCTT CGCCTCAAGG AAACCTAATT

8101	CTACATCCGA GAGACTGCCG AGATCTGTTC GGAAATCAAC GGATGCTCAA CCGATTTCGA

8161	CAGTAATAAT TTGAATCGAA TCGGAGCCTA AAATGAACCC GAGTATATCT CATAAAATTC

8221	TCGGTGAGAG GTCTGTGACT GTCAGTACAA GGTGCCTTCA TTATGCCCTC AACCTTACCA

8281	TACCTCACTG AATGTAGTGT ACCTCTAAAA ATGAAATACA GTGCCAAAAG CCATGGCACT

8341	GAGCTCGTCT AACGGACTTG ATATACAACC AATTAAAACA AATGAAAAGA AATACAGTTC

8401	TTTGTATCAT TTGTAACAAT TACCCTGTAC AAACTAAGGT ATTGAAATCC CACAATATTC

8461	CCAAAGTCCA CCCCTTTCCA AATTGTCATG CCTACAACTC ATATACCAAG CACTAACCTA

8521	CCAAACACCA CTAAAACCCC ACAAAATATA TCTTACCGAA TATACAGTAA CAAGCTACCA

8581	CCACACTCGT TGGGTGCAGT CGCCAGCTTA AAGATATCTA TCCACATCAG CCACAACTCC

8641	CTTCCTTTAA TAAACCGACT ACACCCTTGG CTATTGAGGT TATGAGTGAA TATACTGTAG

8701	ACAAGACACT TTCAAGAAGA CTGTTTCCAA AACGTACCAC TGTCCTCCAC TACAAACACA

8761	CCCAATCTGC TTCTTCTAGT CAAGGTTGCT ACACCGGTAA ATTATAAATC ATCATTTCAT

8821	TAGCAGGGCA GGGCCCTTTT TATAGAGTCT TATACACTAG CGGACCCTGC CGGTAGACCA

8881	ACCCGCAGGC GCGTCAGTTT GCTCCTTCCA TCAATGCGTC GTAGAAACGA CTTACTCCTT

8941	CTTGAGCAGC TCCTTGACCT TGTTGGCAAC AAGTCTCCGA CCTCGGAGGT GGAGGAAGAG

9001	CCTCCGATAT CGGCGGTAGT GATACCAGCC TCGACGGACT CCTTGACGGC AGCCTCAACA

9061	GCGTCACCGG CGGGCTTCAT GTTAAGAGAG AACTTGAGCA TCATGGCGGC AGACAGAATG

9121	GTGGCAATGG GGTTGACCTT CTGCTTGCCG AGATCGGGGG CAGATCCGTG ACAGGGCTCG

9181	TACAGACCGA ACGCCTCGTT GGTGTCGGGC AGAGAAGCCA GAGAGGCGGA GGGCAGCAGA

9241	CCCAGAGAAC CGGGGATGAC GGAGGCCTCG TCGGAGATGA TATCGCCAAA CATGTTGGTG

9301	GTGATGATGA TACCATTCAT CTTGGAGGGC TGCTTGATGA GGATCATGGC GGCCGAGTCG

9361	ATCAGCTGGT GGTTGAGCTC GAGCTGGGGG AATTCGTCCT TGAGGACTCG AGTGACAGTC

9421	TTTCGCCAAA GTCGAGAGGA GGCCAGCACG TTGGCCTTGT CAAGAGACCA CACGGGAAGA

9481	GGGGGGTTGT GCTGAAGGGC CAGGAAGGCG GCCATTCGGG CAATTCGCTC AACCTCAGGA

9541	ACGGAGTAGG TCTCGGTGTC GGAAGCGACG CCAGATCCGT CATCCTCCTT TCGCTCTCCA

9601	AAGTAGATAC CTCCGACGAG CTCTCGGACA ATGATGAAGT CGGTGCCCTC AACGTTTCGG

9661	ATGGGGGAGA GATCGGCGAG CTTGGGCGAC AGCAGCTGGC AGGGTCGCAG GTTGGCGTAC

9721	AGGTTCAGGT CCTTTCGCAG CTTGAGGAGA CCCTGCTCGG GTCGCACGTC GGTTCGTCCG

9781	TCGGGAGTGG TCCATACGGT GTTGGCAGCG CCTCCGACAG CACCGAGCAT AATAGAGTCA

9841	GCCTTTCGGC AGATGTCGAG AGTAGCGTCG GTGATGGGCT CGCCCTCCTT CTCAATGGCA

9901	GCTCCTCCAA TGAGTCGGTC CTCGAACACA AACTCGGTGC CGGAGGCCTC AGCAACAGAC

9961	TTGAGCACCT TGACGGCCTC GGCAATCACC TCGGGGCCAC AGAAGTCGCC GCCGAGAAGA

10021	ACAATCTTCT TGGAGTCAGT CTTGGTCTTC TTAGTTTCGG GTTCCATTGT GGATGTGTGT

10081	GGTTGTATGT GTGATGTGGT GTGTGGAGTG AAAATCTGTG GCTGGCAAAC GCTCTTGTAT

10141	ATATACGCAC TTTTGCCCGT GCTATGTGGA AGACTAAACC TCCGAAGATT GTGACTCAGG

10201	TAGTGCGGTA TCGGCTAGGG ACCCAAACCT TGTCGATGCC GATAGCGCTA TCGAACGTAC

10261	CCAGCCGGCC GGGAGTATGT CGGAGGGGAC ATACGAGATC GTCAAGGGTT TGTGGCCAAC

10321	TGGTAAATAA ATGATGACTC AGGCGACGAC GGAATTCTCA TGTTTGACAG CTTATCAT

rgTAL sequences:

1	ATGGCCCCTT CGCTGGACAG CATCTCCCAT TCGTTCGCAA ACGGCGTTGC ATCTGCTAAG

61	CAGGCAGTGA ACGGCGCATC GACAAACCTG GCTGTGGCTG GCAGCCACCT GCCCACCACA

121	CAAGTCACTC AGGTTGACAT TGTTGAAAAA ATGCTTGCCG CACCCACGGA TAGCACTCTG

181	GAGCTGGACG GATACTCCCT TAATCTGGGC GATGTGGTCA GCGCTGCACG GAAGGGCCGG

241	CCCGTTCGTG TGAAGGATTC TGACGAAATT AGATCTAAGA TCGACAAATC CGTGGAATTT

301	CTTAGATCTC AACTTTCGAT GTCTGTGTAC GGCGTCACTA CAGGCTTCGG CGGTTCGGCT

361	GATACTCGGA CTGAAGACGC TATCTCGCTT CAAAAGGCTC TCCTTGAACA TCAGCTCTGT

421	GGCGTCCTTC CTAGCTCCTT TGACAGCTTT AGACTCGGAA GAGGCCTCGA AAACTCGCTC

481	CCCCTCGAAG TTGTTCGGGG AGCAATGACA ATTCGTGTCA ACTCGCTTAC ACGAGGTCAT

541	TCTGCAGTCC GGCTGGTTGT GCTGGAGGCT CTTACTAATT TTCTGAATCA CGGAATTACC

601	CCCATCGTTC CCCTTCGGGG TACAATTTCT GCAAGCGGTG ACCTGTCTCC CCTGTCGTAT

661	ATCGCAGCTG CCATTTCGGG CCACCCTGAT TCTAAAGTGC ACGTCGTGCA CGAAGGAAAA

721	GAAAAAATCC TGTATGCACG TGAAGCAATG GCTCTCTTCA ATCTTGAACC TGTTGTGCTC

781	GGTCCCAAAG AAGGCCTGGG TCTCGTTAAT GGCACCGCTG TCTCTGCCTC TATGGCTACG

841	CTTGCTCTGC ATGACGCACA CATGCTGTCT CTTCTGTCTC AGTCCCTGAC CGCTATGACT

901	GTGGAGGCTA TGGTTGGTCA CGCCGGTTCG TTCCACCCCT TTCTTCATGA CGTCACTCGA

961	CCCCATCCCA CTCAAATTGA AGTCGCAGGC AACATTAGAA AGCTCCTGGA GGGATCCCGA

1021	TTTGCCGTCC ATCACGAAGA AGAAGTGAAG GTCAAGGACG ATGAGGGAAT CCTGCGGCAA

1081	GATCGTTACC CCCTCAGAAC GTCTCCCCAG TGGCTGGGAC CCCTGGTTTC TGACCTGATC

1141	CATGCCCACG CTGTTCTGAC GATCGAAGCA GGACAATCTA CTACTGATAA TCCCCTGATT

1201	GATGTGGAAA ACAAAACCTC CCATCATGGT GGCAACTTTC AAGCAGCTGC TGTGGCTAAC

1261	ACAATGGAAA AGACACGTCT TGGTCTTGCC CAGATCGGCA AGCTCAACTT CACTCAACTT

1321	ACCGAGATGC TCAACGCAGG AATGAACCGT GGTCTGCCTA GCTGCCTGGC AGCCGAGGAT

1381	CCCTCTCTTT CTTACCATTG TAAAGGCCTC GACATCGCAG CTGCCGCTTA CACTTCCGAG

1441	CTGGGCCACC TTGCAAATCC TGTGACCACT CATGTGCAGC CCGCTGAAAT GGCCAACCAA

1501	GCAGTCAACT CCCTTGCTCT CATCTCGGCA CGACGGACCA CTGAATCCAA CGATGTGCTC

1561	TCCCTCCTCC TTGCAACACA TCTCTACTGT GTCCTTCAGG CTATCGATCT CCGAGCAATC

1621	GAGTTTGAAT TCAAGAAACA GTTTGGTCCC GCCATTGTGT CGCTGATCGA TCAACACTTC

1681	GGTTCCGCCA TGACAGGCTC TAATCTGCGT GACGAGCTTG TCGAGAAAGT TAACAAGACG

1741	CTCGCCAAGC GGCTGGAACA GACAAACAGC TACGATCTTG TTCCTCGATG GCATGATGCC

1801	TTTTCGTTTG CTGCAGGCAC AGTGGTGGAA GTGCTGTCCT CTACGTCTCT TTCTCTTGCC

1861	GCCGTTAACG CCTGGAAAGT GGCCGCTGCC GAGAGCGCAA TCTCTCTTAC CCGGCAAGTC

1921	CGTGAAACCT TTTGGAGCGC AGCCAGCACT TCCAGCCCTG CTCTGTCGTA CCTTTCGCCC

1981	CGGACTCAGA TCCTGTATGC TTTTGTCCGT GAGGAGCTTG GTGTCAAGGC TCGTCGTGGA

2041	GATGTCTTCC TTGGAAAGCA AGAGGTCACA ATTGGTAGCA ACGTGTCGAA AATTTACGAA

2101	GCTATTAAGT CCGGCCGTAT CAACAACGTT CTGCTGAAAA TGCTTGCTTA A

VvSTS1 sequences:

1	ATGGCTTCCG TGGAAGAATT CCGTAATGCT CAGCGAGCCA AAGGCCCCGC CACCATCCTC

61	GCAATTGGCA CCGCCACTCC TGACCATTGC GTCTATCAAT CCGACTACGC CGACTACTAT

121	TTTCGGGTTA CGAAGTCCGA GCACATGACA GAACTGAAAA AAAAATTTAA CCGAATCTGC

181	GACAAGAGCA TGATCAAGAA GCGATACATT CATCTTACGG AGGAAATGCT TGAGGAACAT

241	CCCAACATCG GTGCATATAT GGCCCCTTCG CTTAACATTC GTCAAGAGAT TATTACTGCA

301	GAAGTTCCCA GACTTGGACG AGACGCAGCC CTTAAGGCCC TCAAAGAATG GGGTCAACCC

361	AAGTCCAAGA TTACGCATCT TGTTTTTTGC ACAACCAGCG GCGTCGAGAT GCCTGGTGCC

421	GATTATAAGC TCGCAAATCT TCTCGGCCTG GAGACTAGCG TCCGGCGTGT GATGCTCTAC

481	CATCAGGGTT GCTATGCAGG CGGAACTGTG CTCCGAACGG CAAAAGACCT CGCAGAAAAC

541	AATGCCGGAG CTCGAGTCCT CGTTGTTTGC AGCGAGATCA CAGTCGTGAC CTTCCGTGGC

601	CCCAGCGAGG ACGCTCTCGA CTCCCTCGTC GGCCAAGCTC TCTTCGGTGA TGGCTCTTCT

661	GCCGTGATCG TCGGATCTGA TCCCGATGTC TCCATTGAGC GTCCTCTTTT CCAGCTTGTG

721	AGCGCTGCCC AGACGTTCAT TCCTAACTCT GCAGGCGCTA TTGCAGGAAA TCTTCGGGAG

781	GTTGGCCTCA CCTTCCATCT CTGGCCTAAT GTGCCCACCC TTATCAGCGA GAATATCGAA

841	AAATGTCTGA CCCAGGCATT TGACCCTCTC GGCATCTCGG ATTGGAATAG CCTGTTCTGG

901	ATTGCACATC CTGGCGGTCC TGCAATCCTT GATGCAGTGG AGGCTAAGCT GAACCTTGAG

961	AAAAAGAAAC TGGAAGCCAC AAGACATGTT CTTTCCGAGT ATGGAAACAT GTCTTCCGCA

1021	TGTGTCCTGT TCATTCTGGA TGAGATGCGT AAGAAATCCC TGAAAGGTGA GAAAGCCACC

1081	ACCGGTGAGG GTCTGGACTG GGGAGTGCTC TTCGGATTCG GCCCCGGTCT TACAATCGAG

1141	ACCGTCGTCC TGCACTCCGT TCCCACCGTG ACGAACTAA

Pc4CL2 sequences:

1	ATGGGCGATT GTGTTGCACC CAAGGAAGAT CTTATCTTTC GGAGCAAGCT CCCTGATATC

61	TACATCCCCA AACACCTGCC TCTTCACACA TATTGCTTCG AGAACATCTC TAAAGTCGGT

121	GACAAGTCCT GTCTCATTAA CGGTGCTACG GGTGAGACCT TCACTTATTC GCAAGTGGAA

181	CTTCTCTCCC GTAAGGTGGC ATCGGGTCTC AACAAGCTTG GCATTCAGCA GGGAGACACT

241	ATTATGCTGC TCCTGCCCAA CTCCCCTGAG TATTTTTTCG CCTTTCTTGG CGCCTCCTAC

301	CGTGGTGCCA TTTCGACGAT GGCAAACCCC TTCTTTACAT CCGCAGAAGT CATTAAACAG

361	CTCAAAGCAT CCCTTGCTAA GCTCATTATT ACGCAGGCTT GCTACGTGGA CAAGGTTAAG

421	GATTACGCTG CTGAAAAAAA TATCCAGATT ATTTGTATTG ACGACGCCCC TCAAGACTGC

481	CTTCACTTCT CGAAGCTGAT GGAGGCAGAT GAGAGCGAAA TGCCCGAGGT CGTGATCGAC

541	TCGGATGACG TTGTTGCCCT CCCCTATTCC AGCGGCACCA CGGGTCTTCC TAAAGGCGTG

601	ATGCTCACAC ATAAAGGACT TGTGACATCC GTCGCCCAAC AAGTGGATGG AGATAATCCT

661	AATCTTTATA TGCATTCCGA AGACGTTATG ATCTGCATCC TCCCTCTGTT CCACATCTAC

721	TCTCTTAATG CTGTCCTTTG CTGCGGTCTC CGTGCTGGCG TGACAATTCT TATCATGCAG

781	AAGTTTGATA TTGTTCCTTT CCTGGAGCTG ATTCAGAAGT ATAAAGTTAC GATCGGTCCC

841	TTTGTGCCTC CCATTGTTCT GGCAATTGCA AAGTCCCCTG TTGTTGATAA GTACGATCTG

901	TCGTCCGTTC GAACGGTGAT GTCTGGAGCA GCACCTCTTG GAAAAGAACT GGAAGATGCA

961	GTTAGAGCTA AGTTTCCCAA TGCTAAACTG GGACAAGGTT ACGGTATGAC AGAGGCCGGT

1021	CCTGTTCTTG CAATGTGTCT CGCTTTCGCA AAGGAGCCCT ATGAAATCAA ATCTGGCGCT

1081	TGTGGTACTG TTGTCCGAAA CGCTGAGATG AAGATTGTGG ACCCTGAAAC TAACGCCTCT

1141	CTTCCCAGAA ACCAAAGAGG TGAAATTTGC ATTCGAGGCG ATCAAATTAT GAAGGGATAC

1201	CTGAACGACC CTGAATCTAC GCGAACGACA ATTGACGAGG AAGGTTGGCT TCACACCGGA

1261	GATATTGGAT TCATTGATGA TGATGACGAG CTCTTCATTG TCGACCGTCT TAAAGAGATC

1321	ATCAAGTATA AGGGTTTCCA GGTCGCCCCT GCTGAGCTGG AGGCTCTCCT GCTGACTCAT

1381	CCCACCATTT CTGACGCTGC AGTTGTTCCT ATGATCGACG AAAAGGCTGG AGAAGTGCCC

1441	GTTGCATTTG TGGTTAGAAC CAACGGATTT ACAACGACCG AGGAAGAGAT CAAGCAATTC

1501	GTGTCTAAGC AAGTCGTCTT TTACAAACGT ATTTTTCGGG TCTTCTTTGT GGACGCTATT

1561	CCCAAGTCCC CCTCCGGTAA GATTCTTCGA AAGGATCTTA GAGCTAAGAT CGCCTCTGGA

1621	GACCTTCCTA AATAA

Bbxfpk sequences:

1	ATGACATCTC CCGTCATCGG CACGCCTTGG AAGAAACTCA ATGCTCCTGT GTCCGAGGAA

61	TCTCTTGAGG GAGTGGATAA ATACTGGAGA GTGGCTAACT ACCTCTCTAT TGGTCAGATC

121	TACCTTCGTT CGAATCCCCT CATGAAGGCA CCTTTTACAC GAGAGGATGT GAAACATCGG

181	CTCGTCGGAC ATTGGGGTAC TACACCCGGC CTTAATTTTC TTATCGGTCA TATTAACAGA

241	TTCATTGCAG ACCATGGACA GAACACAGTT ATCATCATGG GTCCTGGCCA TGGTGGCCCC

301	GCAGGAACGT CCCAGAGCTA TCTTGATGGA ACGTATACAG AGACCTTTCC CAAAATTACG

361	AAGGACGAAG CTGGCCTCCA GAAGTTTTTT CGGCAGTTCT CGTATCCCGG AGGAATCCCT

421	AGCCACTTCG CACCTGAGAC GCCCGGTTCT ATCCACGAAG GTGGAGAACT TGGTTACGCT

481	CTCTCGCATG CCTACGGTGC TATTATGGAT AATCCCTCGC TTTTTGTCCC TGCCATTGTT

541	GGAGATGGAG AAGCAGAAAC GGGACCTCTT GCTACTGGTT GGCAGTCTAA CAAGCTTGTG

601	AATCCTCGGA CAGATGGAAT CGTGCTCCCT ATCCTTCATC TGAATGGTTA TAAGATCGCT

661	AACCCTACTA TTCTGTCTCG TATCTCGGAT GAAGAGCTCC ATGAGTTTTT CCACGGAATG

721	GGCTACGAGC CTTATGAATT CGTGGCTGGA TTTGACGACG AGGACCATAT GAGCATCCAC

781	CGTCGATTCG CCGAACTCTG GGAGACCATT TGGGACGAGA TCTGCGACAT TAAGGCAGCT

841	GCACAAACGG ATAATGTTCA CCGACCTTTC TACCCCATGC TCATCTTCCG AACGCCCAAG

901	GGCTGGACTT GTCCCAAGTA TATTGACGGA AAAAAGACCG AAGGTTCTTG GAGAGCTCAC

961	CAGGTCCCCC TTGCTTCCGC TCGTGATACG GAAGCACACT TTGAGGTGCT GAAGAACTGG

1021	CTTGAATCCT ATAAGCCCGA GGAGCTTTTT GATGCAAACG GCGCCGTCAA GGATGACGTC

1081	CTTGCATTTA TGCCTAAAGG AGAGCTCCGA ATCGGCGCCA ACCCCAATGC CAACGGTGGC

1141	GTTATCCGAG ATGATCTGAA ACTGCCTAAC CTGGAAGACT ATGAGGTCAA AGAGGTTGCT

1201	GAATATGGTC ATGGATGGGG ACAGCTTGAG GCCACCCGAA CGCTCGGTGC ATATACCCGT

1261	GACATCATTC GAAACAACCC TCGAGATTTT CGAATTTTCG GACCCGACGA AACAGCATCC

1321	AACCGACTGC AGGCCTCCTA TGAGGTGACG AACAAACAAT GGGATGCCGG ATACATTTCG

1381	GACGAAGTTG ACGAGCACAT GCACGTGTCC GGACAGGTTG TCGAGCAGCT GTCGGAACAT

1441	CAAATGGAAG GATTCCTCGA GGCCTATCTG CTCACCGGAA GACACGGAAT CTGGAGCTCC

1501	TACGAGAGCT TTGTCCACGT TATTGATTCG ATGCTGAATC AACATGCTAA ATGGCTCGAA

1561	GCAACGGTCA GAGAGATTCC CTGGCGAAAG CCCATTGCAT CGATGAATCT TCTGGTGTCC

1621	TCGCACGTCT GGCGACAGGA TCACAACGGT TTCTCCCATC AAGACCCTGG AGTGACCTCG

1681	GTCCTGCTCA ATAAGTGCTT CCATAACGAC CATGTTATCG GTATCTACTT TGCAACCGAC

1741	GCAAATATGC TCCTCGCAAT CGCAGAGAAA TGTTACAAAA GCACGAATAA AATTAATGCA

1801	ATCATTGCCG GCAAGCAGCC TGCCGCAACG TGGCTGACTC TTGATGAGGC ACGAGCCGAA

1861	CTTGCTAAAG GCGCAGCCGC ATGGGATTGG GCTTCCACGG CCAAGAACAA CGATGAGGCC

1921	GAGGTCGTCC TCGCAGCTGC AGGCGACGTG CCTACGCAGG AGATCATGGC CGCCTCTGAT

1981	AAGCTGAAAG AGCTCGGAGT CAAGTTTAAA GTTGTTAATG TTGCTGATCT CCTGTCCCTG

2041	CAGAGCGCCA AGGAAAATGA TGAAGCCCTC AGCGACGAAG AATTCGCAGA TATCTTCACC

2101	GCTGATAAGC CTGTGCTTTT CGCTTATCAC AGCTACGCCC ACGACGTGCG AGGTCTGATT

2161	TACGACCGAC CCAATCATGA CAACTTTAAT GTCCATGGCT ATGAAGAAGA AGGTTCGACC

2221	ACTACCCCTT ATGATATGGT CCGGGTCAAT CGGATCGATC GATATGAGCT CACGGCAGAA

2281	GCTCTGAGAA TGATTGACGC AGACAAGTAT GCTGATAAAA TCGACGAGCT CGAGAAGTTT

2341	CGGGATGAGG CCTTCCAGTT TGCAGTCGAT AAGGGCTATG ATCATCCCGA TTATACCGAT

2401	TGGGTTTACT CTGGCGTCAA CACGGATAAA AAAGGCGCCG TCACCGCAAC CGCCGCAACA

2461	GCAGGCGACA ATGAGTAA

AcxpkA sequences:

1	ATGTCAAAAA CGGCGACAAA TGCGGAACCT ACACTGAAAC CGCAAGAACT TCAGCGCATG

61	GACGCGTATT GGCGCGCTTG TAATTATTTA GCGGCGGGCA TGATCTATCT GCGCGAGAAC

121	CCGCTTCTGA AAGAGCCGCT GCGCCCGGAG CATATTAAAA ACCGTTTACT GGGACATTGG

181	GGAAGCGACC CGGGCCAGTC TTTTGTGTGG GTGCATCTGA ACCGTTTAAT TAGAGAGCAA

241	GATTTAAACA TGATCTACAT CAGCGGACCC GGTCACGGCG CACCCGCTAC GCTTGCGAAC

301	TGCTACCTTG AAGGCACGTA CTCAGAGATC TACCCCGATA AAAGCCATGA CGTGGAGGGT

361	ATGCGTAAGT TCTTCCGCCA ATTTAGCTTT CCGGGCGGCA TCGGATCACA TTGCACGCCG

421	GAAACACCGG GCTCAATCCA TGAAGGCGGA GAACTGGGAT ACTCACTTAG CCACGCATTT

481	GGCGCGGCGT TTGATCATCC GGATCTGATC GTGAACGTGG TGGTTGGCGA CGGAGAAGCA

541	GAAACTGGTC CGATGGCAAC AAGCTGGCAC GCAAATAAGT TTTTAAACCC CGCTCGCGAT

601	GGAGCGGTGC TGCCGATTCT GCATCTTAAC GGCTATAAGA TCGCGAACCC GACGATCCTT

661	GCGCGCATCT CACATGAAGA GCTGGAAGCG CTGTTTACTG GTTACGGATG GAAGCCGTAT

721	TTCGTGGAAG GAAGCGAACC GGAACAGATG CATCAGAAAA TGGCGGCGAC ACTGGACAGC

781	TGTGTGCGCG AAATTAAGGA GATCCAAGAA CAAGCGCGCG AAAGCGGCAA ATGGGAGCGC

841	CCTCGCTGGC CTATGATTGT GCTGCGCTCA CCGAAAGGCT GGACTGGTCC GAAAGAAGTG

901	GATGGACACA AGGTGGAGGA TTTCTGGAGA GCGCATCAAG TGCCGATTTT AGGCGTTAAG

961	GAAAACCCGG AACATCTGAA AATGCTGGAG GCGTGGATGC GCTCATACGA GCCGGAAAAA

1021	CTGTTTGATG AGAGCGGACG TTTAGTTGCA GAACTGCAAG AACTTGCACC GAAGGGCGAC

1081	CGCAGAATGA GCGCGAATCC TCATACGAAC GGCGGCAAAT TACGCAAGCC GCTGGATCTG

1141	CCGGCGTTTT GTGACTTCGC ACTTAAATTT CAGAGACCGG GCGAGATGTA CGCATCATCA

1201	ACAGAGACGC TGGGAACGTA TCTGGCAGAG GTGTTTAGAC GCAATCCGGA GAGCTTTCGT

1261	TTATTCGGCC CGGATGAAAC GGCAAGCAAC AAACTGAGCG GCGTGTACGA AGCGACAAAG

1321	AAAACGTGGG AAGCTGGTTA CAAGCCGGAG GATGCGGATG GCGGCGAACT TGCGGCGGAT

1381	GGCAGAGTTA TGGAGATGCT GAGCGAACAT ACGCTGGAAG GCTGGCTTGA GGGCTATCTG

1441	CTGACTGGTC GCCATGGCTT CTTTGCGACG TATGAAGCGT TCGTGCACGT GATTGATAGC

1501	ATGTTCAATC AGCACGCGAA GTGGCTGGAA AAGTCAAAGA AGGAGATCCG CTGGCGCGCA

1561	CCGATCAGCT CTTTAAATCT GCTGATCACA AGCGTGGTTT GGAGACAAGA TCACAACGGC

1621	TTTACGCACC AAGATCCCGG TTTTCTGGAC ATCGTGGCAA ATAAGAGCGC AGAGGTGACG

1681	CGCATTTATC TGCCGCCGGA TGCGAATTGT TTACTGTCAG TGGCGGACCA TTGTTTAAAG

1741	AGCACAGATT ATGTGAACGT GATCGTTGCG GACAAACAGC CTCATCTTCA ATTTCTGAGC

1801	GCGGACGAGG CAATCAAACA CTGCACAAAG GGCATCGGCA TCTGGGAGTG GGCATCAACA

1861	GACAAGGGCT GTGAACCGGA CGTGGTGATC GCATCAGCTG GTGATATTGC GACAATGGAG

1921	GCACTTGCAG CAGCGGCGCT GCTTCGCGAA CATTTTCCTA AATTAAAAAT CAGATTTGTG

1981	AATGTTGTTG ATTTATTCAG ACTGGTTCCG GAAGATGAAC ATCCTCACGG ACTGCCGGAG

2041	AGAGACTATG ATTCTTTATT TCCTCCGGAC ACGCCGGTGA TTTTTAACTT CCATGGCTAT

2101	CCTCAGCTGA TCCACCGCCT TACGTATCAA CGCAACAACC ACCACAACCT TCACGTGCAT

2161	GGATATCGCG AGAGAGGCAA CATCAACACG CCTCTTGAGC TGGCGATTAT GAACAAAGTG

2221	GATCGCTTTC ATCTTGCGAT GAACGCGATT GATCGCGTTC CCGGTCTTAG AGCAATTGGC

2281	GGACATGCGA AAAACTGGCT GTTCGACCAA GTTACAGAGC ACGTTATGTA CGCGCACGAG

2341	CACGGCATTG ATCCGGAAGC GATCAATGAA TGGACGTGGC CGGAATAA

### **References**

1.	Madzak, C., Treton, B., Blanchin-Roland, S. (2000) Strong hybrid promoters and integrative expression/secretion vectors for quasi-constitutive expression of heterologous proteins in the yeast *Yarrowia lipolytica*. *J. Mol. Microbiol. Biotechnol.* *2* (2), 207-16.

2.	Gu, Y., Ma, J., Zhu, Y., Xu, P. (2020) Refactoring Ehrlich pathway for high-yield 2-phenylethanol production in *Yarrowia lipolytica*. *ACS Synth. Biol.* *9* (3), 623-633.

3.	Wong, L., Engel, J., Jin, E., Holdridge, B., Xu, P. (2017 YaliBricks, a versatile genetic toolkit for streamlined and rapid pathway engineering in *Yarrowia lipolytica*. *Metab. Eng. Commun.* *5*, 68-77.

4.	Lv, Y., Edwards, H., Zhou, J., Xu, P. (2019) Combining 26s rDNA and the Cre-loxP System for Iterative Gene Integration and Efficient Marker Curation in *Yarrowia lipolytica*. *ACS Synth. Biol.* *8* (3), 568-576.

5.	Lv, Y., Marsafari, M., Koffas, M., Zhou, J., Xu, P. (2019) Optimizing oleaginous yeast cell factories for flavonoids and hydroxylated flavonoids biosynthesis. *ACS Synth. Biol.* *8* (11), 2514-2523.

6.	Shen, L., Nishimura, Y., Matsuda, F., Ishii, J., Kondo, A. (2016) Overexpressing enzymes of the Ehrlich pathway and deleting genes of the competing pathway in *Saccharomyces cerevisiae* for increasing 2-phenylethanol production from glucose. *J. Biosci. Bioeng.* *122* (1), 34-39.

7.	Romagnoli, G., Luttik, M. A., Kötter, P., Pronk, J. T., Daran, J.-M. (2012) Substrate specificity of thiamine pyrophosphate-dependent 2-oxo-acid decarboxylases in *Saccharomyces cerevisiae*. *Appl. Environ. Microbiol. 78* (21), 7538-7548.

8.	Hassing, E. J., de Groot, P. A., Marquenie, V. R., Pronk, J. T., Daran, J. G. (2019) Connecting central carbon and aromatic amino acid metabolisms to improve de novo 2-phenylethanol production in *Saccharomyces cerevisiae*. *Metab. Eng.* *56*, 165-180.

9.	Guo, D., Zhang, L., Kong, S., Liu, Z., Li, X., Pan, H. (2018) Metabolic engineering of *Escherichia coli* for production of 2-phenylethanol and 2-phenylethyl acetate from glucose. *J. Agric. Food Chem.* *66* (23), 5886-5891.

10.	Machas, M. S., McKenna, R., Nielsen, D. R. (2017) Expanding upon styrene biosynthesis to engineer a novel route to 2‐phenylethanol. *Biotechnol. J.* *12* (10), 1700310.

11.	Guo, D., Zhang, L., Pan, H., Li, X. (2017) Metabolic engineering of *Escherichia coli* for production of 2-Phenylethylacetate from L-phenylalanine. *MicrobiologyOpen 6* (4), e00486.

12.	Zhang, H., Cao, M., Jiang, X., Zou, H., Wang, C., Xu, X., Xian, M. (2014) De-novo synthesis of 2-phenylethanol by *Enterobacter sp.* CGMCC 5087. *BMC Biotechnol. 14* (1), 30.

13.	Kang, Z., Zhang, C., Du, G., Chen, J. (2014) Metabolic engineering of *Escherichia coli* for production of 2-phenylethanol from renewable glucose. *Appl. Biochem. Biotechnol.* *172* (4), 2012-2021.

14.	Koma, D., Yamanaka, H., Moriyoshi, K., Ohmoto, T., Sakai, K. (2012) Production of aromatic compounds by metabolically engineered *Escherichia coli* with an expanded shikimate pathway. *Appl. Environ. Microbiol. 78* (17), 6203-6216.

15.	Liu, Q., Yu, T., Li, X., Chen, Y., Campbell, K., Nielsen, J., Chen, Y. (2019) Rewiring carbon metabolism in yeast for high level production of aromatic chemicals. *Nat. Commun.* *10* (1), 4976.

16.	Kang, S.-Y., Choi, O., Lee, J. K., Hwang, B. Y., Uhm, T.-B., Hong, Y.-S. (2012) Artificial biosynthesis of phenylpropanoic acids in a tyrosine overproducing *Escherichia coli* strain. *Microb. Cell Fact.* *11* (1), 153.

17.	Shin, S. Y., Han, N. S., Park, Y. C., Kim, M. D., Seo, J. H. (2011) Production of resveratrol from *p-*coumaric acid in recombinant *Saccharomyces cerevisiae* expressing 4-coumarate:coenzyme A ligase and stilbene synthase genes. *Enzyme Microb. Technol.* *48* (1), 48-53.

18.	Rodriguez, A., Kildegaard, K. R., Li, M., Borodina, I., Nielsen, J. (2015) Establishment of a yeast platform strain for production of p-coumaric acid through metabolic engineering of aromatic amino acid biosynthesis. *Metab. Eng.* *31*, 181-8.

19.	Kanelli, M., Mandic, M., Kalakona, M., Vasilakos, S., Kekos, D., Nikodinovic-Runic, J., Topakas, E. (2018) Microbial production of violacein and process optimization for dyeing polyamide fabrics with acquired antimicrobial properties. *Front. Microbiol. 9*, 1495.

20.	Ahmad, W. A., Yusof, N. Z., Nordin, N., Zakaria, Z. A., Rezali, M. F. (2012) Production and characterization of violacein by locally isolated Chromobacterium violaceum grown in agricultural wastes. *Appl. Biochem. Bbiotechnol.167* (5), 1220-1234.

21.	Tong, Y., Zhou, J., Zhang, L., Xu, P. (2019) Engineering oleaginous yeast *Yarrowia lipolytica* for violacein production: extraction, quantitative measurement and culture optimization. *bioRxiv* 687012.
